# Parallel genetic changes underlie integrated craniofacial traits in an adaptive radiation of trophic specialist pupfishes

**DOI:** 10.1101/2021.07.01.450661

**Authors:** Michelle E. St. John, Julia C. Dunker, Emilie J. Richards, Stephanie Romero, Christopher H. Martin

## Abstract

Many factors such as divergence time, shared standing genetic variation, frequency of introgression, and mutation rates can influence the likelihood of whether populations adapt to similar environments via parallel or non-parallel genetic changes. However, the frequency of parallel *vs* non-parallel genetic changes resulting in parallel phenotypic evolution is still unknown. In this study, we used a QTL mapping approach to investigate the genetic basis of highly divergent craniofacial traits between scale- and snail-eating trophic specialist species across similar hypersaline lake environments in an adaptive radiation of pupfishes endemic to San Salvador Island, Bahamas. We raised F2 intercrosses of scale- and snail-eaters from two different lake populations of sympatric specialists, estimated linkage maps, scanned for significant QTL for 30 skeletal and craniofacial traits, and compared the location of QTL between lakes to quantify parallel and non-parallel genetic changes. We found strong support for parallel genetic changes in both lakes for five traits in which we detected a significant QTL in at least one lake. However, many of these shared QTL affected different, but highly correlated craniofacial traits in each lake, suggesting that pleiotropy and trait integration should not be neglected when estimating rates of parallel evolution. We further observed a 23-52% increase in adaptive introgression within shared QTL, suggesting that introgression may be important for parallel evolution. Overall, our results suggest that the same genomic regions contribute to parallel integrated craniofacial phenotypes across lakes. We also highlight the need for more expansive searches for shared QTL when testing for parallel evolution.

## Introduction

Convergent evolution describes the independent evolution of similar phenotypes in response to similar selective pressures and provides strong support for ecological adaptation (Losos 2009; Schluter 2000). This includes both non-parallel genetic changes, such as the evolution of antifreeze glycoproteins in icefishes or the ‘thunniform’ body shape of lamnid sharks and tunas (Chen et al. 1997; Donley et al. 2004), and parallel genetic changes such as tetrodotoxin resistance in snakes and pufferfishes or the evolution of voltage-gated sodium channels in mormyrid and gymnotiform electric fishes (Hopkins 1995; Katz 2006; Jost et al. 2008; Feldman et al. 2009; Zakon et al. 2006). Instances of convergence across independent lineages (i.e., across groups that lack a recent common ancestor and shared genetic backgrounds) provide the strongest evidence for adaptation; however, repeated evolution of similar phenotypes in response to similar selective pressures among lineages derived from the same ancestral population can also provide insight into the process of adaptation. Understanding this process, traditionally known as parallel evolution (Futuyman 1986), is important because it can help to tease apart the contributions of natural selection and shared genetic constraints to similar phenotypes (Schluter et al. 2004; Stuart et al. 2017). Parallel phenotypic evolution can also occur via parallel or non-parallel genetic changes (e.g., Cresko et al. 2004), but even non-parallel genetic changes occurring in the same ancestral genetic background (e.g. Chan et al. 2010; Xie et al. 2019) provide weaker evidence for adaptation than convergence across independent lineages due this shared history. Despite substantial attention, the frequency and likelihood of parallel phenotypic evolution via parallel or non-parallel genetic changes is still relatively unknown (Stern and Orgogozo 2008; Stern 2013; Rosenblum et al. 2014).

Many factors influence whether parallel phenotypic evolution in similar environments is produced by parallel or non-parallel genetic mechanisms. First, recently diverged species exhibit increased probabilities of genetic parallelism when adapting to similar environments. Recently diverged taxa may inhabit similar environments more frequently or they may have similar genetic architectures, similar genetic variance-covariance matrices, or similar genetic backgrounds that produce similar epistatic interactions (Conte et al. 2012; Rosenblum et al. 2014). Second, any mechanism that allows the use of the same adaptive genetic mechanism should increase the likelihood of convergence via parallelism, including the availability of shared standing genetic variation and introgression (Rosenblum et al. 2014). For example, threespine sticklebacks colonized freshwater thousands of times and converged on similar phenotypes largely due to selection on an ancient shared pool of marine standing genetic variation (Jones et al. 2012; Feulner et al. 2013; Nelson and Cresko 2018; Haenel et al. 2019; but see: Chan et al. 2010; Stuart et al. 2017). Similarly, increased adaptive introgression should also make genetic parallelism more likely (Grant et al. 2004; Morjan and Rieseberg 2004; Hedrick 2013; Taylor et al. 2020). Third, adaptive genetic variation with larger effect sizes and fewer pleiotropic effects should be reused more frequently across populations, particularly when a population is far from a new adaptive optimum (Linnen et al., 2013; Orr, 2005; Stern, 2013). Finally, de novo mutations, large mutational target sizes, and polygenic adaptive phenotypes are more likely to result in parallel phenotypic evolution via non-parallel genetic pathways (Wittkopp et al. 2003; Kowalko et al. 2013; Bolnick et al. 2018, but see: Colosimo et al. 2004; Chan et al. 2010; Xie et al. 2019).

Quantitative trait locus (QTL) mapping is often used to infer whether parallel or non-parallel genetic changes underlie parallel phenotypes. However, many QTL studies only investigate a limited number of traits that are controlled by large effect loci, which may bias the literature towards supporting genetic parallelism (Conte et al. 2012). This bias may be exacerbated by the fact that in many QTL studies the genomic regions associated with a parallel phenotype are large, contain many genes, and their effects on phenotypic variance are overestimated in under-powered studies (Beavis 1998). These methodological and experimental limitations reduce confidence in the specific genomic regions associated with a parallel phenotype and, by extension, reduce confidence in whether parallel evolution was due to parallel or non-parallel genetic changes. One possible solution is to compare the genomic regions associated with many different phenotypes across populations (Erickson et al. 2016). In this scenario, shared genomic regions across populations provide strong support for genetic parallelism, except in the likely rare instances of independent de novo mutations within the same region (O’Brown et al. 2015; Xie et al. 2019; Chan et al. 2010).

The San Salvador Island (SSI), Bahamas pupfish radiation is an excellent system for investigating the genetic underpinnings of parallel ecomorph phenotypes because novel trophic specialists occur in sympatry across multiple hypersaline lake populations on the island. The radiation includes three pupfish species: a generalist pupfish (*Cyprinodon variegatus*), a scale-eating (lepidophagous) pupfish (*C. desquamator*), and a snail-eating (durophagous) pupfish (*C. brontotheroides;* Martin and Wainwright 2013). The snail- and scale-eating pupfishes are endemic to SSI and occur in sympatry with one another and the generalist pupfish.

Among lakes, specialists have converged on multivariate phenotypes that are adaptive for their given ecological niche. For example, scale-eaters across all lakes exhibit increased oral jaw size (Martin & Wainwright, 2013; Hernandez et al. 2018) and reduced lower jaw angles during scale-eating strikes which may play a critical role in scale-biting performance during high-speed strikes on their prey (St. John et al. 2020b). Similarly, the snail-eating pupfish exhibits a novel nasal protrusion which may improve oral snail-shelling performance or result from sexual selection (Martin and Wainwright 2013; St. John et al. 2020a). Furthermore, the nasal protrusion of the snail-eating species varies substantially among lake populations (Martin and Feinstein 2014; Hernandez et al. 2018). Despite the importance of these species characteristics, we still do not understand how their genetic architecture varies across populations.

There is some evidence to suggest that parallel genetic changes underlie specialist phenotypes on SSI. First, the SSI radiation is very young, diverging only about 10 kya (Hagey and Mylroie 1995). Second, previous genomic analyses show that many of the alleles associated with trophic specialization arrived on SSI from Caribbean-wide standing genetic variation within generalist pupfish populations, but there are also some de novo adaptive mutations associated with scale-eating (Richards et al. 2021). Scale-eaters form a monophyletic group, suggesting a shared genetic component to the scale-eating phenotype across lakes (Richards and Martin 2017). In contrast, snail-eaters and generalists often genetically cluster together by lake instead of by species—suggesting that non-parallel genetic changes could underlie parallel snail-eater phenotypes across lakes (Martin and Feinstein 2014; Richards and Martin 2017). Furthermore, previous studies have documented strong genetic divergence between scale-eaters from Crescent Pond and all other populations of scale-eater (Richards & Martin, 2017; Richards et al., 2021).

In this study we mapped the genetic basis of 30 skeletal craniofacial and body traits associated with snail- and scale-eating using lab-reared F2 intercrosses from Crescent Pond and Little Lake. We called variants, estimated linkage maps, and performed QTL analyses independently for each F2 population. We found that only one trait— cranial height—mapped to the same genomic region in both Crescent Pond and Little Lake, but 4 of the 5 remaining significant QTL detected in one lake mapped to the same genomic region as a highly correlated craniofacial trait in the second lake. Ultimately, we conclude that parallel evolution through reuse of introgressed adaptive alleles is acting to produce similar snail- and scale-eating phenotypes across lake populations on SSI.

## Methods

### Genetic cross

Currently, pupfish species can been found in 12 hypersaline lakes across the island: generalist pupfish are found in allopatry in five lakes, the generalist and snail-eating pupfish are found in sympatry without the scale-eater in two lakes, the generalist and scale-eater are found in sympatry in a single lake, and all three species are found in sympatry in four lakes (Martin and Feinstein 2014). We collected wild-caught scale-eating and snail-eating pupfishes from two different sympatric populations (containing all three species) on SSI – Crescent Pond and Little Lake—during the years of 2011 and 2013-2015 using seine nets. We brought individuals back to the University of California, Davis or the University of California, Berkeley and a single wild-caught scale-eating female from each lake was allowed to breed freely with a single wild-caught snail-eating male from the same lake resulting in two separate genetic crosses (one cross from Crescent Pond and one cross from Little Lake). At least four F1 offspring from each hybrid population were crossed to produce F2 intercrosses, resulting in 354 individuals from Crescent Pond and 287 individuals from Little Lake included in this study. All fish were maintained in 40-L tanks at 5-10ppt salinity at the University of California, Davis or the University of California, Berkeley. We fed fry a diet of newly hatched *Artemia* nauplii for approximately 40 days post fertilization, after which they were switched to the adult diet of frozen and pellet foods. We euthanized fish in an overdose of MS-222 (Finquel, Inc.) according to the approved University of California, Davis Institutional Animal Care and Use Protocol #17455 or University of California, Berkeley IACUC Protocol AUP-2015-01-7053, and stored them in 95% ethanol.

### Phenotyping

#### Sex and Mate Preferences

For individuals from Crescent Pond, we recorded their sex using their sexually dimorphic body and fin coloration. Male pupfish develop a blue iridescent coloration along their anteriodorsal surface and a black marginal band along their caudal fin (Echelle and Echelle 2020).

Once F2 hybrids reached sexual maturity, we performed mating assays using a subset of the hybrid females from Crescent Pond to estimate mate preferences for snail- or scale-eating mates (N=74). Prior to the mating assays, female fish were isolated for at least twelve hours and conditioned on frozen bloodworms with a 12:12 light:dark cycle. Mating assays occurred in three m diameter kiddie pools (5-10ppts salinity). Pools were covered with gravel substrate and divided in half. In each half, we placed three clear plastic 7.5-L Kritter Keepers in a row containing three conspecific males housed individually to avoid aggression. Size-matched scale-eater males were placed on one side of each arena and snail-eating males on the other. Once the males were placed in individually in clear boxes, a female F2 hybrid from Crescent Pond was placed into the center of one of the three kiddie pools, chosen at random. We considered females acclimated to the pool once they had visited both rows of males, after which we started the seven-minute trial period. During each trial we recorded the amount of time a female spent within one body-length of each species. Each female was tested consecutively in all three pools, and we used the mean of her association time (scale-eater association time / total association time during each 7-minute trial) across all three pools for QTL analysis. Size-matched males were periodically rotated into and among kiddie pools during the 12-month testing period.

#### Morphological Traits

To measure skeletal phenotypes in our F2 intercrosses, we cleared and double-stained each specimen with alizarin red and alcian blue. Before clearing and staining, each fish was skinned and fixed in 95% ethanol. We then fixed specimens in 10% buffered formalin for at least one week and stained batches of individually labeled specimens following Dingerkus and Uhler’s (1977) protocol. We suspended cleared and stained specimens in glycerin, and photographed their left lateral side using a Canon EOS 60D digital SLR camera with a 60 mm macro lens. For each individual, we took two types of photographs: first, we took a whole-body photograph to calculate fin and body measurements and second, a lateral skull image to calculate craniofacial measurements (Figure 1). We used DLTdv8 software (Hedrick 2008) to digitize 11 landmarks on each whole body image and 19 landmarks on each lateral skull image following the morphometric methods described in Martin et al. (2017). For individuals from Crescent Pond, we also weighed the adductor mandibulae muscle mass. Each image included a standardized grid background which we used to calibrate and transform our measurements from pixels into millimeters. In total, we measured 354 individuals from Crescent Pond and 287 individuals from Little Lake. We used R to convert the 30 landmarks into linear distances. To reduce measurement error due to the lateral positioning of the specimens, we took the mean distances from the two clearest skull and whole-body photographs for each individual when possible. If an individual did not have two clear photographs for each orientation, we measured the single clearest photograph. Finally, we size-corrected each trait by using the residuals from a linear model including the log-transformed measurement of each trait as the response variable and log-transformed standard length as the predictor variable. We investigated whether size-corrected traits varied between the two populations using a PCA and a MANOVA test, but found no appreciable difference between them (Fig. S1, num df = 28, approximate *F*-value= 0.34, *P* = 1)

**Figure 1.**
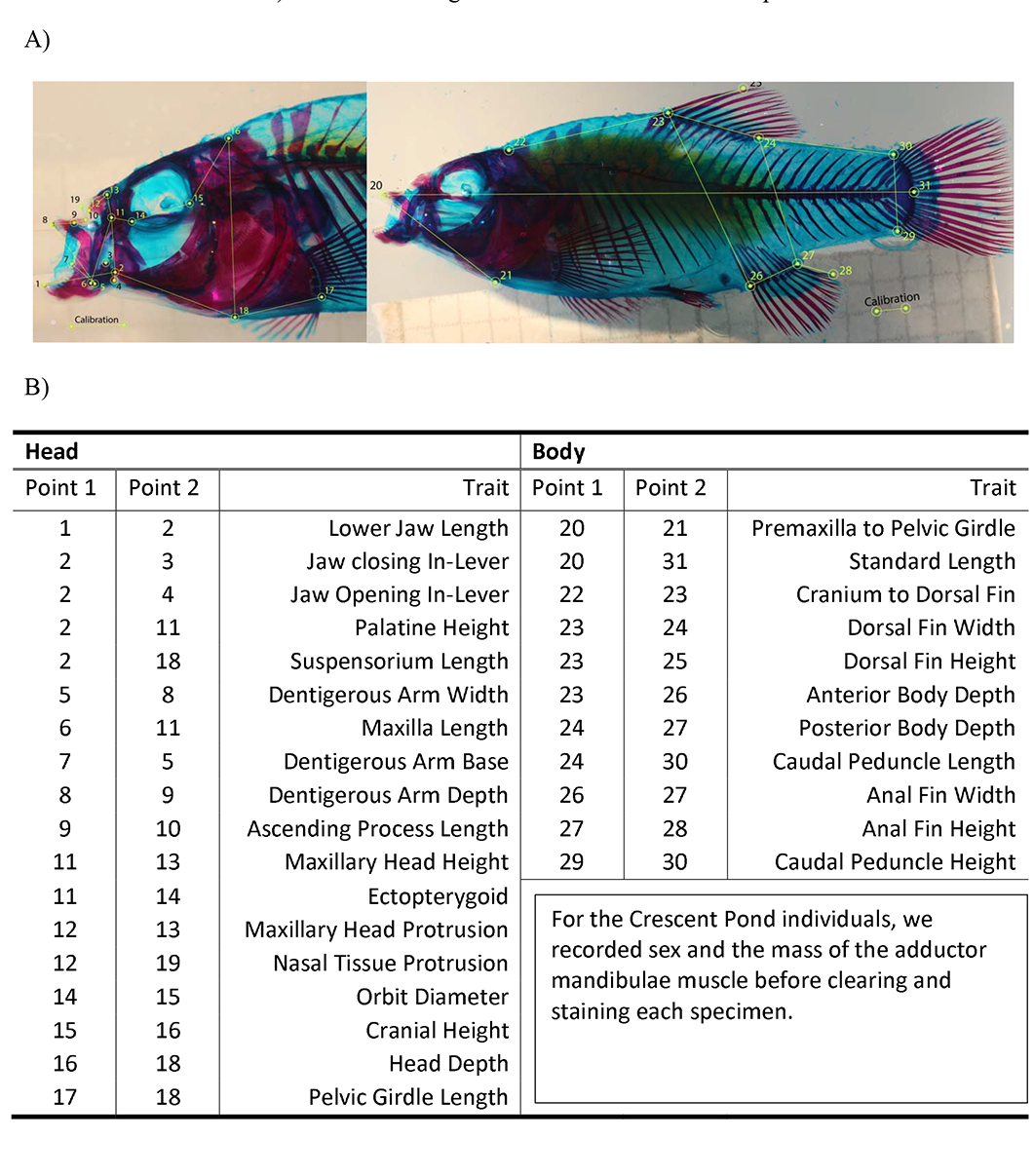
A) Representative photographs of F2 intercross cleared and double-stained specimen used for skeletal morphometrics. Points represent landmarks used to measure linear distances between skeletal traits. B) Table containing the two landmarks that correspond to each trait.

#### Genotyping

We genotyped individuals using three different methods: First, we used whole genome resequencing for the wild-caught F0 parental generation of our Crescent Pond and Little Lake intercrosses. We used DNeasy Blood and Tissue Kits (Qiagen, Inc.) to extract DNA from the muscle tissue of each fish and quantified it on a Qubit 3.0 fluoromether (Thermofisher Scientific, Inc.). Genomic libraries were then prepared at the Vincent J. Coates Genomic Sequencing Center (QB3) using the automated Apollo 324 system (WaterGen Biosystems, Inc.). Samples were fragmented using Covaris sonication and barcoded with Illumina indices. A quality check was also performed on all samples using a Fragment Analyzer (Advanced Analytical Technologies, Inc.). We used 150 paired-end sequencing on an Illumina Hiseq4000 for these four parental samples along with an additional 38 samples that were included in a previous study (Richards and Martin 2017).

Second, in addition to the 190 previously sequenced individuals from Crescent Pond used for a QTL mapping study (Martin et al. 2017), we included an additional 164 F2 individuals from Crescent Pond sequenced using double-digest restriction site associated sequencing (ddRADseq) following similar library prep and sequencing methods described in Martin et al. (2015, 2016, 2017). Briefly, we prepared four indexed libraries each containing 96 barcoded individuals. We sequenced these using 100 single-end high-output mode on two lanes of Illumina Hiseq4000 at the Vincent J. Coates Genomic Sequencing Center (QB3).

Finally, we sequenced all F2 individuals from Little Lake and a subset of previously sequenced, but low-coverage Crescent Pond F2’s (N=84), using Nextera-tagmented reductively-amplified DNA (NextRad) sequencing (Russello et al. 2015). We followed the above methods for DNA extraction and sent samples to SNPsaurus (SNPsaurus, LLC) for quality checking, NextRad library preparation, and 150 single-end sequencing on two lanes of Illumina Hiseq4000 at the University of Oregon sequencing core.

#### Calling Variants

We used the following methods to call variants separately for: 1) the Crescent Pond intercross (2 parents and 354 F2 hybrids), and 2) the Little Lake intercross (2 parents and 285 F2 hybrids): First, we inspected raw read quality using FastQC (Babraham Bioinformatics Institute, v0.11.7) and trimmed reads to their appropriate length (100bp for samples sequenced with ddRAD, and 150bp for samples sequenced with NextRAD) using TrimGalore! (v0.6.4). For samples that were sequenced using both ddRAD and NextRad methods, we concatenated trimmed raw reads into a single file. We next used bwa-mem to map reads from all individuals in an intercross, both parents and offspring, to the *Cyprinodon brontotheroides* reference genome (v 1.0; total sequence length = 1,162,855,435 bp; number of scaffolds = 15,698, scaffold N50 = 32 Mbp; (Richards et al. 2021)). We identified duplicate reads using MarkDuplicates and created BAM indices using BuildBamIndex in the Picard package (http://picard.sourceforge.net(v.2.0.1)). Following the best practices guide from the Genome Analysis Toolkit (v 3.5; (Depristo et al. 2011)), we called and refined our single nucleotide polymorphism (SNP) variant data set using the program HaplotypeCaller. Pupfish lack high-quality known alleles because they are a non-model organism; we therefore used the recommended hard filter criteria (QD < 2.0; FS <; 60; MQRankSum < -12.5; ReadPosRankSum < -8; (Depristo et al. 2011; Marsden et al. 2014)) to filter our SNP variant dataset. Ultimately, we detected 13.7 million variants in our Crescent Pond dataset and 14.4 million variants in our Little Lake dataset.

We used the program STACKS to further filter our dataset and convert our vcf files into phenotype and genotype comma-separated values files that could be imported into the Rqtl program. Specifically, we used the populations program to filter out variants that were not present in both the parental and F2 populations, and to filter out variants found in 10% or less of the population. From this filtering step we retained 36,318 variants with 46.5 mean mappable progeny per site in Crescent Pond and 87,579 variants with 85.984 mean mappable progeny per site in Little Lake.

We continued to filter our datasets using the Rqtl (v1.46-2), and ASMap (v1.0-4) packages (Broman et al. 2003; Taylor and Butler 2017). We started filtering by removing individuals that did not contain any filtered variants and any duplicate individuals. This reduced our Crescent Pond data set to 227 individuals, and our Little Lake data set to 281 individuals. Next, we filtered markers that had >0.98 or <0.1 heterozygosity (Crescent Pond: markers =15,247, Little Lake: markers=14,661). This step also filtered out 13 individuals from Crescent Pond which only contained markers with >0.98 or <0.1 heterozygosity. Before constructing our genetic maps, we set aside markers that appeared to suffer from segregation distortion. We used the pullCross() function from the ASmap package to set aside markers in both data sets that were missing in >75% of individuals, departed from Mendelian ratios (1:2:1), or any co-located markers for the initial construction of the linkage maps. This filtering retained more than twice the number of markers for Crescent Pond than Little Lake. We therefore used a stricter filtering threshold for missing data (i.e., removing markers with >72% missing data) for our Crescent Pond dataset to construct linkage maps of comparable sizes for downstream comparative analyses. At the end of this filtering process the Crescent Pond dataset contained 214 individuals and 657 markers and the Little Lake dataset contained 281 individuals with 490 markers.

#### Linkage Map Construction

We used the mstmap.cross() function to form initial linkage groups and order markers, using the kosambi method for calculating genetic distances and a clustering threshold of *P* = 1 x10^-14^ for Little Lake and *P* = 1 x 10^-20^ for Crescent Pond. After forming these initial linkage groups, we used the pushCross() function from the ASmap package to integrate previously set aside markers back into our map. We pushed markers back based on a segregation ratio of 3:4:3 and we pushed back any markers that had previously been designated as co-located. This increased our map sizes to 817 markers for Crescent Pond and 580 markers for Little Lake. With these additional markers, we re-estimated our linkage map using the est.rf() and formLinkageGroups() functions from the Rqtl package. We used a max recombination fraction of 0.35 and a minimum LOD threshold of 5 to estimate linkage groups for both data sets. We used the droponemarker() command from Rqtl with an error probability of 0.01 to identify and drop problematic markers from the genetic maps, including dropping linkage groups with 3 or fewer markers. Finally, we visually inspected our linkage groups using plotRF() from the Rqtl package, and merged linkage groups which had been incorrectly split up using the mergeCross() function from the ASmap package. Ultimately our final genetic maps included: 1) Crescent Pond: 214 individuals, 743 markers, 24 linkage groups and 2) Little Lake: 281 individuals, 540 markers, and 24 linkage groups (**Figure *2***).

**Figure 2.**
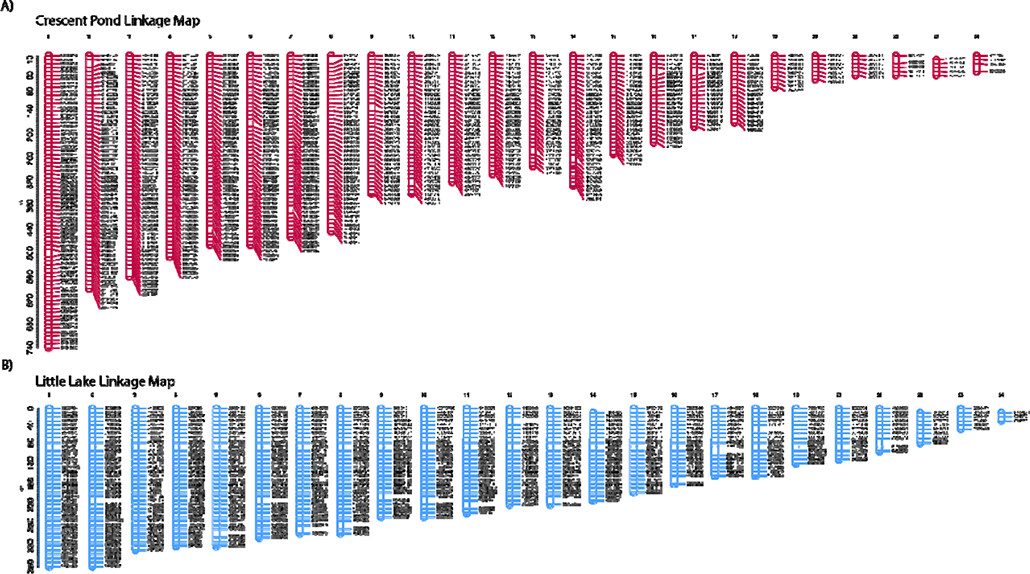
Linkage maps for A) Crescent Pond and B) Little Lake crosses. The Crescent Pond linkage map was estimated from 743 markers and the Little Lake linkage map was estimated from 540 markers. Both maps were generated from crosses between a scale-eater (*C. desquamator*) and snail-eater (*C. brontotheroides*) from the respective lakes.

#### QTL Analyses

We mapped QTL for 29 skeletal traits for both populations, and additional morphological (adductor mandibulae muscle mass) and behavioral traits (mate preference) for Crescent Pond. We used the Rqtl2 package (v0.22-11) to calculate genotype probabilities with a multipoint hidden Markov model using an error probability of 0.0001 and a Kosambi map function. We calculated kinship matrices to account for the relationship among individuals in two ways: 1) overall kinship, which represents the proportion of shared alleles between individuals, and 2) kinship calculated using the leave-one-chromosome-out method (LOCO). We used the scan1() function to perform three separate genome scans using a single-qtl model by: 1) Haley-Knott regression, 2) a linear mixed model using the overall kinship matrix, and 3) a linear mixed model using the LOCO kinship matrix. For our Crescent Pond data set we also included sex as an additive covariate. We assessed the significance of all three models using two significance thresholds *P* < 0.1 and 0.05 based on 1000 permutations each, using the scan1perm() function. As noted above the scan1() function can use several different methods to determine if a region is significantly associated with a given phenotype (Broman et al., 2019; Haley & Knott, 1992; Yang, Zaitlen, Goddard, Visscher, & Price, 2014; Yu et al., 2006), however, it is clear from previous theoretical work that many of these methods may suffer from type II error depending on the size of an organism’s genome, the density of markers in a linkage map, or the complexity of the phenotypic traits being measured (Lander and Botstein 1989; Risch 1990). We therefore relaxed the *P*-value cut off from 0.05 to 0.1 to capture potentially important genomic regions.

This relaxation is further supported by the LOD scores associated with regions significant at the *P*<0.1 level because they all exceed the traditional threshold of 3 (Nyholt 2000), the more conservative threshold of ∼3.3 (Lander and Kruglyak 1995; Nyholt 2000), the suggestive threshold of 1.86 (Lander and Kruglyak 1995), and are in line with estimates of significant LOD thresholds in previous studies (Erickson et al. 2016). All three of these methods detected similar QTLs and moving forward we only used the Haley-Knott regression method.

For each trait, we calculated the location of the maximum LOD score, and used the fit1() function to re-fit a single-QTL model at this location. We used the newly calculated LOD score to estimate the proportion of variance explained by the QTL and to calculate a *P*-value associated with each significant QTL ( ^2^-test). We also used the location of the maximum LOD score to calculate 95% Bayes credible intervals using the bayes_int() function from the Rqtl2 package. We note that the maximum LOD score associated with every trait across both ponds exceeded the suggestive threshold of 1.86 (Lander and Kruglyak 1995). We used the find.markerpos() function from Rqtl to determine where markers in each linkage map fell within the reference genome. With this information we were able to determine the scaffolds/positions from the reference genome that fell within the 95% credible intervals for each putative QTL. Finally, we used the maxmarg() function from the Rqtl2 package to find the genotype with the maximum marginal probability at the location of the maximum LOD. We used these genotypes to visualize the relationship between genotype and phenotypes.

#### Identifying adaptive alleles within QTL regions

For each scaffold that fell within a QTL’s credible interval we calculated the minimum and maximum position for that scaffold (that was identified in the putative QTL region) and searched the *C. brontotheroides* reference genome for annotated genes within the region. We then compared this list to a previously published list of genes that 1) contained or were adjacent to (within 20 kbp) fixed or nearly fixed (*Fst* > 0.95) SNPs between specialist species on SSI, and 2) showed significant evidence of a hard selective sweep in both the site frequency spectrum-based method SweeD (Pavlidis et al. 2013) and the linkage-disequilibrium-based method OmegaPlus (Alachiotis et al. 2012). We hereafter refer to these loci as adaptive alleles. We also noted whether adaptive alleles within QTL regions were classified as de novo, introgressed, or as standing genetic variation on SSI (Richards et al. 2021). We used a bootstrap resampling method to determine whether the observed proportions of adaptive alleles originating from de novo, introgression, or standing genetic variation found within QTL 95% credible intervals were different than the proportions expected when drawn from the genome at random. We used the boot (v. 1.3-25) package (Buckland et al. 1998; Canty and Ripley 2021) to resample our entire adaptive allele dataset (with replacement) 10,000 times. We then used the boot.ci() command from the boot package to calculated the 95% credible intervals around expected proportions of de novo, introgressed, and standing adaptive alleles. We performed these calculations separately for scale-eater and snail-eater adaptive alleles.

## Results

### Linkage Map Construction

We identified 24 linkage groups from 743 markers for Crescent Pond, and 24 linkage groups from 540 markers for Little Lake (**Figure *2***). Previous karyotypes of *Cyprinodon* species estimated 24 diploid chromosomes, matching the linkage groups in this study (Liu & Echelle, 2013; Stevenson, 1981). The total map length for Crescent Pond was 7335 cM and the total map length for Little Lake was 5330; the largest linkage groups for each map were 740 cM and 380 cM, respectively, and inter-marker map distance did not exceed 20cM in either map. To compare our maps and to determine if the same genomic regions were being reused across lakes, we identified where each marker was located in our reference genome. Overall, we found 324 markers in both maps that were within 10 Kbp of one another, indicating that 60% of the Little Lake map was also present in the Crescent Pond map and 44% of the Crescent Pond map was present in the Little Lake map (**Figure 3**).

**Figure 3.**
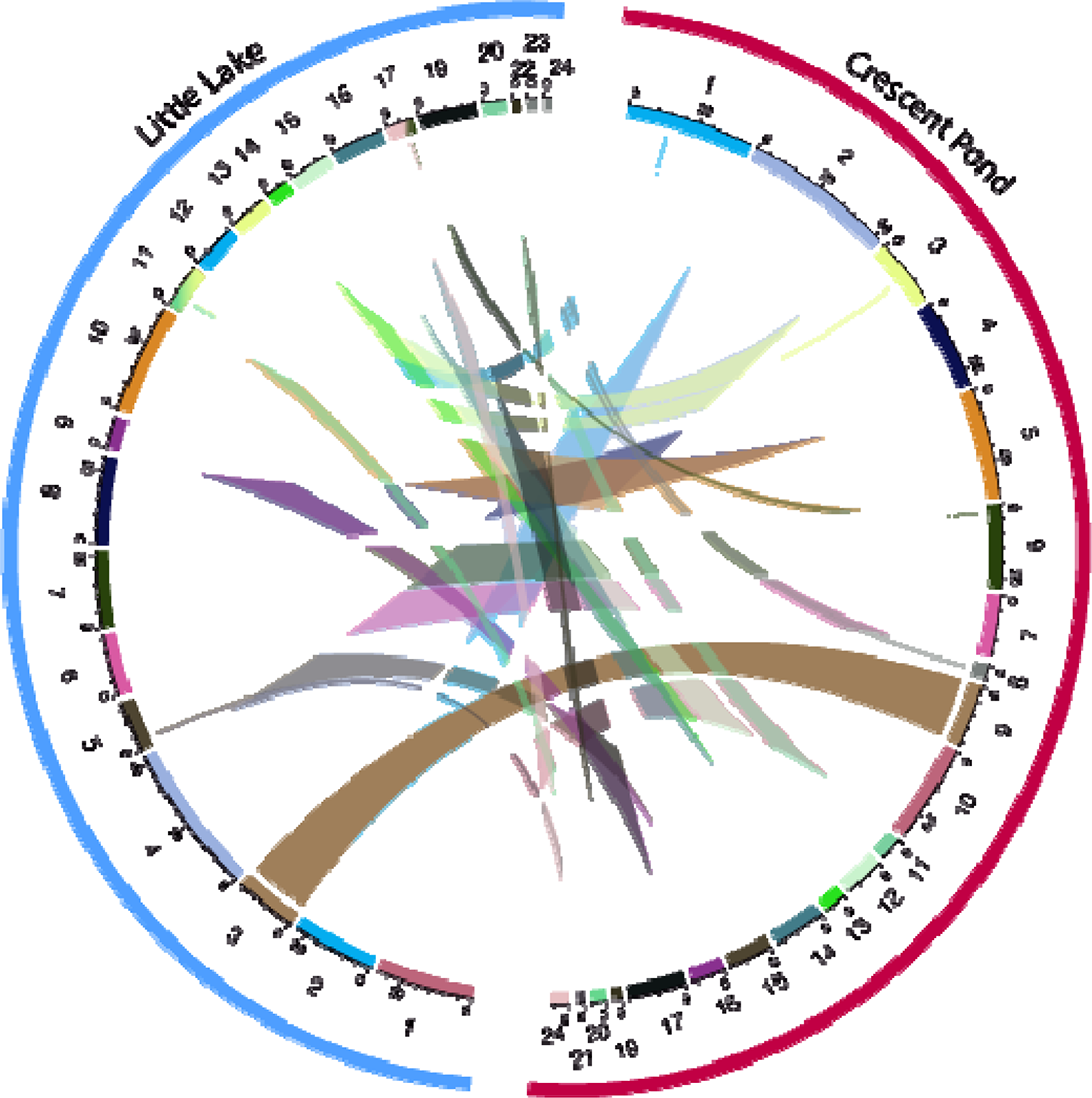
Circos plot depicting the relationship between the Crescent Pond (red) and Little L linkage maps (blue), which share 324 markers within 10 kbp of one another. Numbers surrounding each semi-circle represent linkage group numbers in each lake. Markers that are shared across lakes are connected via the colored lines.

### Craniofacial QTL

We detected three significant QTL in Crescent Pond and five QTL in Little Lake (**Table *1***, **Table *2***). In Crescent Pond, we identified QTL associated with the depth of the dentigerous arm of the premaxilla, cranial height, and adductor mandibulae muscle mass. Cranial height in Crescent Pond mapped to linkage group (LG) 10. Dentigerous arm depth and adductor mandibulae muscle mass both mapped to LG 13, which also contained the max LOD scores for two additional jaw traits (jaw opening in-lever and maxillary head height; **Table *2***). The 95% credible intervals for all these traits overlapped, suggesting that LG 13 may contain a single pleiotropic locus or many loci that affect all four traits.

**Table 1.**
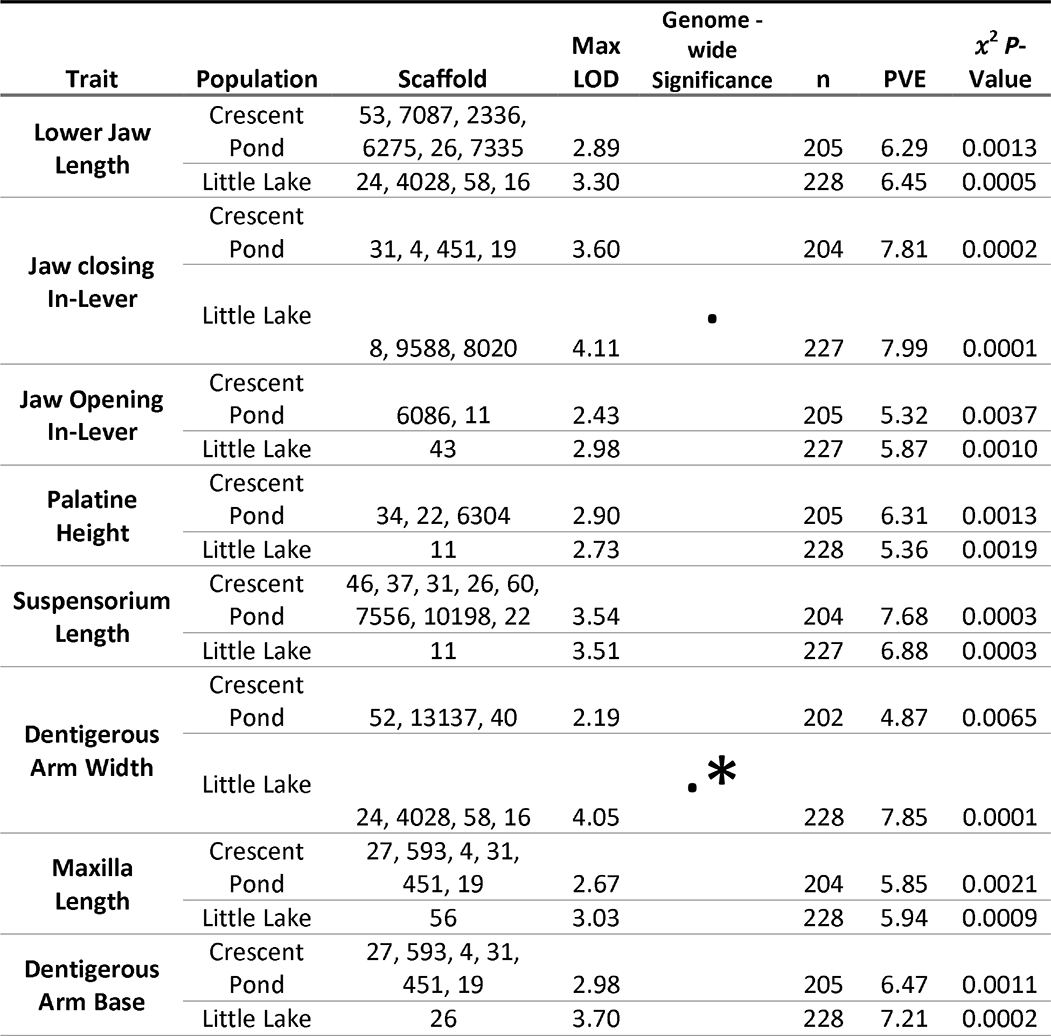
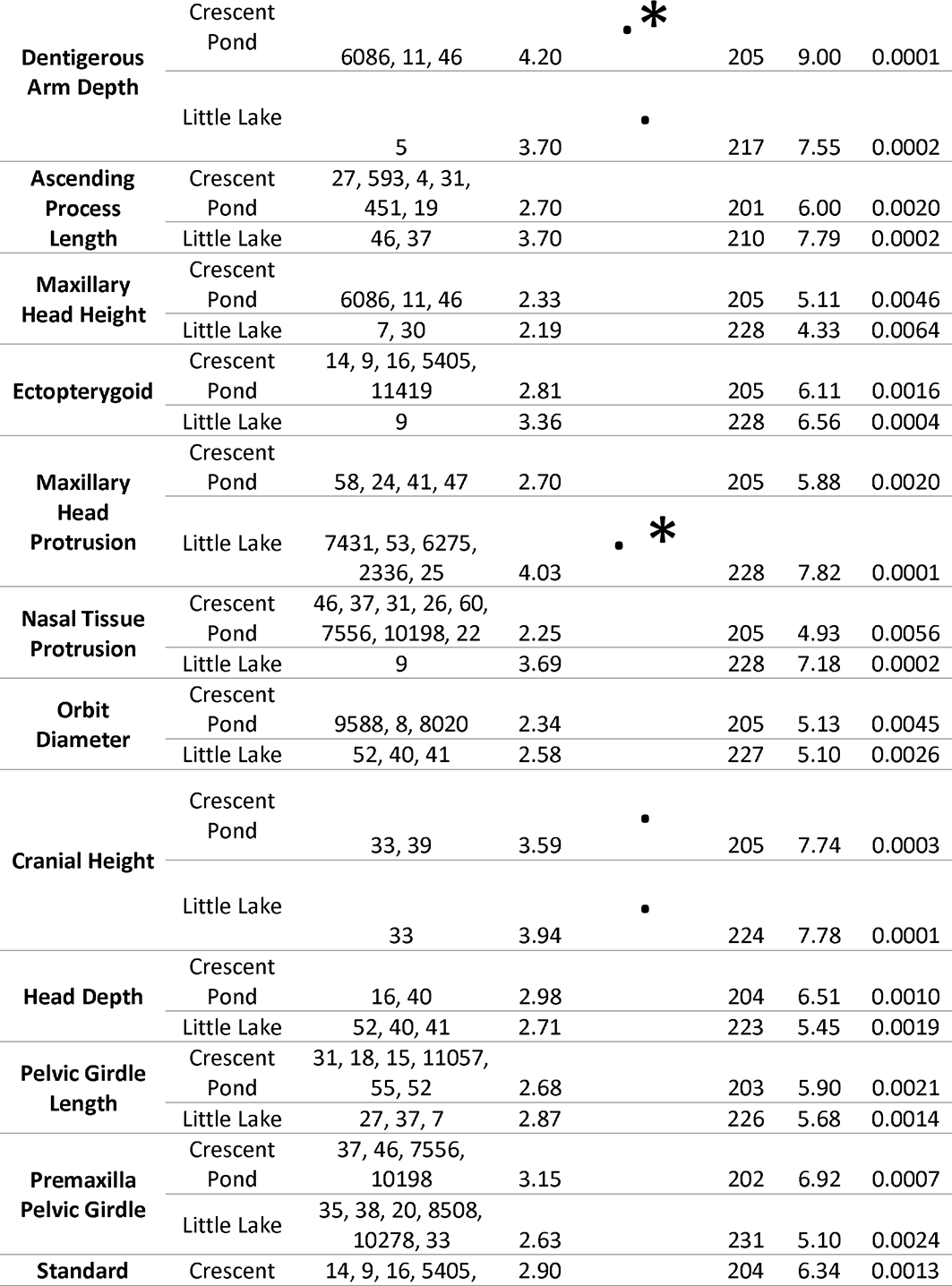
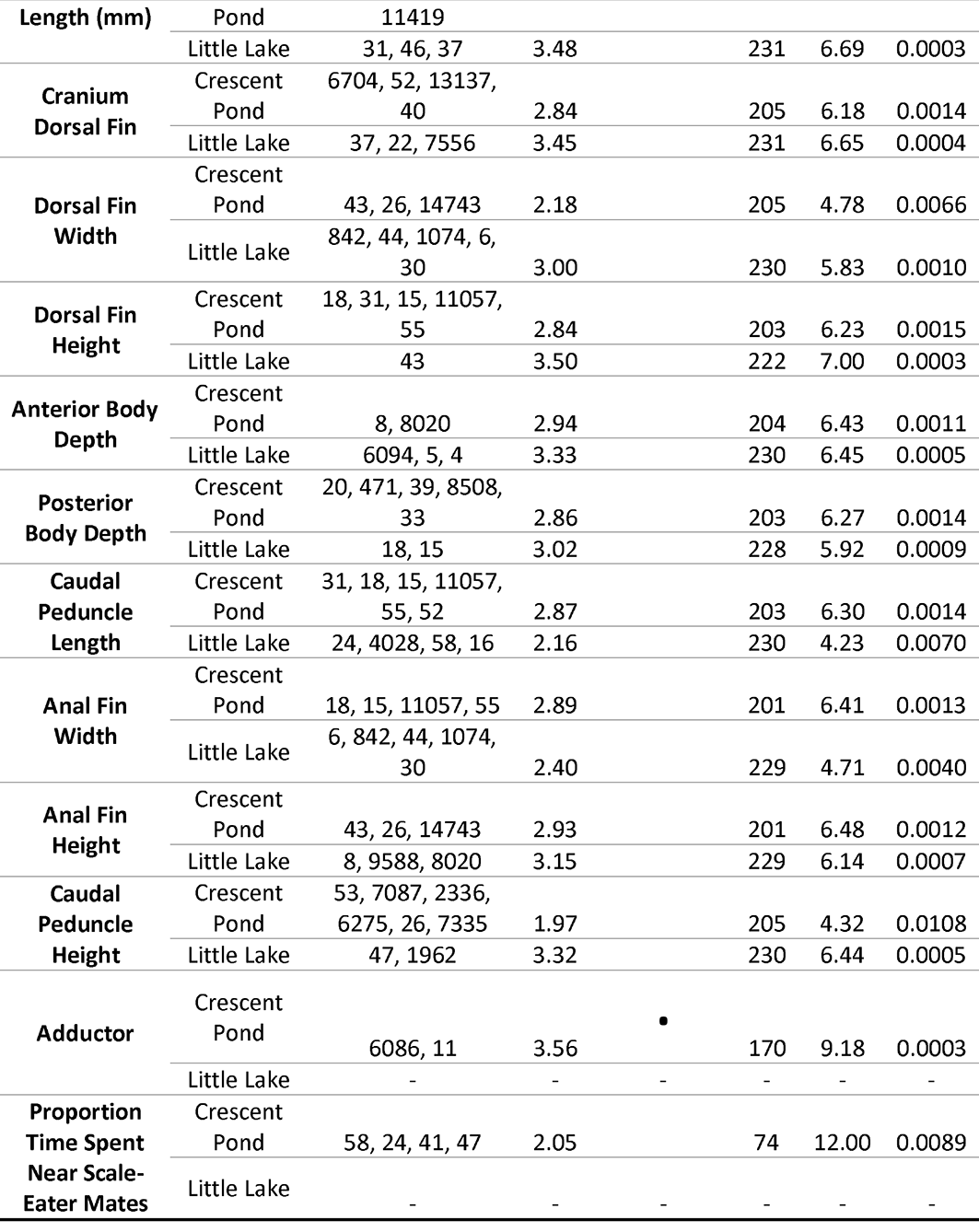
Maximum LOD scores for all 29 traits measured in Little Lake and Crescent Pond mapping crosses. A genome scan with a single-QTL model by Haley-Knott regression was used to identify the position with the highest LOD score, 95% Bayesian credible intervals, and the genome-wide significance level for each trait (*P*< 0.1: · ; *P* < 0.05: *). We also report the scaffold numbers of genomic regions that fell within the 95% credible intervals associated with the maximum LOD position for each trait, the number of individuals phenotyped, percent variance explained (PVE) by the max LOD region, and the uncorrected *P*-value associated with each max LOD region.

**Table 2.** Position of maximum LOD score and 95% credible intervals for each trait in the Little Lake linkage map and the Crescent Pond linkage map. Colors represent corresponding linkage groups across lakes. Asterisks represent traits that were marginally significant at the *P* < 0.1 level <u>in the genome scan.</u>

| Little Lake |  |  |  |  | Crescent Pond |  |  |  |  |
| --- | --- | --- | --- | --- | --- | --- | --- | --- | --- |
| Trait | Sig. | LG | Position<br>genomewide |  | Trait | Sig. | LG | Position<br>genomewide |  |
|  |  |  | Max LOD score | 95% CI |  |  |  | Max LOD score | 95% CI |
| Cranial Height | * | 1 | 259 | (250,270) | Suspensorium Length |  | 1 | 566 | (20,730) |
| Premaxilla to Pelvic Girdle |  | 1 | 146 | (0,350) | Nasal Tissue Protrusion |  | 1 | 570 | (0,740) |
| Cranium to Dorsal Fin |  | 2 | 303 | (160,380) | Premaxilla to Pelvic Girdle |  | 1 | 568 | (310,600) |
| Lower Jaw Length |  | 3 | 9 | (0,340) | Ectopterygoid |  | 3 | 272 | (0,560) |
| Dentigerous Arm Width | * | 3 | 9 | (0,340) | Standard Length (mm) |  | 3 | 50 | (40,500) |
| Caudal Peduncle Length |  | 3 | 168 | (0,340) | Dentigerous Arm Width |  | 4 | 317 | (40,510) |
| Dorsal Fin width |  | 4 | 187 | (10,310) | Cranium to Dorsal Fin |  | 4 | 89 | (30,510) |
| Anal Fin Width |  | 4 | 14 | (0,280) | Lower Jaw Length |  | 5 | 136 | (0,470) |
| Dentigerous Arm Depth | * | 6 | 79 | (20,90) | Caudal Peduncle height |  | 5 | 381 | (0,470) |
| Anterior Body Depth |  | 6 | 289 | (0,300) | Jaw Closing In-Lever |  | 6 | 380 | (150,410) |
| Orbit Diameter |  | 8 | 266 | (0,290) | Maxilla Length |  | 6 | 468 | (0,480) |
| Head Depth |  | 8 | 206 | (0,290) | Dentigerous Arm Base |  | 6 | 107 | (0,480) |
| Jaw Closing In-Lever | * | 9 | 54 | (40,90) | Ascending Proces Length |  | 6 | 106 | (0,470) |
| Anal Fin Height |  | 9 | 100 | (70,240) | Pelvic Girdle Length |  | 8 | 370 | (20,435) |
| Maxillary Head Protrusion | * | 10 | 35 | (0,260) | Dorsal Fin Height |  | 8 | 91 | (0,380) |
| Ascending Process Length |  | 12 | 119 | (90,150) | Caudal Peduncle Length |  | 8 | 258 | (30,425) |
| Standard Length (mm) |  | 12 | 200 | (50,210) | Anal Fin Width |  | 8 | 190 | (110,400) |
| Ectopterygoid |  | 13 | 170 | (150,180) | Maxillary Head Protrusion |  | 9 | 300 | (0,350) |
| Nasal Tissue Protrusion |  | 13 | 193 | (20,200) | Proportion Time Spent Near<br>Scale-Eater Males |  | 9 | 166 | (50,340) |
| Palatine Height |  | 14 | 147 | (110,210) | Cranial Height | * | 10 | 204 | (130,340) |
| Suspensorium Length |  | 14 | 153 | (70,180) | Posterior Body Depth |  | 10 | 270 | (0,330) |
| Jaw Opening In-Lever |  | 16 | 58 | (40,140) | Palatine Height |  | 11 | 70 | (0,310) |
| Dorsal Fin Height |  | 16 | 52 | (40,60) | Head Depth |  | 12 | 111 | (100,280) |
| Pelvic Girdle Length |  | 17 | 50 | (10,160) | Opening In-Lever |  | 13 | 10 | (0,90) |
| Maxillary Head Height |  | 18 | 122 | (30,160) | Dentigerous Arm Depth | * | 13 | 2 | (0,250) |
| Caudal Peduncle Height |  | 19 | 44 | (20,90) | Maxillary Head Height |  | 13 | 170 | (0,280) |
| Dentigerous Arm Base |  | 21 | 74 | (0,100) | Adductor Mandibulae Mass | * | 13 | 2 | (0,70) |
| Maxilla Length |  | 22 | 40 | (20,50) | Dorsal Fin Width |  | 14 | 305 | (30,330) |
| Posterior Body Depth |  | 24 | 30 | (10,30) | Anal Fin Width |  | 14 | 330 | (280,330) |
|  |  |  |  |  | Orbit Diameter |  | 16 | 107 | (0,190) |
|  |  |  |  |  | Anterior Body Depth |  | 16 | 170 | (10,220) |

In Little Lake, we detected significant QTL associated with jaw closing in-lever (i.e. height of the coronoid process on the articular: LG9), width and depth of the dentigerous arm of the premaxilla (LG3 and LG6), maxillary head protrusion (LG10), and cranial height (LG1; **Table *1***,**Table *2***). The 95% credible interval for dentigerous arm width on LG3 also contained the max LOD score for lower jaw length, suggesting that either a single pleiotropic locus or a cluster of loci in this region may be controlling both traits.

### Candidate genes and adaptive alleles within QTL regions

#### Cranial height

Cranial height was the only trait with statistically significant or marginally significant QTL in both lakes (Figure 4, *P* < 0.1). While the QTL occurred on different linkage groups between maps, we found a high degree of synteny between these linkage groups indicating that the QTL is located in the same genomic region in both lakes (**Table *2***, **Figure 3**). We also found the same overdominant genetic pattern in both lake crosses: heterozygotes showed increased cranial height relative to homozygous individuals (**Figure *5***).

**Figure 4.**
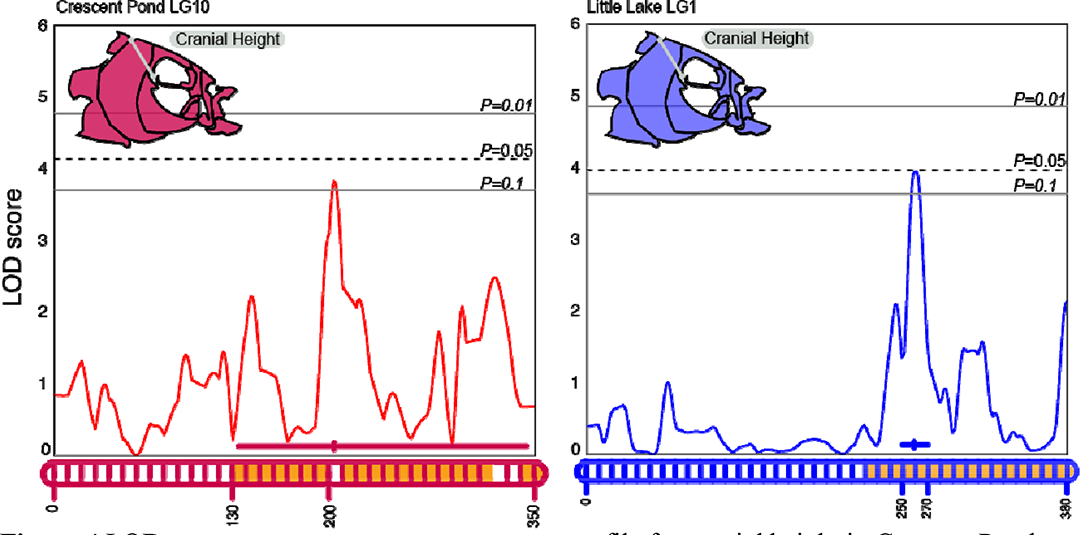
LOD profile for cranial height in Crescent Pond (red) and Little Lake (blue) F2 hybrids. LOD profiles were estimated by a Haley-Knott regression and are plotted relative to the position along the implicated linkage group (LG 10 for Crescent Pond, LG 1 for Little Lake) which are represented along the X-axis. Genome wide significance levels of P = 0.1, 0.05, and 0.01 are shown by the grey horizontal lines. Linkage groups along the X-axis also show the position of maximum LOD along with 95% Bayesian credible intervals. The orange fill color within the linkage groups corresponds to overlapping regions of scaffold 33 between crosses.

**Figure 5.**
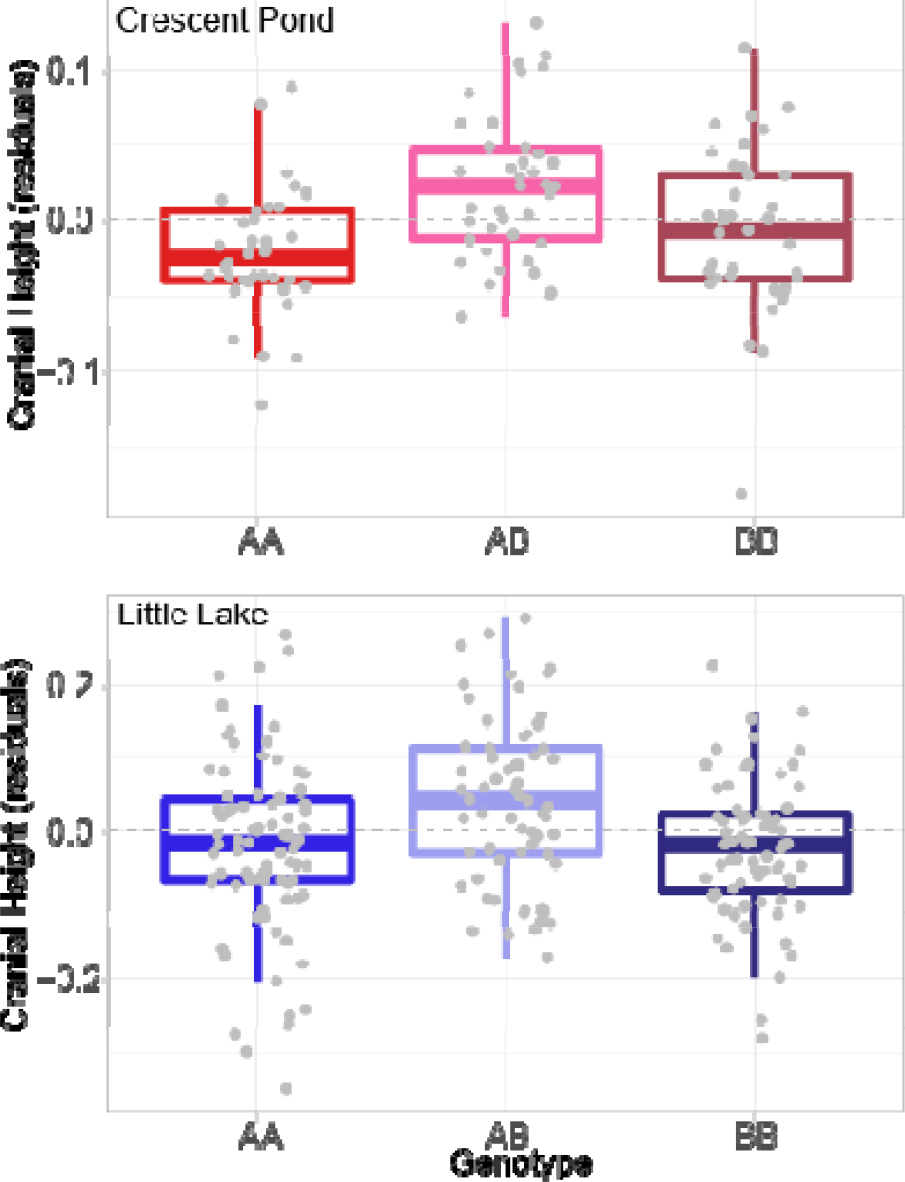
Cranial height phenotypes (size-corrected residuals) for each genotype in Crescent Pond (red) and Little Lake (blue). Both lakes show that heterozygotes (AB) exhibit greater cranial heights than homozygous parental genotypes.

We found 44 genes within scaffold 33 that fell partially or fully within the 95% credible intervals of the QTL in both lakes (Table 1, Table S1). Only three of these genes contained adaptive alleles within 20 kb: *wdr31*, *bri3bp*, and *gnaq* (**Table *3***). Interestingly, *gnaq* is well known to be associated with craniofacial development (Hall et al. 2007; Shirley et al. 2013) and is differentially expressed between our specialist species in developing larvae (McGirr and Martin 2020).

**Table 3.**
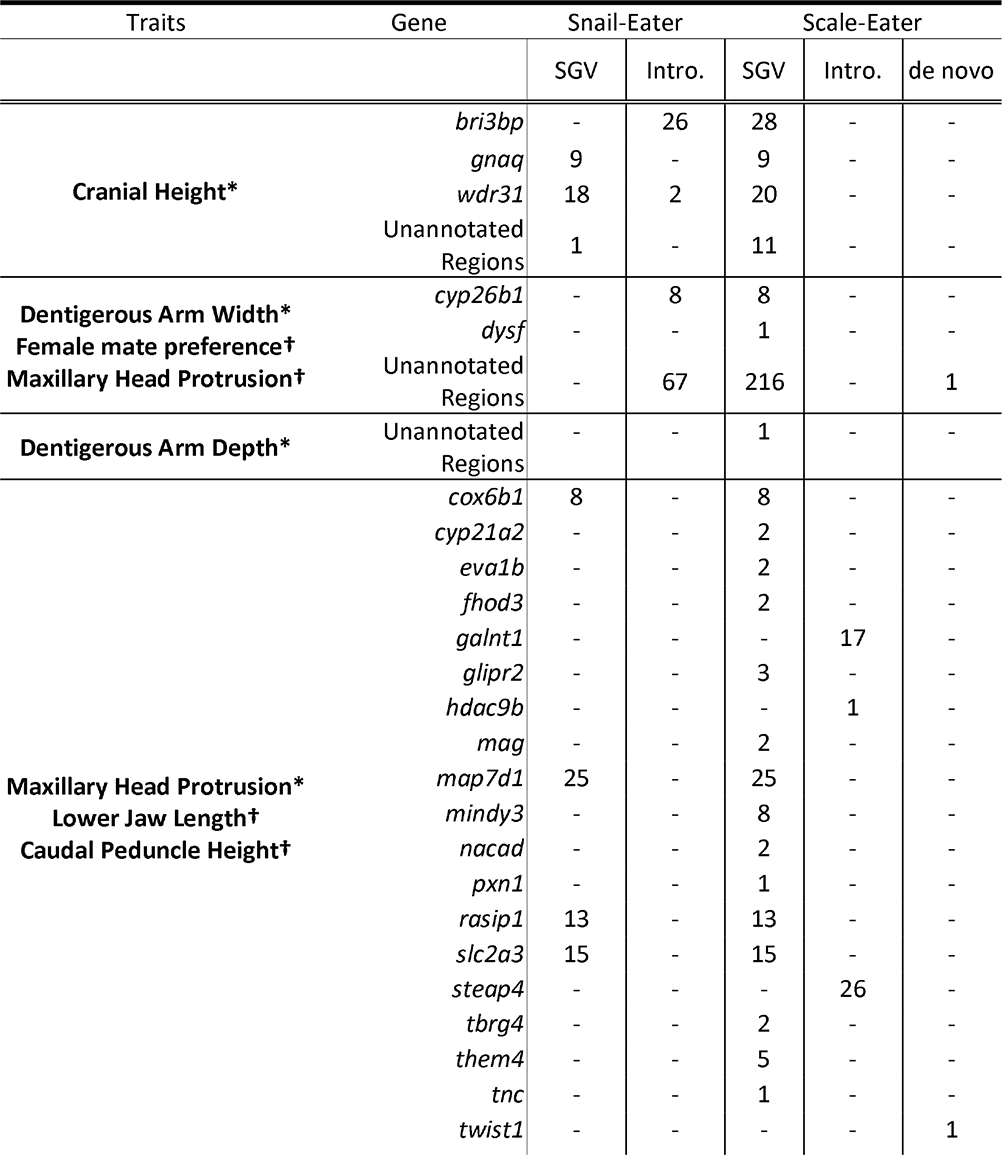
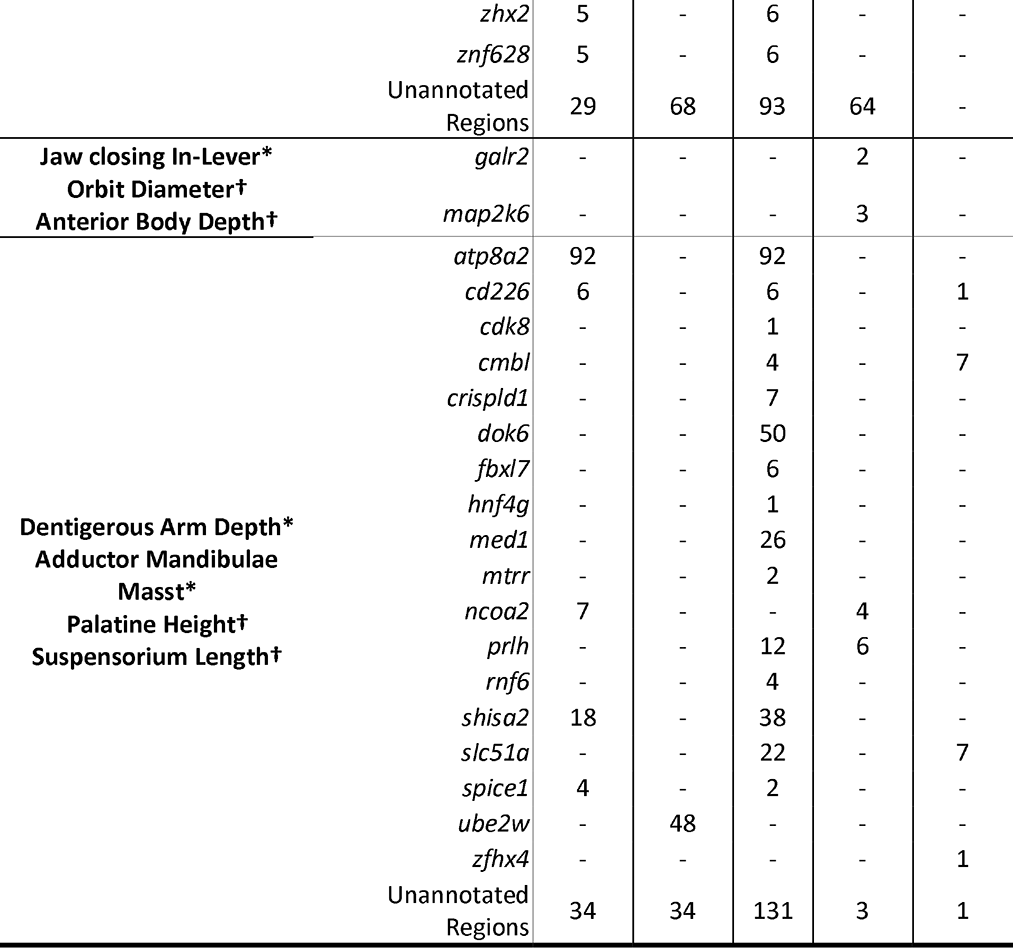
Number of adaptive alleles and any genes within 20 kbp found in trait QTL with maximum LOD scores for both lakes. Adaptive alleles were categorized as either standing genetic variation (SGV), introgression (Intro.), or de novo mutations, and were estimated independently for snail-eaters and scale-eaters in a previous study (Richards et al. 2021). Asterisks represent traits that were significant at the P < 0.1 level in the genome-wide scan, while crosses show traits that corresponded to the same locations in the alternate lake.

#### Dentigerous Arm Width

We found that regions on scaffolds 58 and 24 were associated with a significant QTL for dentigerous arm width in Little Lake and contained the max LOD scores for maxillary head protrusion and female mate preference in Crescent Pond (**Table *1***, **Table *2***). We found 161 genes which fell partially or completely within these shared regions, but only 2 genes, *dysf* and *cyp26b1*, which contained adaptive alleles within 20 kbp (**Table *3***). The *dysf* gene provides instructions for making a protein called dysferlin, which is found in the sarcolemma membrane that surrounds muscle fibers (Liu et al. 1998). This could indicate a role for muscle development in affecting skeletal development of the maxilla and premaxilla.

#### Dentigerous Arm Depth

The QTL for dentigerous arm depth in Little Lake was associated with LG 6, which corresponds to LG 7 in Crescent Pond, however, no traits from Crescent Pond mapped to this linkage group (**Table *2***, **Figure 3**). Instead, dentigerous arm depth in Crescent Pond was associated with LG 13 and did not share any similar genomic regions with those associated with dentigerous arm depth in Little Lake. We found 80 genes completely or partially within the 95% credible region for this QTL in Little Lak, but none contained adaptive alleles based on our criteria (Figure S1). In fact, only a single adaptive allele was found in this QTL region, but it was in an unannotated region of the genome (**Table *3***).

#### Maxillary Head Protrusion

Maxillary head protrusion in Little Lake mapped to a QTL region on LG10 which corresponds to the max LOD scores for both lower jaw length and caudal peduncle height in Crescent Pond (**Table *2***, **Figure 3**). Across lakes, all three traits were associated with scaffolds 53, 2336, and 6275. We found 528 genes partially or fully within these shared regions, but only 21 of these genes contained adaptive alleles within 20 kbp (**Table *3***). One of these genes, *twist1*, contains a non-synonymous substitution fixed in scale-eating pupfish on San Salvador Island, Bahamas (Richards et al. 2021). *Twist1* is a transcription factor and oncogene associated with palate development and oral jaw size in model organisms (Parsons et al. 2014; Teng et al. 2018).

#### Jaw Closing In-Lever

The QTL for jaw closing in-lever was associated with LG 9 in Little Lake, which corresponds to the max LOD scores for orbit diameter and anterior body depth in Crescent Pond (**Table *2***, **Figure 3**). Scaffolds 8 and 8020 were associated with all three of these traits. We found 13 genes which partially or completely fell within these shared regions, and only two genes, *map2k6* and *galr2*, which contained adaptive alleles within 20 kbp (**Table *3***). *Galr2* was also previously detected within a significant QTL for lower jaw length in pupfish (Martin et al. 2017).

#### Dentigerous Arm Depth and Adductor Mandibulae Muscle Mass

Finally, in Crescent Pond the QTL for dentigerous arm depth and adductor mandibulae muscle mass mapped to the exact same location on LG 13 (95% CI dentigerous arm depth (0, 250), adductor mandibulae muscle mass (0,70). This linkage group corresponds to LG14 in Little Lake, which contains the max LOD scores for both palatine height and suspensorium length (**Table *3***). We found 52 genes that overlapped between these regions, 18 of which contained adaptive alleles. Furthermore, three of the genes— *ube2w, ncoa2, and prlh*—contained adaptive alleles that introgressed from Laguna Bavaro in the Dominican Republic to snail-eating pupfish (*ube2w*), from Lake Cunningham, New Providence Island to scale-eating pupfish (*ncoa2*), or from North Carolina, USA to scale-eating pupfish (*prlh)*. We also found four genes that contained adaptive alleles within 20 kbp that arose from de novo mutations: *cd226, cmbl, slc51a,* and *zfhx;* however, only one adaptive allele in *slc51a* is found within a coding region.

### Origins of adaptive alleles

Adaptive alleles originating from standing genetic variation across the Caribbean were most common within shared QTL regions between lakes (86.03% within scale-eater populations, and 53.32% within snail-eating populations; **Table *3***). However, observed proportions within shared QTL were significantly less than expected by chance (scale-eater expected 95% CI: (88.33%-90.37%), snail-eater expected 95% CI: (62%-67%;10,000 bootstrapped iterations). Instead, we found more introgressed scale-eater and snail-eater adaptive variants in shared QTL regions than expected by chance (Scale-eater observed: 12.13% introgressed, scale-eater expected 95% CI: (7.96%-9.88%); Snail-eater observed: 46.67% introgressed, snail-eater expected 95% CI: (32.22%-37.06%)). Finally, we found that about 1.83% of adaptive alleles within overlapping regions between lakes originated from de novo mutations in scale-eaters, however, this fell within the predicted null range (95% CI: (1.29%-2.17%)).

## Discussion

### Parallel genetic changes underlie 5 out of 6 of craniofacial QTL

We found evidence supporting both parallel and non-parallel genetic changes in an adaptive radiation of trophic specialist pupfishes. A single significant QTL was associated with cranial height in both lakes and mapped to the same genomic region, suggesting that parallel genetic changes are responsible for variation in this trait in both lakes. On the other hand, significant QTLs were identified for premaxilla dentigerous arm depth in each lake, but they mapped to different locations, indicating that this trait is associated with non-parallel genetic changes. We found an additional three traits with significant QTLs (dentigerous arm width, jaw closing in-lever, maxillary head protrusion) in the Little Lake population that were not detected in Crescent Pond. However, all genomic regions associated with these traits in Little Lake also mapped to the max LOD score for this same integrated suite of craniofacial traits in Crescent Pond. Therefore, rather than assume independent QTL for each trait, we conservatively conclude that the same genomic regions are being reused in each lake and affect a highly integrated suite of craniofacial traits. Overall, we found that 5 out of the 6 significant QTLs were reused in some way across lakes suggesting that parallel genetic changes underly adaptive phenotypes in the San Salvador Island pupfish radiation.

### High level of QTL reuse across ponds

Overall, we found that about 16% (1 out of 6) of the identified QTL regions corresponded to non-parallel changes and 84% (5 out of 6) corresponded to parallel genetic changes—either affecting the same phenotypic trait or a tightly correlated craniofacial trait— across populations. The presence of both non-parallel and parallel genetic changes leading to convergent phenotypes across lakes has been documented previously. For example, Colosimo et al. (2004) investigated the genetic basis of armor plate morphology in two independent threespine stickleback populations and found a single large effect locus on LG 4 in the two populations. However, they also noted a potential difference in the dominance relationships of alleles across ponds at this location, and found additional differences in modifier QTLs between populations, suggesting that both parallel and non-parallel genetic changes could lead to the loss of armor plating. Similarly, Erickson et al. (2016) found evidence for both parallel (43% of QTL regions overlapped between at least two populations) and non-parallel (57% of QTL regions were found in only a single population) evolution in a QTL study investigating the genetic basis of 36 skeletal phenotypes in three independent threespine stickleback populations. However, our findings suggest that pupfish exhibit a much higher proportion of parallel evolution than previously documented in stickleback. In fact, Conte et al. (2012) estimated that the probability of convergence via gene reuse is only 32-55% —which is 1.5 to 2.5 times lower than our current finding— although this may be underestimated (Stern 2013).

Pupfish may have a higher rate of parallel evolution than other model fish speciation systems for a few reasons. First, the pupfish radiation is recent, although comparable in age to glacial stickleback populations, with specialist species diverging less than 10kya (Hagey and Mylroie 1995; Martin and Wainwright 2013), and parallel evolution is predicted to be more likely when populations or species have recently diverged (Rosenblum et al. 2014). This may be because recently diverged species are more likely to experience similar environments, have access to similar pools of genetic variation (either due to standing genetic variation or introgression), or similar genetic constraints. Second, the genomic basis of pupfish skeletal traits may be primarily controlled by cis-regulatory elements, which evolve more quickly and have less negative pleiotropy which may make them more likely to undergo parallel evolution (Stern and Orgogozo 2008). However, a previous study of allele-specific expression in the pupfish system found strong evidence that two cis-regulatory alleles were associated with skeletal development, but trans-acting elements predominated overall (McGirr and Martin 2021).

In part, the increased proportion of parallel evolution estimated in this study results from our relaxed thresholds for detecting and categorizing shared QTL regions. Previous QTL studies have typically searched for evidence of parallel evolution by only looking for one-to-one mapping in which the same genomic regions are associated with the same trait across populations at a genome-wide level of significance in each (Colosimo et al. 2004; Conte et al. 2012). While this method provides the most clear-cut examples of parallel evolution, we argue that it vastly underestimates its frequency in nature. For example, this method would not consider reuse of the same genomic regions for integrated morphological traits as parallel evolution, a pattern seen in this study and in Erickson et al. (2016). Furthermore, the strict one-to-one significance method for detecting parallel evolution does not include consideration of the hierarchy and diversity of convergence and parallel evolution, which can span morphological traits, ecotypes, performance, or even fitness (James et al. 2020; Rosenblum et al., 2014; Stern, 2013; Martin and Wainwright 2013). Ultimately, we argue that our method of quantifying parallel evolution provides a more wholistic view of the process and better captures the frequency of reuse of adaptive genetic variation in nature.

### Few QTL may affect many highly integrated craniofacial traits

There are several processes that may cause the same genomic regions to be associated with different traits between lakes. First, these genomic regions may be highly pleiotropic and affect several traits simultaneously. For example, Albert et al. (2007) found that that on average a single QTL affected 3.5 phenotypic traits in an analysis of 54 body traits in three-spine stickleback. Wagner et al. (2008) found a similar pattern in QTL analyses of 70 skeletal traits in mice, where a single QTL affected on average 7.8 phenotypic traits (the maximum being 30).

A second possibility is that a single QTL region may contain several tightly linked causative variants that are responsible for variation in many traits. Correlated phenotypic traits are generally assumed to have a shared genetic basis, but it is extremely difficult to determine if this is due to pleiotropy or tight linkage between genomic regions (Lynch and Walsh 1998; Gardner and Latta 2007; Paaby and Rockman 2013; Wright et al. 2010).

Finally, it may be that differences in methodology or sample sizes between lakes enable us to detect significant QTL for some traits in one lake and not the other. For example, our analyses of Little Lake allowed us to detect significant QTL for effect sizes greater than 6.54 PVE at 80% power, but we could only detect significant QTL for effect sizes greater than 8.41 PVE at 80% power in Crescent Pond due to our lower sample size for this cross (Sen et al. 2007). However, this level of power is typical in many non-model QTL studies (Ashton et al. 2017). The ability to detect a significant QTL in one lake but not the other may be further explained by our use of different sequencing methods between populations. However, a critical component of our analyses involved searching for regions within 10 kbp of one another across maps to provide confidence that if we detected a significant QTL in one lake and not the other that it was not simply because that genomic region was not captured by the sequencing. For example, in Little Lake we detected a significant QTL associated with dentigerous arm depth on LG 6 but did not find any traits associated with this region of the genome in Crescent Pond.

### QTL are associated with different craniofacial traits across different lakes

In this study we found an intriguing pattern of different traits mapping to the same region of the genome across lake populations. One potential explanation for this is that there are different relationships between traits in each lake, and we find some evidence of this in our phenotypic data. Figure S2 depicts correlation matrices between traits in 1) Little Lake and 2) Crescent Pond, and ^2^ comparisons of these two matrices reveals that the relationship between traits varies significantly between lakes ( ^2^=3135.99, df = 756, *P*< 3.6e-29). For example, the relationship between maxillary head protrusion and lower jaw length is more than two times stronger in Little Lake compared to Crescent Pond (Pearson’s r_LL_=0.27, Pearson’s r_CP_ = 0.12), the relationship between dentigerous arm depth and suspensorium length is 1.8 times stronger in Little Lake than in Crescent Pond (Pearson’s r_LL_=0.45, Pearson’s r_CP_ = 0.24), and the relationship between jaw closing in-lever and anterior body depth is more than two times stronger in Crescent Pond than in Little Lake (Pearson’s r_CP_ = 0.23, Pearson’s r_LL_=0.11).

This pattern may be explained by different epistatic interactions in each lake. For example, Juenger et al. (2005) detected significant QTL-QTL interactions in one mapping population of *Arabidopsis* but found no evidence of the same interactions in the other population. When we investigated the relationship between phenotype and genotype for cranial height, we found the same overdominance pattern in both lakes (Figure 5). However, the presence of epistatic interactions may also be an obstacle for QTL detection. In a mapping study of body weight in chicken, Carlborg et al. (2006) were only able to detect a single weak QTL despite the extremely divergent phenotypes between parental lines. However, when accounting for epistatic interactions, Carlborg et al. identified several additional significant QTL regions that explained a large amount of variation in body weights.

Finally, our method for searching for putative QTL regions may have led to this pattern. Similar studies have searched for influential genomic regions by first identifying a putative QTL in a single population, and then searching the already identified linkage group in the second population for any signal of a QTL associated with the same phenotype, often using relaxed LOD thresholds closer to the suggestive cut-off (LOD> 1.8, e.g., Erickson et al. 2016). Our approach, however, independently identified the positions of maximum LOD for all traits across the entire linkage map before searching for similar implicated regions between populations. We argue that our approach minimizes bias, because there are no prior expectations about which traits should be associated with a given genomic region within a suite of integrated traits, and reduces false positives because we only examine the maximum LOD position for each trait.

### Identifying causative regions within QTL

Multiple mapping populations across lakes may also be particularly useful for identifying candidate causal alleles. We found that one out of our six unique QTL regions mapped to the same genomic location across lakes and was associated with the same phenotypic trait—cranial height (Figure 4). In Crescent Pond, we found that a region of 110 cM was associated with this trait (LG10, position: 204, 95% CI (130,340)), which contained 426 genes. However, when we compared this region to the region independently identified in our Little Lake analysis, we found that the overlapping region was reduced to 20cM (LG1, position: 259, 95% CI (250-270)) and contained only 44 genes—a reduction of more than 80%. We found a similar pattern in the additional four QTL regions that mapped to the same genomic location across maps but were associated with different phenotypic traits and observed an average 56% reduction in region size. As noted above, Erickson et al. (2016) used a similar method of identifying candidate QTL regions across three hybrid populations of stickleback, and found that 43% of identified QTL regions were shared across two or more populations; however, they did not investigate whether these QTL regions completely or partially overlapped.

We also searched for adaptive alleles within QTL region that were identified in a previous study as 1) nearly fixed between species (F_st_ >0.95) and 2) showed significant evidence of a hard selective sweep (Richards et al. 2021). Overall, we found 789 shared genes within shared QTL regions across lakes, and that 45 of those genes contained adaptive variants (5.7%). This is a six-fold increase from the genome-wide expectation of 0.91% (176 genes associated with at least one adaptive variant / 19304 annotated genic regions), suggesting that these specific regions are important for adaptation to scale- and snail-feeding in wild pupfish. For example, a variant in *twist1* was found within the region associated with maxillary head protrusion in Little Lake (which also overlapped with lower jaw length and caudal peduncle height in Crescent Pond). In model organisms, *twist1* is associated with palate and jaw development (Parsons et al. 2014; Teng et al. 2018), and previous genome-wide association scans in pupfish showed that a region containing *twist1* was significantly associated with oral jaw size in the system (Richards et al. 2021). Similarly, we found that variants associated with *galr2* fell within the QTL region associated with jaw closing in-lever in Little Lake (which also overlapped with regions associated with orbit diameter and anterior body depth in Crescent Pond; scaffolds 8 and 8020), and previous QTL mapping studies, gene expression studies, and genome-wide association analyses have all implicated regions containing *galr2* with oral jaw development in pupfish (McGirr and Martin 2016; Martin et al. 2017; Richards et al. 2021).

### Increased use of introgressed adaptive variants in QTL regions

We found that most genetic variation within shared QTL regions was also segregating across outgroup Caribbean generalist populations characterized by Richards et al. (2021; 86.04% within scale-eater populations, and 53.32% within snail-eating populations). Furthermore, we found more introgressed adaptive alleles from both scale-eater (observed: 12.13% introgressed, expected 95% CI: (7.96%-9.88%)) and snail-eater populations in shared QTL regions than expected by chance (observed: 46.67%, expected 95% CI: (32.22%-37.06%)). This supports the prediction that standing genetic variation and introgressed variation should underlie parallel genetic changes (Stern 2013; Thompson et al. 2019). Finally, we found that only 1.83% of adaptive alleles within shared QTL regions across both lakes originated from de novo mutations on San Salvador Island. While this percentage did not differ significantly from the expected estimates (expected 95% CI: 1.3%-2.17%) it does not eliminate the possibility that de novo mutations play an important adaptive role in pupfish evolution.

## Conclusion

In conclusion, we found that a single QTL region was responsible for variation in cranial height in both populations, and an additional four QTL regions were responsible for variation in different craniofacial traits across lakes, suggesting that parallel genetic changes underlie integrated suites of adaptive craniofacial phenotypes on San Salvador Island. Adaptive alleles were more commonly found within these detected QTL regions, and more of these adaptive alleles arrived on SSI via introgression than expected by chance. Finally, we argue that investigating QTL regions across populations in concert with estimation of hard selective sweeps in wild populations is a powerful tool for identifying potential causative regions of the genome affecting adaptive divergence.

## Acknowledgements

We thank the University of California, Davis, the University of California, Berkeley, the University of North Carolina at Chapel Hill, NSF CAREER 1749764, NIH 5R01DE027052-02, and BSF 2016136 for funding to CHM. The Bahamas Environmental Science and Technology Commission and the Ministry of Agriculture provided permission to export fish and conduct this research. Rochelle Hanna, Velda Knowles, Troy Day, and the Gerace Research Centre provided logistical assistance in the field. All animal care protocols were approved by the University of California, Davis and the University of California, Berkeley Animal Care

## Author Contributions

MESJ and CHM designed research; MESJ, CHM, JCD, and SR performed data collection; MESJ and EJR performed data analysis; MESJ and CHM wrote the paper. CHM provided funding.

## Data Accessibility

Data will be deposited to Dryad and NCBI. Genomes are archived at the National Center for Biotechnology Information BioProject Database (Accessions: PRJNA690558; PRJNA394148, PRJNA391309; and PRJNA305422).

