## Supplemental Tables and Figures for "Parallel genetic changes underlie integrated craniofacial traits in an adaptive radiation of trophic specialist pupfishes"

**Supplemental Figures and Tables:**

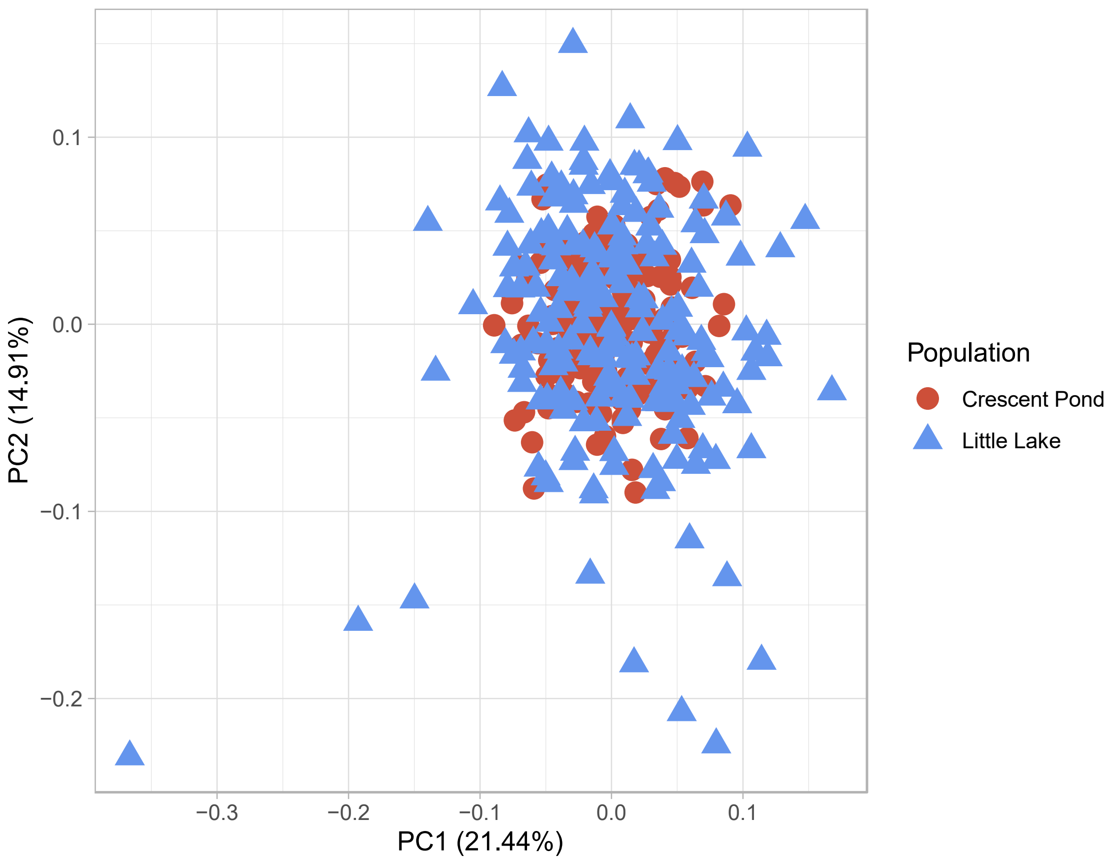


**Figure S1** Principal component analysis depicting phenotypic variation across 28 size corrected skeletal traits for Crescent Pond F2 hybrids (red circles) and Little Lake F2 hybrids (blue triangles). Skeletal traits were calculated as the mean from two lateral photographs from each individual. There was no significant difference between Crescent Pond and Little Lake individuals (MANOVA, df = 28, approximate F-value= 0.34, $P = 1$).

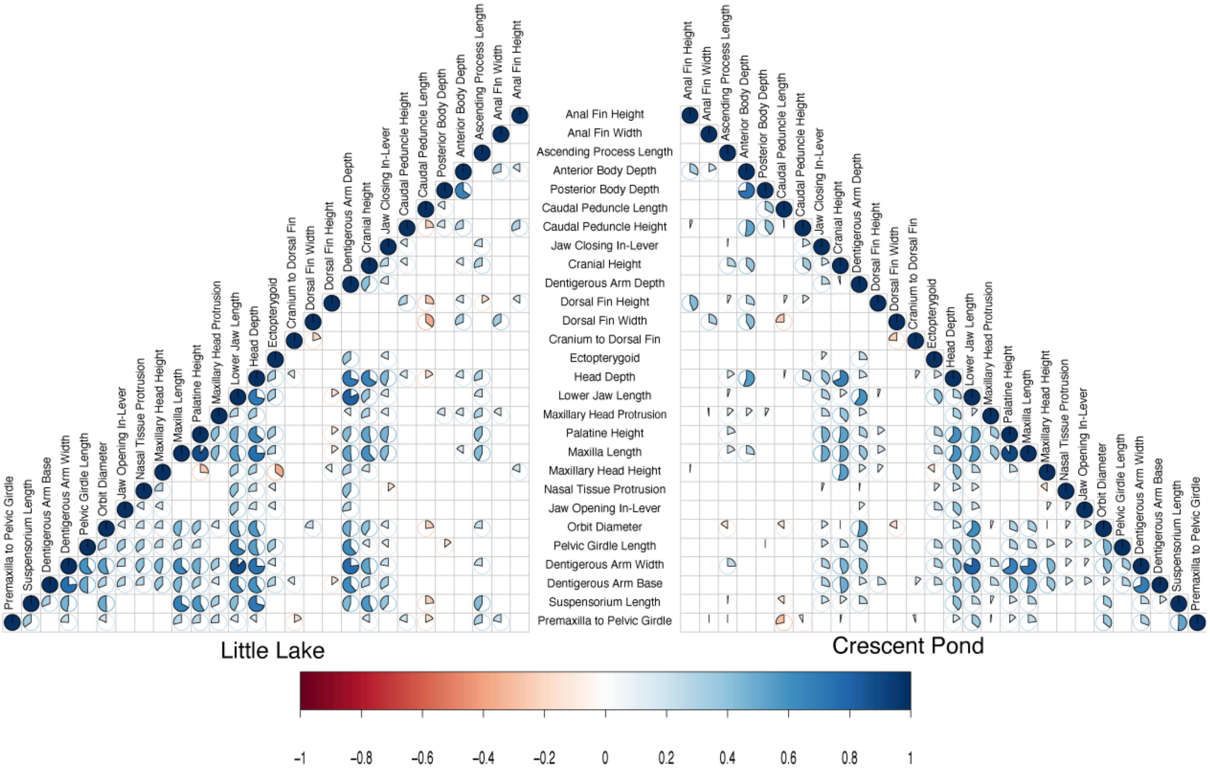


**Supplemental Figure S2** Correlation matrices depicting the relationship between phenotypic traits in Little Lake and Crescent Pond. Visualized pie charts represent relationships that are significant at the $P < 0.05$ level. Red pie charts represent negative relationships and blue pie charts represent positive relationships.

**Gene Table Supplement:**

**Supplemental Table 1** List of genes that fall within or partially within significant QTL regions.

| Trait | Gene | Scaffold | Population |
|---|---|---|---|
| Adductor Mandibulae Height | abca4 | HiC_scaffold_11 | CP |
| Adductor Mandibulae Height | agrp | HiC_scaffold_11 | CP |
| Adductor Mandibulae Height | ahrr | HiC_scaffold_11 | CP |
| Adductor Mandibulae Height | arhgap29 | HiC_scaffold_11 | CP |
| Adductor Mandibulae Height | b2m | HiC_scaffold_11 | CP |
| Adductor Mandibulae Height | bco1 | HiC_scaffold_11 | CP |
| Adductor Mandibulae Height | bloc1s5 | HiC_scaffold_11 | CP |
| Adductor Mandibulae Height | brpf3 | HiC_scaffold_11 | CP |
| Adductor Mandibulae Height | ccdc63 | HiC_scaffold_11 | CP |
| Adductor Mandibulae Height | cebpe | HiC_scaffold_11 | CP |
| Adductor Mandibulae Height | cngb3 | HiC_scaffold_11 | CP |
| Adductor Mandibulae Height | col11a1 | HiC_scaffold_11 | CP |
| Adductor Mandibulae Height | cpne3 | HiC_scaffold_11 | CP |
| Adductor Mandibulae Height | dcaf11 | HiC_scaffold_11 | CP |
| Adductor Mandibulae Height | dph2 | HiC_scaffold_11 | CP |
| Adductor Mandibulae Height | eef1e1 | HiC_scaffold_11 | CP |
| Adductor Mandibulae Height | emp3 | HiC_scaffold_11 | CP |
| Adductor Mandibulae Height | fam168b | HiC_scaffold_11 | CP |
| Adductor Mandibulae Height | fam83e | HiC_scaffold_11 | CP |
| Adductor Mandibulae Height | fen1 | HiC_scaffold_11 | CP |
| Adductor Mandibulae Height | fitm1 | HiC_scaffold_11 | CP |
| Adductor Mandibulae Height | foxj3 | HiC_scaffold_11 | CP |
| Adductor Mandibulae Height | garem1 | HiC_scaffold_11 | CP |
| Adductor Mandibulae Height | glyctk | HiC_scaffold_11 | CP |
| Adductor Mandibulae Height | hbp1 | HiC_scaffold_11 | CP |
| Adductor Mandibulae Height | hmcn2 | HiC_scaffold_11 | CP |
| Adductor Mandibulae Height | kazn | HiC_scaffold_11 | CP |
| Adductor Mandibulae Height | kazna | HiC_scaffold_11 | CP |
| Adductor Mandibulae Height | kbp | HiC_scaffold_11 | CP |
| Adductor Mandibulae Height | kif20a | HiC_scaffold_11 | CP |
| Adductor Mandibulae Height | mag | HiC_scaffold_11 | CP |
| Adductor Mandibulae Height | mak | HiC_scaffold_11 | CP |
| Adductor Mandibulae Height | mcur1 | HiC_scaffold_11 | CP |
| Adductor Mandibulae Height | mon1b | HiC_scaffold_11 | CP |
| Adductor Mandibulae Height | myh6 | HiC_scaffold_11 | CP |
| Adductor Mandibulae Height | myh7 | HiC_scaffold_11 | CP |
| Adductor Mandibulae Height | ngdn | HiC_scaffold_11 | CP |

| | | | |
|---|---|---|---|
| Adductor Mandibulae Height | nrros | HiC_scaffold_11 | CP |
| Adductor Mandibulae Height | ntng1 | HiC_scaffold_11 | CP |
| Adductor Mandibulae Height | olfm3 | HiC_scaffold_11 | CP |
| Adductor Mandibulae Height | pabpn1 | HiC_scaffold_11 | CP |
| Adductor Mandibulae Height | pck2 | HiC_scaffold_11 | CP |
| Adductor Mandibulae Height | pdcd6 | HiC_scaffold_11 | CP |
| Adductor Mandibulae Height | phldb2 | HiC_scaffold_11 | CP |
| Adductor Mandibulae Height | pigm | HiC_scaffold_11 | CP |
| Adductor Mandibulae Height | pkia | HiC_scaffold_11 | CP |
| Adductor Mandibulae Height | plch2 | HiC_scaffold_11 | CP |
| Adductor Mandibulae Height | plcxd2 | HiC_scaffold_11 | CP |
| Adductor Mandibulae Height | pomp | HiC_scaffold_11 | CP |
| Adductor Mandibulae Height | ppcs | HiC_scaffold_11 | CP |
| Adductor Mandibulae Height | prdm2 | HiC_scaffold_11 | CP |
| Adductor Mandibulae Height | prkdc | HiC_scaffold_11 | CP |
| Adductor Mandibulae Height | prmt6 | HiC_scaffold_11 | CP |
| Adductor Mandibulae Height | psme1 | HiC_scaffold_11 | CP |
| Adductor Mandibulae Height | ramp3 | HiC_scaffold_11 | CP |
| Adductor Mandibulae Height | rec8 | HiC_scaffold_11 | CP |
| Adductor Mandibulae Height | rgs9bp-b | HiC_scaffold_11 | CP |
| Adductor Mandibulae Height | rmdn1 | HiC_scaffold_11 | CP |
| Adductor Mandibulae Height | rnpc3 | HiC_scaffold_11 | CP |
| Adductor Mandibulae Height | rps20 | HiC_scaffold_11 | CP |
| Adductor Mandibulae Height | serpinb1 | HiC_scaffold_11 | CP |
| Adductor Mandibulae Height | serpinb10 | HiC_scaffold_11 | CP |
| Adductor Mandibulae Height | serpinb1b | HiC_scaffold_11 | CP |
| Adductor Mandibulae Height | serpinb6 | HiC_scaffold_11 | CP |
| Adductor Mandibulae Height | sh3glb1 | HiC_scaffold_11 | CP |
| Adductor Mandibulae Height | siglec1 | HiC_scaffold_11 | CP |
| Adductor Mandibulae Height | siglec10 | HiC_scaffold_11 | CP |
| Adductor Mandibulae Height | siglec13 | HiC_scaffold_11 | CP |
| Adductor Mandibulae Height | siglec14 | HiC_scaffold_11 | CP |
| Adductor Mandibulae Height | siglec9 | HiC_scaffold_11 | CP |
| Adductor Mandibulae Height | ski | HiC_scaffold_11 | CP |
| Adductor Mandibulae Height | slc22a17 | HiC_scaffold_11 | CP |
| Adductor Mandibulae Height | slc35b3 | HiC_scaffold_11 | CP |
| Adductor Mandibulae Height | snx16 | HiC_scaffold_11 | CP |
| Adductor Mandibulae Height | tecr | HiC_scaffold_11 | CP |
| Adductor Mandibulae Height | tfap2a | HiC_scaffold_11 | CP |
| Adductor Mandibulae Height | thap6 | HiC_scaffold_11 | CP |
| Adductor Mandibulae Height | thtpa | HiC_scaffold_11 | CP |

| | | | |
|---|---|---|---|
| Adductor Mandibulae Height | tm9sf1 | HiC_scaffold_11 | CP |
| Adductor Mandibulae Height | tmem14c | HiC_scaffold_11 | CP |
| Adductor Mandibulae Height | tmem51 | HiC_scaffold_11 | CP |
| Adductor Mandibulae Height | tmem56-b | HiC_scaffold_11 | CP |
| Adductor Mandibulae Height | txndc5 | HiC_scaffold_11 | CP |
| Adductor Mandibulae Height | utp3 | HiC_scaffold_11 | CP |
| Adductor Mandibulae Height | vav3 | HiC_scaffold_11 | CP |
| Adductor Mandibulae Height | wdr19 | HiC_scaffold_11 | CP |
| Adductor Mandibulae Height | wdr37 | HiC_scaffold_11 | CP |
| Adductor Mandibulae Height | wwp1 | HiC_scaffold_11 | CP |
| Adductor Mandibulae Height | zc2hc1a | HiC_scaffold_11 | CP |
| Adductor Mandibulae Height | zfhx4 | HiC_scaffold_11 | CP |
| Cranial Height | aak1 | HiC_scaffold_33 | CP |
| Cranial Height | acacb | HiC_scaffold_33 | CP |
| Cranial Height | acdh-11 | HiC_scaffold_33 | CP |
| Cranial Height | adra1a | HiC_scaffold_33 | CP |
| Cranial Height | adrb4c | HiC_scaffold_33 | CP |
| Cranial Height | aebp1 | HiC_scaffold_33 | CP |
| Cranial Height | aifm3 | HiC_scaffold_33 | CP |
| Cranial Height | akr1a1a | HiC_scaffold_33 | CP |
| Cranial Height | aldh3b1 | HiC_scaffold_33 | CP |
| Cranial Height | amy2 | HiC_scaffold_33 | CP |
| Cranial Height | ank1 | HiC_scaffold_33 | CP |
| Cranial Height | ank1 | HiC_scaffold_33 | LL |
| Cranial Height | ankrd13a | HiC_scaffold_33 | CP |
| Cranial Height | ankrd39 | HiC_scaffold_33 | CP |
| Cranial Height | antxr1 | HiC_scaffold_33 | CP |
| Cranial Height | aopep | HiC_scaffold_39 | CP |
| Cranial Height | ap1b1 | HiC_scaffold_33 | CP |
| Cranial Height | ap1b1 | HiC_scaffold_33 | LL |
| Cranial Height | aqp3 | HiC_scaffold_33 | CP |
| Cranial Height | arl6ip4 | HiC_scaffold_33 | CP |
| Cranial Height | arl6ip4 | HiC_scaffold_33 | LL |
| Cranial Height | arrdc3 | HiC_scaffold_33 | CP |
| Cranial Height | arsk | HiC_scaffold_33 | CP |
| Cranial Height | asb6 | HiC_scaffold_33 | CP |
| Cranial Height | atp6v0a2 | HiC_scaffold_33 | CP |
| Cranial Height | atp6v0a2 | HiC_scaffold_33 | LL |
| Cranial Height | bag4 | HiC_scaffold_33 | CP |
| Cranial Height | bco2 | HiC_scaffold_33 | CP |
| Cranial Height | bin3 | HiC_scaffold_33 | CP |

| | | | |
|---|---|---|---|
| Cranial Height | bmp1 | HiC_scaffold_33 | CP |
| Cranial Height | bri3bp | HiC_scaffold_33 | CP |
| Cranial Height | bri3bp | HiC_scaffold_33 | LL |
| Cranial Height | c2cd4cc2cd4_family | HiC_scaffold_33 | CP |
| Cranial Height | c2cd4cc2cd4_family | HiC_scaffold_33 | LL |
| Cranial Height | c9orf78 | HiC_scaffold_33 | CP |
| Cranial Height | carnmt1 | HiC_scaffold_33 | CP |
| Cranial Height | ccdc117 | HiC_scaffold_33 | CP |
| Cranial Height | ccdc117 | HiC_scaffold_33 | LL |
| Cranial Height | ccdc157 | HiC_scaffold_33 | CP |
| Cranial Height | ccdc92 | HiC_scaffold_33 | CP |
| Cranial Height | ccnh | HiC_scaffold_33 | CP |
| Cranial Height | cemip2 | HiC_scaffold_33 | CP |
| Cranial Height | ciao1a | HiC_scaffold_33 | CP |
| Cranial Height | cldn22 | HiC_scaffold_33 | CP |
| Cranial Height | clip1 | HiC_scaffold_33 | CP |
| Cranial Height | cmlkr1 | HiC_scaffold_33 | CP |
| Cranial Height | coe2 | HiC_scaffold_33 | CP |
| Cranial Height | coe2 | HiC_scaffold_33 | LL |
| Cranial Height | cox7c | HiC_scaffold_33 | CP |
| Cranial Height | crat | HiC_scaffold_33 | CP |
| Cranial Height | ctrc | HiC_scaffold_33 | CP |
| Cranial Height | dao | HiC_scaffold_33 | CP |
| Cranial Height | dbn1 | HiC_scaffold_33 | CP |
| Cranial Height | dbnl | HiC_scaffold_33 | CP |
| Cranial Height | ddr2 | HiC_scaffold_33 | CP |
| Cranial Height | dguok | HiC_scaffold_33 | CP |
| Cranial Height | dguok | HiC_scaffold_33 | LL |
| Cranial Height | disp3 | HiC_scaffold_33 | CP |
| Cranial Height | dnajb5 | HiC_scaffold_33 | CP |
| Cranial Height | dnm1l | HiC_scaffold_33 | CP |
| Cranial Height | dpysl2 | HiC_scaffold_33 | CP |
| Cranial Height | dusp18 | HiC_scaffold_33 | CP |
| Cranial Height | dusp26 | HiC_scaffold_33 | CP |
| Cranial Height | ehd1 | HiC_scaffold_33 | CP |
| Cranial Height | eif4ebp1 | HiC_scaffold_33 | CP |
| Cranial Height | eif4ebp1 | HiC_scaffold_33 | LL |
| Cranial Height | elac1 | HiC_scaffold_39 | CP |
| Cranial Height | elmod3 | HiC_scaffold_33 | CP |
| Cranial Height | elovl7 | HiC_scaffold_39 | CP |
| Cranial Height | emid1 | HiC_scaffold_33 | CP |

| | | | |
|---|---|---|---|
| Cranial Height | emid1 | HiC_scaffold_33 | LL |
| Cranial Height | epx | HiC_scaffold_33 | CP |
| Cranial Height | epx | HiC_scaffold_33 | LL |
| Cranial Height | erap2 | HiC_scaffold_33 | CP |
| Cranial Height | ercc8 | HiC_scaffold_39 | CP |
| Cranial Height | es1 | HiC_scaffold_33 | CP |
| Cranial Height | ewsr1 | HiC_scaffold_33 | CP |
| Cranial Height | ewsr1 | HiC_scaffold_33 | LL |
| Cranial Height | f2r | HiC_scaffold_33 | CP |
| Cranial Height | fabp1 | HiC_scaffold_39 | CP |
| Cranial Height | fam173b | HiC_scaffold_39 | CP |
| Cranial Height | fam219a | HiC_scaffold_33 | CP |
| Cranial Height | fam222a | HiC_scaffold_33 | CP |
| Cranial Height | fancg | HiC_scaffold_33 | CP |
| Cranial Height | fgfr1a | HiC_scaffold_33 | CP |
| Cranial Height | ficd | HiC_scaffold_33 | CP |
| Cranial Height | foxb2 | HiC_scaffold_33 | CP |
| Cranial Height | foxb2 | HiC_scaffold_33 | LL |
| Cranial Height | foxd5-a | HiC_scaffold_39 | CP |
| Cranial Height | foxn4 | HiC_scaffold_33 | CP |
| Cranial Height | fzd10-a | HiC_scaffold_33 | CP |
| Cranial Height | gal3st1 | HiC_scaffold_33 | CP |
| Cranial Height | gas2l1 | HiC_scaffold_33 | CP |
| Cranial Height | gas2l1 | HiC_scaffold_33 | LL |
| Cranial Height | gatc | HiC_scaffold_33 | CP |
| Cranial Height | gcnt1 | HiC_scaffold_33 | CP |
| Cranial Height | gfra2 | HiC_scaffold_33 | CP |
| Cranial Height | gimap2 | HiC_scaffold_33 | CP |
| Cranial Height | gimap4 | HiC_scaffold_33 | CP |
| Cranial Height | gimap7 | HiC_scaffold_33 | CP |
| Cranial Height | gimap8 | HiC_scaffold_33 | CP |
| Cranial Height | gins4 | HiC_scaffold_33 | CP |
| Cranial Height | git2 | HiC_scaffold_33 | CP |
| Cranial Height | gltp | HiC_scaffold_33 | CP |
| Cranial Height | gna14 | HiC_scaffold_33 | CP |
| Cranial Height | gna14 | HiC_scaffold_33 | LL |
| Cranial Height | gnaq | HiC_scaffold_33 | CP |
| Cranial Height | gnaq | HiC_scaffold_33 | LL |
| Cranial Height | gnrh1 | HiC_scaffold_33 | CP |
| Cranial Height | gpat4 | HiC_scaffold_33 | CP |
| Cranial Height | gramd2b | HiC_scaffold_33 | CP |

| | | | |
|---|---|---|---|
| Cranial Height | grk5 | HiC_scaffold_33 | CP |
| Cranial Height | gtf2h3 | HiC_scaffold_33 | CP |
| Cranial Height | hapln1 | HiC_scaffold_33 | CP |
| Cranial Height | hcar2 | HiC_scaffold_33 | CP |
| Cranial Height | hcar2 | HiC_scaffold_33 | LL |
| Cranial Height | hip1r | HiC_scaffold_33 | CP |
| Cranial Height | hip1r | HiC_scaffold_33 | LL |
| Cranial Height | homer1 | HiC_scaffold_33 | CP |
| Cranial Height | hspb11 | HiC_scaffold_33 | CP |
| Cranial Height | ier5l | HiC_scaffold_33 | CP |
| Cranial Height | igsf9 | HiC_scaffold_33 | CP |
| Cranial Height | ine | HiC_scaffold_33 | CP |
| Cranial Height | ine | HiC_scaffold_33 | LL |
| Cranial Height | iqgap1 | HiC_scaffold_33 | CP |
| Cranial Height | iscu | HiC_scaffold_33 | CP |
| Cranial Height | jmy | HiC_scaffold_33 | CP |
| Cranial Height | kansl3 | HiC_scaffold_33 | CP |
| Cranial Height | kctd10 | HiC_scaffold_33 | CP |
| Cranial Height | kctd9 | HiC_scaffold_33 | CP |
| Cranial Height | kiaa0825 | HiC_scaffold_33 | CP |
| Cranial Height | kif24 | HiC_scaffold_33 | CP |
| Cranial Height | kin14e | HiC_scaffold_33 | CP |
| Cranial Height | klf9 | HiC_scaffold_33 | CP |
| Cranial Height | klhl10 | HiC_scaffold_33 | CP |
| Cranial Height | klhl10 | HiC_scaffold_33 | LL |
| Cranial Height | kmt5aa | HiC_scaffold_33 | CP |
| Cranial Height | kmt5aa | HiC_scaffold_33 | LL |
| Cranial Height | kntc1 | HiC_scaffold_33 | CP |
| Cranial Height | koza | HiC_scaffold_33 | CP |
| Cranial Height | lgi3 | HiC_scaffold_33 | CP |
| Cranial Height | limk2 | HiC_scaffold_33 | CP |
| Cranial Height | limk2 | HiC_scaffold_33 | LL |
| Cranial Height | lix1 | HiC_scaffold_33 | CP |
| Cranial Height | lox | HiC_scaffold_33 | CP |
| Cranial Height | loxhd1 | HiC_scaffold_39 | CP |
| Cranial Height | lsm1 | HiC_scaffold_33 | CP |
| Cranial Height | lysmd3 | HiC_scaffold_33 | CP |
| Cranial Height | lztr1 | HiC_scaffold_33 | CP |
| Cranial Height | lzts1 | HiC_scaffold_33 | CP |
| Cranial Height | mapk4 | HiC_scaffold_39 | CP |
| Cranial Height | mblac2 | HiC_scaffold_33 | CP |

| | | | |
|---|---|---|---|
| Cranial Height | mctp1 | HiC_scaffold_33 | CP |
| Cranial Height | me2 | HiC_scaffold_39 | CP |
| Cranial Height | med22 | HiC_scaffold_33 | CP |
| Cranial Height | mfsd3 | HiC_scaffold_33 | CP |
| Cranial Height | mier3 | HiC_scaffold_39 | CP |
| Cranial Height | mlxip | HiC_scaffold_33 | CP |
| Cranial Height | mmab | HiC_scaffold_33 | CP |
| Cranial Height | mn1 | HiC_scaffold_33 | CP |
| Cranial Height | mob1a | HiC_scaffold_33 | CP |
| Cranial Height | mob1a | HiC_scaffold_33 | LL |
| Cranial Height | morc2a | HiC_scaffold_33 | CP |
| Cranial Height | mspa | HiC_scaffold_33 | CP |
| Cranial Height | mthfd2 | HiC_scaffold_33 | CP |
| Cranial Height | mthfd2 | HiC_scaffold_33 | LL |
| Cranial Height | mtx3 | HiC_scaffold_33 | CP |
| Cranial Height | mvk | HiC_scaffold_33 | CP |
| Cranial Height | myo1h | HiC_scaffold_33 | CP |
| Cranial Height | ncs1 | HiC_scaffold_33 | CP |
| Cranial Height | ndufaf2 | HiC_scaffold_39 | CP |
| Cranial Height | nefh | HiC_scaffold_33 | CP |
| Cranial Height | nefl | HiC_scaffold_33 | CP |
| Cranial Height | neurl3 | HiC_scaffold_33 | CP |
| Cranial Height | nfu1 | HiC_scaffold_33 | CP |
| Cranial Height | nkx2-6 | HiC_scaffold_33 | CP |
| Cranial Height | nkx6-3 | HiC_scaffold_33 | CP |
| Cranial Height | nodal | HiC_scaffold_33 | CP |
| Cranial Height | nodal | HiC_scaffold_33 | LL |
| Cranial Height | nol6 | HiC_scaffold_33 | CP |
| Cranial Height | nono | HiC_scaffold_33 | CP |
| Cranial Height | npc1l1 | HiC_scaffold_33 | CP |
| Cranial Height | npy1r | HiC_scaffold_33 | CP |
| Cranial Height | omg | HiC_scaffold_33 | CP |
| Cranial Height | osbp2 | HiC_scaffold_33 | CP |
| Cranial Height | osbp2 | HiC_scaffold_33 | LL |
| Cranial Height | p2rx2 | HiC_scaffold_33 | CP |
| Cranial Height | p2ry14 | HiC_scaffold_33 | CP |
| Cranial Height | p2ry14 | HiC_scaffold_33 | LL |
| Cranial Height | pcsk5 | HiC_scaffold_33 | CP |
| Cranial Height | pde4d | HiC_scaffold_39 | CP |
| Cranial Height | pdlim2 | HiC_scaffold_33 | CP |
| Cranial Height | pebp4 | HiC_scaffold_33 | CP |

| | | | |
|---|---|---|---|
| Cranial Height | pes1 | HiC_scaffold_33 | CP |
| Cranial Height | pgbd3 | HiC_scaffold_33 | CP |
| Cranial Height | pgbd3 | HiC_scaffold_33 | LL |
| Cranial Height | pgm5 | HiC_scaffold_39 | CP |
| Cranial Height | phyhip | HiC_scaffold_33 | CP |
| Cranial Height | pik3ip1 | HiC_scaffold_33 | CP |
| Cranial Height | pik3ip1 | HiC_scaffold_33 | LL |
| Cranial Height | pip5k1b | HiC_scaffold_33 | CP |
| Cranial Height | pitpnb | HiC_scaffold_33 | CP |
| Cranial Height | pitpnm2 | HiC_scaffold_33 | CP |
| Cranial Height | pitpnm2 | HiC_scaffold_33 | LL |
| Cranial Height | pla2g3 | HiC_scaffold_33 | CP |
| Cranial Height | plcl1 | HiC_scaffold_33 | CP |
| Cranial Height | plekha2 | HiC_scaffold_33 | CP |
| Cranial Height | plk2 | HiC_scaffold_39 | CP |
| Cranial Height | pole | HiC_scaffold_33 | CP |
| Cranial Height | polr3d | HiC_scaffold_33 | CP |
| Cranial Height | polr3g | HiC_scaffold_33 | CP |
| Cranial Height | ppp1r3c | HiC_scaffold_33 | CP |
| Cranial Height | prlhr | HiC_scaffold_33 | CP |
| Cranial Height | prune2 | HiC_scaffold_33 | CP |
| Cranial Height | prune2 | HiC_scaffold_33 | LL |
| Cranial Height | psap | HiC_scaffold_33 | CP |
| Cranial Height | psap | HiC_scaffold_33 | LL |
| Cranial Height | psbp1 | HiC_scaffold_33 | CP |
| Cranial Height | psmd9 | HiC_scaffold_33 | CP |
| Cranial Height | ptch1 | HiC_scaffold_39 | CP |
| Cranial Height | ptger4 | HiC_scaffold_39 | CP |
| Cranial Height | pxmp2 | HiC_scaffold_33 | CP |
| Cranial Height | rab11fip1 | HiC_scaffold_33 | CP |
| Cranial Height | rab3c | HiC_scaffold_39 | CP |
| Cranial Height | rabgef1 | HiC_scaffold_33 | CP |
| Cranial Height | rasa1 | HiC_scaffold_33 | CP |
| Cranial Height | rasl10b | HiC_scaffold_33 | CP |
| Cranial Height | rasl10b | HiC_scaffold_33 | LL |
| Cranial Height | rfesd | HiC_scaffold_33 | CP |
| Cranial Height | rflna | HiC_scaffold_33 | CP |
| Cranial Height | rgmb | HiC_scaffold_33 | CP |
| Cranial Height | rhbdd3 | HiC_scaffold_33 | CP |
| Cranial Height | rhbdd3 | HiC_scaffold_33 | LL |
| Cranial Height | rilpl1 | HiC_scaffold_33 | CP |

| | | | |
|---|---|---|---|
| Cranial Height | rilpl1 | HiC_scaffold_33 | LL |
| Cranial Height | rilpl2 | HiC_scaffold_33 | CP |
| Cranial Height | rilpl2 | HiC_scaffold_33 | LL |
| Cranial Height | rimbp2 | HiC_scaffold_33 | CP |
| Cranial Height | riok2 | HiC_scaffold_33 | CP |
| Cranial Height | rnf214 | HiC_scaffold_33 | CP |
| Cranial Height | rnf223 | HiC_scaffold_33 | CP |
| Cranial Height | rorb | HiC_scaffold_33 | CP |
| Cranial Height | rph3a | HiC_scaffold_33 | CP |
| Cranial Height | rsrc2 | HiC_scaffold_33 | CP |
| Cranial Height | rtkn | HiC_scaffold_33 | CP |
| Cranial Height | sart3 | HiC_scaffold_33 | CP |
| Cranial Height | sds | HiC_scaffold_33 | CP |
| Cranial Height | seca | HiC_scaffold_33 | CP |
| Cranial Height | setbp1 | HiC_scaffold_33 | CP |
| Cranial Height | sgsm1 | HiC_scaffold_33 | CP |
| Cranial Height | slc15a4 | HiC_scaffold_33 | CP |
| Cranial Height | slc25a37 | HiC_scaffold_33 | CP |
| Cranial Height | slc2a11 | HiC_scaffold_33 | CP |
| Cranial Height | slc2a8 | HiC_scaffold_33 | CP |
| Cranial Height | slc6a4 | HiC_scaffold_33 | CP |
| Cranial Height | slc7a4 | HiC_scaffold_33 | CP |
| Cranial Height | slc8b1 | HiC_scaffold_33 | CP |
| Cranial Height | slf1 | HiC_scaffold_33 | CP |
| Cranial Height | smim15 | HiC_scaffold_39 | CP |
| Cranial Height | smn1 | HiC_scaffold_33 | CP |
| Cranial Height | smtn | HiC_scaffold_33 | CP |
| Cranial Height | smtn | HiC_scaffold_33 | LL |
| Cranial Height | smtnl1 | HiC_scaffold_33 | CP |
| Cranial Height | smtnl1 | HiC_scaffold_33 | LL |
| Cranial Height | smyd1 | HiC_scaffold_39 | CP |
| Cranial Height | snrnp200 | HiC_scaffold_33 | CP |
| Cranial Height | snx2 | HiC_scaffold_33 | CP |
| Cranial Height | sorbs3 | HiC_scaffold_33 | CP |
| Cranial Height | srrd | HiC_scaffold_33 | CP |
| Cranial Height | ssbp2 | HiC_scaffold_33 | CP |
| Cranial Height | ssh1 | HiC_scaffold_33 | CP |
| Cranial Height | star | HiC_scaffold_33 | CP |
| Cranial Height | stx2 | HiC_scaffold_33 | CP |
| Cranial Height | sv2c | HiC_scaffold_33 | CP |
| Cranial Height | svop | HiC_scaffold_33 | CP |

| | | | |
|---|---|---|---|
| Cranial Height | tacc1 | HiC_scaffold_33 | CP |
| Cranial Height | tbx5 | HiC_scaffold_33 | CP |
| Cranial Height | tcf7l1a | HiC_scaffold_33 | CP |
| Cranial Height | tchp | HiC_scaffold_33 | CP |
| Cranial Height | tcn2 | HiC_scaffold_33 | CP |
| Cranial Height | tcn2 | HiC_scaffold_33 | LL |
| Cranial Height | tctn2 | HiC_scaffold_33 | CP |
| Cranial Height | tctn2 | HiC_scaffold_33 | LL |
| Cranial Height | tent2 | HiC_scaffold_33 | CP |
| Cranial Height | tfip11 | HiC_scaffold_33 | CP |
| Cranial Height | thbs4b | HiC_scaffold_33 | CP |
| Cranial Height | thoc5 | HiC_scaffold_33 | CP |
| Cranial Height | tmem119 | HiC_scaffold_33 | CP |
| Cranial Height | tmem127 | HiC_scaffold_33 | CP |
| Cranial Height | tmem132c | HiC_scaffold_33 | CP |
| Cranial Height | tmem132d | HiC_scaffold_33 | CP |
| Cranial Height | tmem161b | HiC_scaffold_33 | CP |
| Cranial Height | tmem167a | HiC_scaffold_33 | CP |
| Cranial Height | tmem230 | HiC_scaffold_33 | CP |
| Cranial Height | tmem248 | HiC_scaffold_33 | CP |
| Cranial Height | tnks | HiC_scaffold_33 | CP |
| Cranial Height | tpcn1 | HiC_scaffold_33 | CP |
| Cranial Height | trafd1 | HiC_scaffold_33 | CP |
| Cranial Height | trafd1 | HiC_scaffold_33 | LL |
| Cranial Height | trpm3 | HiC_scaffold_33 | CP |
| Cranial Height | trpm6 | HiC_scaffold_33 | CP |
| Cranial Height | tspan36 | HiC_scaffold_39 | CP |
| Cranial Height | ttc28 | HiC_scaffold_33 | CP |
| Cranial Height | ttc37 | HiC_scaffold_33 | CP |
| Cranial Height | tutl | HiC_scaffold_33 | CP |
| Cranial Height | ube3b | HiC_scaffold_33 | CP |
| Cranial Height | ubl4aa | HiC_scaffold_33 | CP |
| Cranial Height | ulk1 | HiC_scaffold_33 | CP |
| Cranial Height | unc45b | HiC_scaffold_33 | CP |
| Cranial Height | usp30 | HiC_scaffold_33 | CP |
| Cranial Height | usp39 | HiC_scaffold_33 | CP |
| Cranial Height | usp39 | HiC_scaffold_33 | LL |
| Cranial Height | vcp | HiC_scaffold_33 | CP |
| Cranial Height | vegt | HiC_scaffold_33 | CP |
| Cranial Height | vps13c | HiC_scaffold_33 | CP |
| Cranial Height | vps33a | HiC_scaffold_33 | CP |

| | | | |
|---|---|---|---|
| Cranial Height | wdr31 | HiC_scaffold_33 | CP |
| Cranial Height | wdr31 | HiC_scaffold_33 | LL |
| Cranial Height | wdr66 | HiC_scaffold_33 | CP |
| Cranial Height | wscd2 | HiC_scaffold_33 | CP |
| Cranial Height | xrcc4 | HiC_scaffold_33 | CP |
| Cranial Height | zfand5 | HiC_scaffold_33 | CP |
| Cranial Height | znf180 | HiC_scaffold_33 | CP |
| Cranial Height | znf366 | HiC_scaffold_39 | CP |
| Cranial Height | znf608 | HiC_scaffold_33 | CP |
| Dentigerous Arm Depth | abca4 | HiC_scaffold_11 | CP |
| Dentigerous Arm Depth | abhd10 | HiC_scaffold_11 | CP |
| Dentigerous Arm Depth | abi1 | HiC_scaffold_11 | CP |
| Dentigerous Arm Depth | acad11 | HiC_scaffold_11 | CP |
| Dentigerous Arm Depth | acbd5a | HiC_scaffold_11 | CP |
| Dentigerous Arm Depth | ackr4 | HiC_scaffold_11 | CP |
| Dentigerous Arm Depth | adamts12 | HiC_scaffold_5 | LL |
| Dentigerous Arm Depth | adamts7 | HiC_scaffold_5 | LL |
| Dentigerous Arm Depth | adgrg4 | HiC_scaffold_11 | CP |
| Dentigerous Arm Depth | agrp | HiC_scaffold_11 | CP |
| Dentigerous Arm Depth | agtr1 | HiC_scaffold_11 | CP |
| Dentigerous Arm Depth | ahrr | HiC_scaffold_11 | CP |
| Dentigerous Arm Depth | akap13 | HiC_scaffold_5 | LL |
| Dentigerous Arm Depth | amer2 | HiC_scaffold_11 | CP |
| Dentigerous Arm Depth | ankh | HiC_scaffold_11 | CP |
| Dentigerous Arm Depth | ankrd33b | HiC_scaffold_11 | CP |
| Dentigerous Arm Depth | ano9 | HiC_scaffold_5 | LL |
| Dentigerous Arm Depth | ap2a2 | HiC_scaffold_5 | LL |
| Dentigerous Arm Depth | apod | HiC_scaffold_11 | CP |
| Dentigerous Arm Depth | arfgef1 | HiC_scaffold_11 | CP |
| Dentigerous Arm Depth | arhgap21 | HiC_scaffold_11 | CP |
| Dentigerous Arm Depth | arhgap29 | HiC_scaffold_11 | CP |
| Dentigerous Arm Depth | armc1 | HiC_scaffold_11 | CP |
| Dentigerous Arm Depth | arpp19 | HiC_scaffold_5 | LL |
| Dentigerous Arm Depth | arx | HiC_scaffold_11 | CP |
| Dentigerous Arm Depth | asap1 | HiC_scaffold_11 | CP |
| Dentigerous Arm Depth | atp8a2 | HiC_scaffold_11 | CP |
| Dentigerous Arm Depth | b2m | HiC_scaffold_11 | CP |
| Dentigerous Arm Depth | b4galt1 | HiC_scaffold_11 | CP |
| Dentigerous Arm Depth | bbs4 | HiC_scaffold_5 | LL |
| Dentigerous Arm Depth | bco1 | HiC_scaffold_11 | CP |
| Dentigerous Arm Depth | bdh1 | HiC_scaffold_11 | CP |

| | | | |
|---|---|---|---|
| Dentigerous Arm Depth | bhlhe22 | HiC_scaffold_11 | CP |
| Dentigerous Arm Depth | bloc1s5 | HiC_scaffold_11 | CP |
| Dentigerous Arm Depth | boc | HiC_scaffold_11 | CP |
| Dentigerous Arm Depth | brpf3 | HiC_scaffold_11 | CP |
| Dentigerous Arm Depth | c1qtnf9 | HiC_scaffold_11 | CP |
| Dentigerous Arm Depth | c8g | HiC_scaffold_11 | CP |
| Dentigerous Arm Depth | ca1 | HiC_scaffold_11 | CP |
| Dentigerous Arm Depth | cacnb2 | HiC_scaffold_11 | CP |
| Dentigerous Arm Depth | calml4 | HiC_scaffold_5 | LL |
| Dentigerous Arm Depth | caprin2 | HiC_scaffold_11 | CP |
| Dentigerous Arm Depth | cbln2 | HiC_scaffold_11 | CP |
| Dentigerous Arm Depth | ccdc106 | HiC_scaffold_11 | CP |
| Dentigerous Arm Depth | ccdc58 | HiC_scaffold_11 | CP |
| Dentigerous Arm Depth | ccdc63 | HiC_scaffold_11 | CP |
| Dentigerous Arm Depth | ccl20 | HiC_scaffold_11 | CP |
| Dentigerous Arm Depth | ccne1 | HiC_scaffold_5 | LL |
| Dentigerous Arm Depth | ccr1 | HiC_scaffold_11 | CP |
| Dentigerous Arm Depth | cct5 | HiC_scaffold_11 | CP |
| Dentigerous Arm Depth | cd226 | HiC_scaffold_11 | CP |
| Dentigerous Arm Depth | cd276 | HiC_scaffold_5 | LL |
| Dentigerous Arm Depth | cd38 | HiC_scaffold_11 | CP |
| Dentigerous Arm Depth | cd81 | HiC_scaffold_5 | LL |
| Dentigerous Arm Depth | cdh10 | HiC_scaffold_11 | CP |
| Dentigerous Arm Depth | cdh12 | HiC_scaffold_11 | CP |
| Dentigerous Arm Depth | cdh18 | HiC_scaffold_11 | CP |
| Dentigerous Arm Depth | cdh20 | HiC_scaffold_11 | CP |
| Dentigerous Arm Depth | cdh6 | HiC_scaffold_11 | CP |
| Dentigerous Arm Depth | cdh7 | HiC_scaffold_11 | CP |
| Dentigerous Arm Depth | cdk13 | HiC_scaffold_11 | CP |
| Dentigerous Arm Depth | cdk8 | HiC_scaffold_11 | CP |
| Dentigerous Arm Depth | cdv3 | HiC_scaffold_11 | CP |
| Dentigerous Arm Depth | cebpe | HiC_scaffold_11 | CP |
| Dentigerous Arm Depth | cela2a | HiC_scaffold_11 | CP |
| Dentigerous Arm Depth | chmp4c | HiC_scaffold_11 | CP |
| Dentigerous Arm Depth | chmp5 | HiC_scaffold_11 | CP |
| Dentigerous Arm Depth | chrna7 | HiC_scaffold_5 | LL |
| Dentigerous Arm Depth | chst2 | HiC_scaffold_11 | CP |
| Dentigerous Arm Depth | cldn15 | HiC_scaffold_5 | LL |
| Dentigerous Arm Depth | cln6 | HiC_scaffold_5 | LL |
| Dentigerous Arm Depth | clul1 | HiC_scaffold_11 | CP |
| Dentigerous Arm Depth | cmbl | HiC_scaffold_11 | CP |

| | | | |
|---|---|---|---|
| Dentigerous Arm Depth | cngb3 | HiC_scaffold_11 | CP |
| Dentigerous Arm Depth | col11a1 | HiC_scaffold_11 | CP |
| Dentigerous Arm Depth | colec12 | HiC_scaffold_11 | CP |
| Dentigerous Arm Depth | cops5 | HiC_scaffold_11 | CP |
| Dentigerous Arm Depth | cpa6 | HiC_scaffold_11 | CP |
| Dentigerous Arm Depth | cpb1 | HiC_scaffold_11 | CP |
| Dentigerous Arm Depth | cpeb1 | HiC_scaffold_5 | LL |
| Dentigerous Arm Depth | cpne3 | HiC_scaffold_11 | CP |
| Dentigerous Arm Depth | crh | HiC_scaffold_11 | CP |
| Dentigerous Arm Depth | crispld1 | HiC_scaffold_11 | CP |
| Dentigerous Arm Depth | cry-dash | HiC_scaffold_11 | CP |
| Dentigerous Arm Depth | csnk1g1 | HiC_scaffold_5 | LL |
| Dentigerous Arm Depth | csrnp1 | HiC_scaffold_11 | CP |
| Dentigerous Arm Depth | cstb | HiC_scaffold_11 | CP |
| Dentigerous Arm Depth | dcaf11 | HiC_scaffold_11 | CP |
| Dentigerous Arm Depth | dhcr7 | HiC_scaffold_5 | LL |
| Dentigerous Arm Depth | dlec1 | HiC_scaffold_11 | CP |
| Dentigerous Arm Depth | dnajb6 | HiC_scaffold_11 | CP |
| Dentigerous Arm Depth | dnajc13 | HiC_scaffold_11 | CP |
| Dentigerous Arm Depth | dok6 | HiC_scaffold_11 | CP |
| Dentigerous Arm Depth | dph2 | HiC_scaffold_11 | CP |
| Dentigerous Arm Depth | dpp6 | HiC_scaffold_11 | CP |
| Dentigerous Arm Depth | drd3 | HiC_scaffold_11 | CP |
| Dentigerous Arm Depth | drosha | HiC_scaffold_11 | CP |
| Dentigerous Arm Depth | dsel | HiC_scaffold_11 | CP |
| Dentigerous Arm Depth | dusp28 | HiC_scaffold_5 | LL |
| Dentigerous Arm Depth | eef1e1 | HiC_scaffold_11 | CP |
| Dentigerous Arm Depth | ell2 | HiC_scaffold_5 | LL |
| Dentigerous Arm Depth | eloc | HiC_scaffold_11 | CP |
| Dentigerous Arm Depth | emc9 | HiC_scaffold_11 | CP |
| Dentigerous Arm Depth | emilin2 | HiC_scaffold_11 | CP |
| Dentigerous Arm Depth | emp3 | HiC_scaffold_11 | CP |
| Dentigerous Arm Depth | erya | HiC_scaffold_11 | CP |
| Dentigerous Arm Depth | esyt2 | HiC_scaffold_11 | CP |
| Dentigerous Arm Depth | eya1 | HiC_scaffold_11 | CP |
| Dentigerous Arm Depth | f13a1 | HiC_scaffold_11 | CP |
| Dentigerous Arm Depth | f13e9.13 | HiC_scaffold_5 | LL |
| Dentigerous Arm Depth | fam168b | HiC_scaffold_11 | CP |
| Dentigerous Arm Depth | fam214a | HiC_scaffold_5 | LL |
| Dentigerous Arm Depth | fam49b | HiC_scaffold_11 | CP |
| Dentigerous Arm Depth | fam83e | HiC_scaffold_11 | CP |

| | | | |
|---|---|---|---|
| Dentigerous Arm Depth | fastkd3 | HiC_scaffold_11 | CP |
| Dentigerous Arm Depth | fbxl7 | HiC_scaffold_11 | CP |
| Dentigerous Arm Depth | fen1 | HiC_scaffold_11 | CP |
| Dentigerous Arm Depth | fitm1 | HiC_scaffold_11 | CP |
| Dentigerous Arm Depth | flt3 | HiC_scaffold_11 | CP |
| Dentigerous Arm Depth | foxh1 | HiC_scaffold_11 | CP |
| Dentigerous Arm Depth | foxj3 | HiC_scaffold_11 | CP |
| Dentigerous Arm Depth | gabarapl2 | HiC_scaffold_5 | LL |
| Dentigerous Arm Depth | gad2 | HiC_scaffold_11 | CP |
| Dentigerous Arm Depth | garem1 | HiC_scaffold_11 | CP |
| Dentigerous Arm Depth | gars | HiC_scaffold_11 | CP |
| Dentigerous Arm Depth | gdap1 | HiC_scaffold_11 | CP |
| Dentigerous Arm Depth | ggh | HiC_scaffold_11 | CP |
| Dentigerous Arm Depth | gimap4 | HiC_scaffold_11 | CP |
| Dentigerous Arm Depth | gli3 | HiC_scaffold_11 | CP |
| Dentigerous Arm Depth | glyctk | HiC_scaffold_11 | CP |
| Dentigerous Arm Depth | gnb5b | HiC_scaffold_5 | LL |
| Dentigerous Arm Depth | gnrhr2 | HiC_scaffold_5 | LL |
| Dentigerous Arm Depth | gorasp1 | HiC_scaffold_11 | CP |
| Dentigerous Arm Depth | gpr12 | HiC_scaffold_11 | CP |
| Dentigerous Arm Depth | gpr141 | HiC_scaffold_11 | CP |
| Dentigerous Arm Depth | gpr17 | HiC_scaffold_11 | CP |
| Dentigerous Arm Depth | gpt2l | HiC_scaffold_11 | CP |
| Dentigerous Arm Depth | gramd1c | HiC_scaffold_11 | CP |
| Dentigerous Arm Depth | gramd2a | HiC_scaffold_5 | LL |
| Dentigerous Arm Depth | gtf3a | HiC_scaffold_11 | CP |
| Dentigerous Arm Depth | hacd1 | HiC_scaffold_11 | CP |
| Dentigerous Arm Depth | hbp1 | HiC_scaffold_11 | CP |
| Dentigerous Arm Depth | hgd | HiC_scaffold_11 | CP |
| Dentigerous Arm Depth | hhatl | HiC_scaffold_11 | CP |
| Dentigerous Arm Depth | hmcn2 | HiC_scaffold_11 | CP |
| Dentigerous Arm Depth | hnf4g | HiC_scaffold_11 | CP |
| Dentigerous Arm Depth | idh2 | HiC_scaffold_5 | LL |
| Dentigerous Arm Depth | il20rb | HiC_scaffold_5 | LL |
| Dentigerous Arm Depth | impa1 | HiC_scaffold_11 | CP |
| Dentigerous Arm Depth | insig1 | HiC_scaffold_11 | CP |
| Dentigerous Arm Depth | insy1 | HiC_scaffold_5 | LL |
| Dentigerous Arm Depth | itga11 | HiC_scaffold_5 | LL |
| Dentigerous Arm Depth | jph1 | HiC_scaffold_11 | CP |
| Dentigerous Arm Depth | kazn | HiC_scaffold_11 | CP |
| Dentigerous Arm Depth | kazna | HiC_scaffold_11 | CP |

| | | | |
|---|---|---|---|
| Dentigerous Arm Depth | kbp | HiC_scaffold_11 | CP |
| Dentigerous Arm Depth | kbtbd2 | HiC_scaffold_11 | CP |
| Dentigerous Arm Depth | kcnb2 | HiC_scaffold_11 | CP |
| Dentigerous Arm Depth | kif13b | HiC_scaffold_5 | LL |
| Dentigerous Arm Depth | kif20a | HiC_scaffold_11 | CP |
| Dentigerous Arm Depth | klhl40b | HiC_scaffold_11 | CP |
| Dentigerous Arm Depth | kpna1 | HiC_scaffold_11 | CP |
| Dentigerous Arm Depth | limd2 | HiC_scaffold_11 | CP |
| Dentigerous Arm Depth | lnx2 | HiC_scaffold_11 | CP |
| Dentigerous Arm Depth | loxl1 | HiC_scaffold_5 | LL |
| Dentigerous Arm Depth | lpin2 | HiC_scaffold_11 | CP |
| Dentigerous Arm Depth | lsm5 | HiC_scaffold_11 | CP |
| Dentigerous Arm Depth | lypla1 | HiC_scaffold_11 | CP |
| Dentigerous Arm Depth | lztfl1 | HiC_scaffold_11 | CP |
| Dentigerous Arm Depth | maf1 | HiC_scaffold_11 | CP |
| Dentigerous Arm Depth | mag | HiC_scaffold_11 | CP |
| Dentigerous Arm Depth | mak | HiC_scaffold_11 | CP |
| Dentigerous Arm Depth | map3k15 | HiC_scaffold_11 | CP |
| Dentigerous Arm Depth | mastl | HiC_scaffold_11 | CP |
| Dentigerous Arm Depth | mc4r | HiC_scaffold_11 | CP |
| Dentigerous Arm Depth | mcl1 | HiC_scaffold_5 | LL |
| Dentigerous Arm Depth | mcur1 | HiC_scaffold_11 | CP |
| Dentigerous Arm Depth | med1 | HiC_scaffold_11 | CP |
| Dentigerous Arm Depth | mesd | HiC_scaffold_5 | LL |
| Dentigerous Arm Depth | mettl4 | HiC_scaffold_11 | CP |
| Dentigerous Arm Depth | mllt10 | HiC_scaffold_11 | CP |
| Dentigerous Arm Depth | mon1b | HiC_scaffold_11 | CP |
| Dentigerous Arm Depth | mrc1 | HiC_scaffold_11 | CP |
| Dentigerous Arm Depth | mrpl15 | HiC_scaffold_11 | CP |
| Dentigerous Arm Depth | mrpl46 | HiC_scaffold_5 | LL |
| Dentigerous Arm Depth | msc | HiC_scaffold_11 | CP |
| Dentigerous Arm Depth | msrb2 | HiC_scaffold_11 | CP |
| Dentigerous Arm Depth | mtfr1 | HiC_scaffold_11 | CP |
| Dentigerous Arm Depth | mtmr6 | HiC_scaffold_11 | CP |
| Dentigerous Arm Depth | mtrr | HiC_scaffold_11 | CP |
| Dentigerous Arm Depth | mtss1l | HiC_scaffold_5 | LL |
| Dentigerous Arm Depth | mup20 | HiC_scaffold_11 | CP |
| Dentigerous Arm Depth | mybl1 | HiC_scaffold_11 | CP |
| Dentigerous Arm Depth | myd88 | HiC_scaffold_11 | CP |
| Dentigerous Arm Depth | myh6 | HiC_scaffold_11 | CP |
| Dentigerous Arm Depth | myh7 | HiC_scaffold_11 | CP |

| | | | |
|---|---|---|---|
| Dentigerous Arm Depth | myo5a | HiC_scaffold_5 | LL |
| Dentigerous Arm Depth | myo9a | HiC_scaffold_5 | LL |
| Dentigerous Arm Depth | naa50 | HiC_scaffold_11 | CP |
| Dentigerous Arm Depth | ncapg2 | HiC_scaffold_11 | CP |
| Dentigerous Arm Depth | nck1 | HiC_scaffold_11 | CP |
| Dentigerous Arm Depth | ncoa2 | HiC_scaffold_11 | CP |
| Dentigerous Arm Depth | neto1 | HiC_scaffold_11 | CP |
| Dentigerous Arm Depth | nfi1 | HiC_scaffold_11 | CP |
| Dentigerous Arm Depth | nfx1 | HiC_scaffold_11 | CP |
| Dentigerous Arm Depth | ngdn | HiC_scaffold_11 | CP |
| Dentigerous Arm Depth | nlrc3 | HiC_scaffold_11 | CP |
| Dentigerous Arm Depth | nlrp1 | HiC_scaffold_11 | CP |
| Dentigerous Arm Depth | nlrp12 | HiC_scaffold_5 | LL |
| Dentigerous Arm Depth | nom1 | HiC_scaffold_11 | CP |
| Dentigerous Arm Depth | nrn1 | HiC_scaffold_11 | CP |
| Dentigerous Arm Depth | nrros | HiC_scaffold_11 | CP |
| Dentigerous Arm Depth | ntng1 | HiC_scaffold_11 | CP |
| Dentigerous Arm Depth | ntrk3 | HiC_scaffold_5 | LL |
| Dentigerous Arm Depth | nup58 | HiC_scaffold_11 | CP |
| Dentigerous Arm Depth | olfm3 | HiC_scaffold_11 | CP |
| Dentigerous Arm Depth | onecut1 | HiC_scaffold_5 | LL |
| Dentigerous Arm Depth | oplah | HiC_scaffold_11 | CP |
| Dentigerous Arm Depth | oprk1 | HiC_scaffold_11 | CP |
| Dentigerous Arm Depth | otol1 | HiC_scaffold_5 | LL |
| Dentigerous Arm Depth | otulin | HiC_scaffold_11 | CP |
| Dentigerous Arm Depth | oxsr1 | HiC_scaffold_11 | CP |
| Dentigerous Arm Depth | pabpn1 | HiC_scaffold_11 | CP |
| Dentigerous Arm Depth | pan3 | HiC_scaffold_11 | CP |
| Dentigerous Arm Depth | pck2 | HiC_scaffold_11 | CP |
| Dentigerous Arm Depth | pcolce2 | HiC_scaffold_11 | CP |
| Dentigerous Arm Depth | pdcd6 | HiC_scaffold_11 | CP |
| Dentigerous Arm Depth | pde7a | HiC_scaffold_11 | CP |
| Dentigerous Arm Depth | pdia4 | HiC_scaffold_11 | CP |
| Dentigerous Arm Depth | pdk3 | HiC_scaffold_11 | CP |
| Dentigerous Arm Depth | pdpr | HiC_scaffold_5 | LL |
| Dentigerous Arm Depth | pdx1 | HiC_scaffold_11 | CP |
| Dentigerous Arm Depth | pex2 | HiC_scaffold_11 | CP |
| Dentigerous Arm Depth | pgbd2 | HiC_scaffold_11 | CP |
| Dentigerous Arm Depth | phex | HiC_scaffold_11 | CP |
| Dentigerous Arm Depth | phldb2 | HiC_scaffold_11 | CP |
| Dentigerous Arm Depth | pi15a | HiC_scaffold_11 | CP |

| | | | |
|---|---|---|---|
| Dentigerous Arm Depth | pigm | HiC_scaffold_11 | CP |
| Dentigerous Arm Depth | pign | HiC_scaffold_11 | CP |
| Dentigerous Arm Depth | pim2 | HiC_scaffold_11 | CP |
| Dentigerous Arm Depth | pkia | HiC_scaffold_11 | CP |
| Dentigerous Arm Depth | pkp3 | HiC_scaffold_5 | LL |
| Dentigerous Arm Depth | pks15/1 | HiC_scaffold_11 | CP |
| Dentigerous Arm Depth | plcd1 | HiC_scaffold_11 | CP |
| Dentigerous Arm Depth | plch2 | HiC_scaffold_11 | CP |
| Dentigerous Arm Depth | plcxd2 | HiC_scaffold_11 | CP |
| Dentigerous Arm Depth | plod2 | HiC_scaffold_11 | CP |
| Dentigerous Arm Depth | plscr2 | HiC_scaffold_11 | CP |
| Dentigerous Arm Depth | pnoc | HiC_scaffold_5 | LL |
| Dentigerous Arm Depth | pola1 | HiC_scaffold_11 | CP |
| Dentigerous Arm Depth | polr1d | HiC_scaffold_11 | CP |
| Dentigerous Arm Depth | pomp | HiC_scaffold_11 | CP |
| Dentigerous Arm Depth | pop4 | HiC_scaffold_5 | LL |
| Dentigerous Arm Depth | pou6f2 | HiC_scaffold_11 | CP |
| Dentigerous Arm Depth | ppcs | HiC_scaffold_11 | CP |
| Dentigerous Arm Depth | ppp1r16a | HiC_scaffold_11 | CP |
| Dentigerous Arm Depth | ppp1r42 | HiC_scaffold_11 | CP |
| Dentigerous Arm Depth | prdm14 | HiC_scaffold_11 | CP |
| Dentigerous Arm Depth | prdm2 | HiC_scaffold_11 | CP |
| Dentigerous Arm Depth | prex2 | HiC_scaffold_11 | CP |
| Dentigerous Arm Depth | prkdc | HiC_scaffold_11 | CP |
| Dentigerous Arm Depth | prlh | HiC_scaffold_11 | CP |
| Dentigerous Arm Depth | prmt6 | HiC_scaffold_11 | CP |
| Dentigerous Arm Depth | proc | HiC_scaffold_5 | LL |
| Dentigerous Arm Depth | prpf4b | HiC_scaffold_11 | CP |
| Dentigerous Arm Depth | prtfdc1 | HiC_scaffold_11 | CP |
| Dentigerous Arm Depth | psma4 | HiC_scaffold_5 | LL |
| Dentigerous Arm Depth | psme1 | HiC_scaffold_11 | CP |
| Dentigerous Arm Depth | psme2 | HiC_scaffold_11 | CP |
| Dentigerous Arm Depth | ptprn2 | HiC_scaffold_11 | CP |
| Dentigerous Arm Depth | puf60 | HiC_scaffold_11 | CP |
| Dentigerous Arm Depth | qtrt2 | HiC_scaffold_11 | CP |
| Dentigerous Arm Depth | rala | HiC_scaffold_11 | CP |
| Dentigerous Arm Depth | ralyl | HiC_scaffold_11 | CP |
| Dentigerous Arm Depth | ramp3 | HiC_scaffold_11 | CP |
| Dentigerous Arm Depth | rbm33 | HiC_scaffold_11 | CP |
| Dentigerous Arm Depth | rbpms2 | HiC_scaffold_5 | LL |
| Dentigerous Arm Depth | rdh10a | HiC_scaffold_11 | CP |

| | | | |
|---|---|---|---|
| Dentigerous Arm Depth | rdh12 | HiC_scaffold_11 | CP |
| Dentigerous Arm Depth | rec8 | HiC_scaffold_11 | CP |
| Dentigerous Arm Depth | relch | HiC_scaffold_11 | CP |
| Dentigerous Arm Depth | rgs20 | HiC_scaffold_11 | CP |
| Dentigerous Arm Depth | rgs9bp-b | HiC_scaffold_11 | CP |
| Dentigerous Arm Depth | rmdn1 | HiC_scaffold_11 | CP |
| Dentigerous Arm Depth | rnf152 | HiC_scaffold_11 | CP |
| Dentigerous Arm Depth | rnf6 | HiC_scaffold_11 | CP |
| Dentigerous Arm Depth | rnh1 | HiC_scaffold_11 | CP |
| Dentigerous Arm Depth | rnh1 | HiC_scaffold_5 | LL |
| Dentigerous Arm Depth | rnpc3 | HiC_scaffold_11 | CP |
| Dentigerous Arm Depth | rp1 | HiC_scaffold_11 | CP |
| Dentigerous Arm Depth | rpl21 | HiC_scaffold_11 | CP |
| Dentigerous Arm Depth | rpl7 | HiC_scaffold_11 | CP |
| Dentigerous Arm Depth | rps17 | HiC_scaffold_5 | LL |
| Dentigerous Arm Depth | rps20 | HiC_scaffold_11 | CP |
| Dentigerous Arm Depth | rrs1 | HiC_scaffold_11 | CP |
| Dentigerous Arm Depth | rxfp3 | HiC_scaffold_5 | LL |
| Dentigerous Arm Depth | sag | HiC_scaffold_5 | LL |
| Dentigerous Arm Depth | sbspon | HiC_scaffold_11 | CP |
| Dentigerous Arm Depth | scamp2 | HiC_scaffold_5 | LL |
| Dentigerous Arm Depth | scamp5-a | HiC_scaffold_5 | LL |
| Dentigerous Arm Depth | scrib | HiC_scaffold_11 | CP |
| Dentigerous Arm Depth | sec22c | HiC_scaffold_11 | CP |
| Dentigerous Arm Depth | sec61g | HiC_scaffold_11 | CP |
| Dentigerous Arm Depth | sema4b | HiC_scaffold_5 | LL |
| Dentigerous Arm Depth | sema5a | HiC_scaffold_11 | CP |
| Dentigerous Arm Depth | senp8 | HiC_scaffold_5 | LL |
| Dentigerous Arm Depth | serpinb1 | HiC_scaffold_11 | CP |
| Dentigerous Arm Depth | serpinb10 | HiC_scaffold_11 | CP |
| Dentigerous Arm Depth | serpinb1b | HiC_scaffold_11 | CP |
| Dentigerous Arm Depth | serpinb6 | HiC_scaffold_11 | CP |
| Dentigerous Arm Depth | sgk3 | HiC_scaffold_11 | CP |
| Dentigerous Arm Depth | sh3glb1 | HiC_scaffold_11 | CP |
| Dentigerous Arm Depth | sh3kbp1 | HiC_scaffold_11 | CP |
| Dentigerous Arm Depth | shhb | HiC_scaffold_11 | CP |
| Dentigerous Arm Depth | shisa2 | HiC_scaffold_11 | CP |
| Dentigerous Arm Depth | si:ch211-238a12.2 | HiC_scaffold_5 | LL |
| Dentigerous Arm Depth | siglec1 | HiC_scaffold_11 | CP |
| Dentigerous Arm Depth | siglec10 | HiC_scaffold_11 | CP |
| Dentigerous Arm Depth | siglec13 | HiC_scaffold_11 | CP |

| | | | |
|---|---|---|---|
| Dentigerous Arm Depth | siglec14 | HiC_scaffold_11 | CP |
| Dentigerous Arm Depth | siglec9 | HiC_scaffold_11 | CP |
| Dentigerous Arm Depth | ski | HiC_scaffold_11 | CP |
| Dentigerous Arm Depth | skida1 | HiC_scaffold_11 | CP |
| Dentigerous Arm Depth | slc22a13 | HiC_scaffold_11 | CP |
| Dentigerous Arm Depth | slc22a17 | HiC_scaffold_11 | CP |
| Dentigerous Arm Depth | slc35b3 | HiC_scaffold_11 | CP |
| Dentigerous Arm Depth | slc35g2 | HiC_scaffold_11 | CP |
| Dentigerous Arm Depth | slc39a12 | HiC_scaffold_11 | CP |
| Dentigerous Arm Depth | slc4a2 | HiC_scaffold_11 | CP |
| Dentigerous Arm Depth | slc51a | HiC_scaffold_11 | CP |
| Dentigerous Arm Depth | slco5a1 | HiC_scaffold_11 | CP |
| Dentigerous Arm Depth | smarcd3 | HiC_scaffold_11 | CP |
| Dentigerous Arm Depth | smchd1 | HiC_scaffold_11 | CP |
| Dentigerous Arm Depth | snx1 | HiC_scaffold_5 | LL |
| Dentigerous Arm Depth | snx16 | HiC_scaffold_11 | CP |
| Dentigerous Arm Depth | socs6 | HiC_scaffold_11 | CP |
| Dentigerous Arm Depth | sox17a | HiC_scaffold_11 | CP |
| Dentigerous Arm Depth | spag16 | HiC_scaffold_11 | CP |
| Dentigerous Arm Depth | spata13 | HiC_scaffold_11 | CP |
| Dentigerous Arm Depth | spice1 | HiC_scaffold_11 | CP |
| Dentigerous Arm Depth | sppl2a | HiC_scaffold_5 | LL |
| Dentigerous Arm Depth | st14 | HiC_scaffold_11 | CP |
| Dentigerous Arm Depth | stard5 | HiC_scaffold_5 | LL |
| Dentigerous Arm Depth | sun3 | HiC_scaffold_5 | LL |
| Dentigerous Arm Depth | sv2b | HiC_scaffold_5 | LL |
| Dentigerous Arm Depth | tagln3 | HiC_scaffold_11 | CP |
| Dentigerous Arm Depth | tcf24 | HiC_scaffold_11 | CP |
| Dentigerous Arm Depth | tecr | HiC_scaffold_11 | CP |
| Dentigerous Arm Depth | terf1 | HiC_scaffold_11 | CP |
| Dentigerous Arm Depth | tfap2a | HiC_scaffold_11 | CP |
| Dentigerous Arm Depth | tfrc | HiC_scaffold_11 | CP |
| Dentigerous Arm Depth | thap6 | HiC_scaffold_11 | CP |
| Dentigerous Arm Depth | thtpa | HiC_scaffold_11 | CP |
| Dentigerous Arm Depth | tlnrd1 | HiC_scaffold_5 | LL |
| Dentigerous Arm Depth | tm9sf1 | HiC_scaffold_11 | CP |
| Dentigerous Arm Depth | tmem14c | HiC_scaffold_11 | CP |
| Dentigerous Arm Depth | tmem236 | HiC_scaffold_11 | CP |
| Dentigerous Arm Depth | tmem51 | HiC_scaffold_11 | CP |
| Dentigerous Arm Depth | tmem56-b | HiC_scaffold_11 | CP |
| Dentigerous Arm Depth | tmprss7 | HiC_scaffold_11 | CP |

| | | | |
|---|---|---|---|
| Dentigerous Arm Depth | tnk2 | HiC_scaffold_11 | CP |
| Dentigerous Arm Depth | topbp1-a | HiC_scaffold_11 | CP |
| Dentigerous Arm Depth | tph1 | HiC_scaffold_5 | LL |
| Dentigerous Arm Depth | tram1l1 | HiC_scaffold_11 | CP |
| Dentigerous Arm Depth | trim55 | HiC_scaffold_11 | CP |
| Dentigerous Arm Depth | trim69 | HiC_scaffold_5 | LL |
| Dentigerous Arm Depth | trip4 | HiC_scaffold_5 | LL |
| Dentigerous Arm Depth | trp53inp1 | HiC_scaffold_5 | LL |
| Dentigerous Arm Depth | trpa1 | HiC_scaffold_11 | CP |
| Dentigerous Arm Depth | trpc1 | HiC_scaffold_11 | CP |
| Dentigerous Arm Depth | trpm7 | HiC_scaffold_5 | LL |
| Dentigerous Arm Depth | tshz3 | HiC_scaffold_5 | LL |
| Dentigerous Arm Depth | tssc4 | HiC_scaffold_5 | LL |
| Dentigerous Arm Depth | tssk1b | HiC_scaffold_11 | CP |
| Dentigerous Arm Depth | tstd3 | HiC_scaffold_11 | CP |
| Dentigerous Arm Depth | txndc5 | HiC_scaffold_11 | CP |
| Dentigerous Arm Depth | tyms | HiC_scaffold_11 | CP |
| Dentigerous Arm Depth | u2surp | HiC_scaffold_11 | CP |
| Dentigerous Arm Depth | ube2w | HiC_scaffold_11 | CP |
| Dentigerous Arm Depth | ube3c | HiC_scaffold_11 | CP |
| Dentigerous Arm Depth | ubl7 | HiC_scaffold_5 | LL |
| Dentigerous Arm Depth | urad | HiC_scaffold_11 | CP |
| Dentigerous Arm Depth | usf3 | HiC_scaffold_11 | CP |
| Dentigerous Arm Depth | usp12 | HiC_scaffold_11 | CP |
| Dentigerous Arm Depth | utp3 | HiC_scaffold_11 | CP |
| Dentigerous Arm Depth | vav3 | HiC_scaffold_11 | CP |
| Dentigerous Arm Depth | vcpip1 | HiC_scaffold_5 | LL |
| Dentigerous Arm Depth | vil1 | HiC_scaffold_11 | CP |
| Dentigerous Arm Depth | vipr1 | HiC_scaffold_11 | CP |
| Dentigerous Arm Depth | vps35 | HiC_scaffold_5 | LL |
| Dentigerous Arm Depth | vstm2a | HiC_scaffold_11 | CP |
| Dentigerous Arm Depth | wasf3 | HiC_scaffold_11 | CP |
| Dentigerous Arm Depth | wdr19 | HiC_scaffold_11 | CP |
| Dentigerous Arm Depth | wdr37 | HiC_scaffold_11 | CP |
| Dentigerous Arm Depth | wdr60 | HiC_scaffold_11 | CP |
| Dentigerous Arm Depth | wwp1 | HiC_scaffold_11 | CP |
| Dentigerous Arm Depth | xcc-b100_1894 | HiC_scaffold_11 | CP |
| Dentigerous Arm Depth | xkr9 | HiC_scaffold_11 | CP |
| Dentigerous Arm Depth | yes1 | HiC_scaffold_11 | CP |
| Dentigerous Arm Depth | yme1l1 | HiC_scaffold_11 | CP |
| Dentigerous Arm Depth | ythdf2 | HiC_scaffold_11 | CP |

| | | | |
|---|---|---|---|
| Dentigerous Arm Depth | zc2hc1a | HiC_scaffold_11 | CP |
| Dentigerous Arm Depth | zdhhc23 | HiC_scaffold_11 | CP |
| Dentigerous Arm Depth | zfand1 | HiC_scaffold_11 | CP |
| Dentigerous Arm Depth | zfhx4 | HiC_scaffold_11 | CP |
| Dentigerous Arm Depth | zic1 | HiC_scaffold_11 | CP |
| Dentigerous Arm Depth | zkscan5 | HiC_scaffold_5 | LL |
| Dentigerous Arm Depth | znf235 | HiC_scaffold_11 | CP |
| Dentigerous Arm Depth | znf25 | HiC_scaffold_5 | LL |
| Dentigerous Arm Depth | znf45 | HiC_scaffold_5 | LL |
| Dentigerous Arm Depth | znf507 | HiC_scaffold_5 | LL |
| Dentigerous Arm Depth | znf569 | HiC_scaffold_5 | LL |
| Dentigerous Arm Depth | znf609 | HiC_scaffold_5 | LL |
| Dentigerous Arm Depth | znf652 | HiC_scaffold_11 | CP |
| Dentigerous Arm Depth | znf710 | HiC_scaffold_5 | LL |
| Dentigerous Arm Width | abca1 | HiC_scaffold_24 | LL |
| Dentigerous Arm Width | abca4 | HiC_scaffold_24 | LL |
| Dentigerous Arm Width | abca7 | HiC_scaffold_24 | LL |
| Dentigerous Arm Width | abr | HiC_scaffold_24 | LL |
| Dentigerous Arm Width | acadvl | HiC_scaffold_24 | LL |
| Dentigerous Arm Width | acan | HiC_scaffold_24 | LL |
| Dentigerous Arm Width | acbp4 | HiC_scaffold_24 | LL |
| Dentigerous Arm Width | acy3.2 | HiC_scaffold_24 | LL |
| Dentigerous Arm Width | adamtsl1 | HiC_scaffold_24 | LL |
| Dentigerous Arm Width | adcy2 | HiC_scaffold_24 | LL |
| Dentigerous Arm Width | adgra3 | HiC_scaffold_24 | LL |
| Dentigerous Arm Width | adgrl3 | HiC_scaffold_24 | LL |
| Dentigerous Arm Width | agfg1 | HiC_scaffold_24 | LL |
| Dentigerous Arm Width | alpk1 | HiC_scaffold_58 | LL |
| Dentigerous Arm Width | ami | HiC_scaffold_24 | LL |
| Dentigerous Arm Width | arap2 | HiC_scaffold_24 | LL |
| Dentigerous Arm Width | arhgef11 | HiC_scaffold_24 | LL |
| Dentigerous Arm Width | arl2 | HiC_scaffold_24 | LL |
| Dentigerous Arm Width | atp6ap1 | HiC_scaffold_24 | LL |
| Dentigerous Arm Width | b3gat3 | HiC_scaffold_24 | LL |
| Dentigerous Arm Width | bad | HiC_scaffold_24 | LL |
| Dentigerous Arm Width | bank1 | HiC_scaffold_24 | LL |
| Dentigerous Arm Width | bcl6b | HiC_scaffold_24 | LL |
| Dentigerous Arm Width | brms1la | HiC_scaffold_24 | LL |
| Dentigerous Arm Width | btn2a1 | HiC_scaffold_24 | LL |
| Dentigerous Arm Width | btn2a2 | HiC_scaffold_24 | LL |
| Dentigerous Arm Width | c1ql4 | HiC_scaffold_24 | LL |

| | | | |
|---|---|---|---|
| Dentigerous Arm Width | cabp4 | HiC_scaffold_24 | LL |
| Dentigerous Arm Width | capg | HiC_scaffold_24 | LL |
| Dentigerous Arm Width | card6 | HiC_scaffold_24 | LL |
| Dentigerous Arm Width | cbln1 | HiC_scaffold_24 | LL |
| Dentigerous Arm Width | ccdc149b | HiC_scaffold_24 | LL |
| Dentigerous Arm Width | cct7 | HiC_scaffold_24 | LL |
| Dentigerous Arm Width | cd48 | HiC_scaffold_24 | LL |
| Dentigerous Arm Width | cdca9 | HiC_scaffold_24 | LL |
| Dentigerous Arm Width | chordc1 | HiC_scaffold_24 | LL |
| Dentigerous Arm Width | chrnb1 | HiC_scaffold_24 | LL |
| Dentigerous Arm Width | chst12 | HiC_scaffold_24 | LL |
| Dentigerous Arm Width | clcn5 | HiC_scaffold_24 | LL |
| Dentigerous Arm Width | cldn7a | HiC_scaffold_24 | LL |
| Dentigerous Arm Width | cldnd1 | HiC_scaffold_24 | LL |
| Dentigerous Arm Width | clec10a | HiC_scaffold_24 | LL |
| Dentigerous Arm Width | clec12b | HiC_scaffold_24 | LL |
| Dentigerous Arm Width | clec20a | HiC_scaffold_24 | LL |
| Dentigerous Arm Width | cmas | HiC_scaffold_24 | LL |
| Dentigerous Arm Width | cnpy3 | HiC_scaffold_24 | LL |
| Dentigerous Arm Width | coro1b | HiC_scaffold_24 | LL |
| Dentigerous Arm Width | cpras1 | HiC_scaffold_24 | LL |
| Dentigerous Arm Width | cpz | HiC_scaffold_24 | LL |
| Dentigerous Arm Width | ctdnep1a | HiC_scaffold_24 | LL |
| Dentigerous Arm Width | cyld | HiC_scaffold_24 | LL |
| Dentigerous Arm Width | cyp26b1 | HiC_scaffold_24 | LL |
| Dentigerous Arm Width | dctn1 | HiC_scaffold_24 | LL |
| Dentigerous Arm Width | dctn6 | HiC_scaffold_24 | LL |
| Dentigerous Arm Width | ddit4l | HiC_scaffold_24 | LL |
| Dentigerous Arm Width | dennd4c | HiC_scaffold_24 | LL |
| Dentigerous Arm Width | dgkd | HiC_scaffold_24 | LL |
| Dentigerous Arm Width | dmrta1 | HiC_scaffold_24 | LL |
| Dentigerous Arm Width | dnai2 | HiC_scaffold_24 | LL |
| Dentigerous Arm Width | dok1 | HiC_scaffold_24 | LL |
| Dentigerous Arm Width | dok7 | HiC_scaffold_24 | LL |
| Dentigerous Arm Width | dtx4 | HiC_scaffold_24 | LL |
| Dentigerous Arm Width | dysf | HiC_scaffold_24 | LL |
| Dentigerous Arm Width | eif5a | HiC_scaffold_24 | LL |
| Dentigerous Arm Width | elavl2 | HiC_scaffold_24 | LL |
| Dentigerous Arm Width | elp5 | HiC_scaffold_24 | LL |
| Dentigerous Arm Width | emc4 | HiC_scaffold_24 | LL |
| Dentigerous Arm Width | endod1 | HiC_scaffold_24 | LL |

| | | | |
|---|---|---|---|
| Dentigerous Arm Width | epd | HiC_scaffold_58 | LL |
| Dentigerous Arm Width | epd2 | HiC_scaffold_58 | LL |
| Dentigerous Arm Width | epo | HiC_scaffold_24 | LL |
| Dentigerous Arm Width | ern1 | HiC_scaffold_24 | LL |
| Dentigerous Arm Width | etnppl | HiC_scaffold_24 | LL |
| Dentigerous Arm Width | fabp2 | HiC_scaffold_24 | LL |
| Dentigerous Arm Width | fcgr2 | HiC_scaffold_24 | LL |
| Dentigerous Arm Width | fn1 | HiC_scaffold_24 | LL |
| Dentigerous Arm Width | fxr1 | HiC_scaffold_58 | LL |
| Dentigerous Arm Width | g0s2 | HiC_scaffold_24 | LL |
| Dentigerous Arm Width | gab1 | HiC_scaffold_24 | LL |
| Dentigerous Arm Width | gabarap | HiC_scaffold_24 | LL |
| Dentigerous Arm Width | gba3 | HiC_scaffold_24 | LL |
| Dentigerous Arm Width | gdi1 | HiC_scaffold_24 | LL |
| Dentigerous Arm Width | gimap3 | HiC_scaffold_24 | LL |
| Dentigerous Arm Width | gimap4 | HiC_scaffold_24 | LL |
| Dentigerous Arm Width | gimap5 | HiC_scaffold_24 | LL |
| Dentigerous Arm Width | gimap6 | HiC_scaffold_24 | LL |
| Dentigerous Arm Width | gimap7 | HiC_scaffold_24 | LL |
| Dentigerous Arm Width | gimap8 | HiC_scaffold_24 | LL |
| Dentigerous Arm Width | gpha2 | HiC_scaffold_24 | LL |
| Dentigerous Arm Width | gpr12 | HiC_scaffold_24 | LL |
| Dentigerous Arm Width | gpr26 | HiC_scaffold_24 | LL |
| Dentigerous Arm Width | gpr4 | HiC_scaffold_24 | LL |
| Dentigerous Arm Width | gps2 | HiC_scaffold_24 | LL |
| Dentigerous Arm Width | gvin1 | HiC_scaffold_24 | LL |
| Dentigerous Arm Width | haus4 | HiC_scaffold_58 | LL |
| Dentigerous Arm Width | hdlbp | HiC_scaffold_24 | LL |
| Dentigerous Arm Width | hgfac | HiC_scaffold_24 | LL |
| Dentigerous Arm Width | hmx1 | HiC_scaffold_24 | LL |
| Dentigerous Arm Width | hmx2 | HiC_scaffold_24 | LL |
| Dentigerous Arm Width | hnrnpc | HiC_scaffold_24 | LL |
| Dentigerous Arm Width | hspa12b | HiC_scaffold_24 | LL |
| Dentigerous Arm Width | htr2a | HiC_scaffold_24 | LL |
| Dentigerous Arm Width | irs1-b | HiC_scaffold_24 | LL |
| Dentigerous Arm Width | itih6 | HiC_scaffold_24 | LL |
| Dentigerous Arm Width | kcnip4 | HiC_scaffold_24 | LL |
| Dentigerous Arm Width | kdm6b | HiC_scaffold_24 | LL |
| Dentigerous Arm Width | kirrel1 | HiC_scaffold_58 | LL |
| Dentigerous Arm Width | klhl33 | HiC_scaffold_24 | LL |
| Dentigerous Arm Width | lgi2 | HiC_scaffold_24 | LL |

| | | | |
|---|---|---|---|
| Dentigerous Arm Width | lpcat4 | HiC_scaffold_24 | LL |
| Dentigerous Arm Width | lrfn2 | HiC_scaffold_24 | LL |
| Dentigerous Arm Width | ltb4r | HiC_scaffold_24 | LL |
| Dentigerous Arm Width | ltb4r2 | HiC_scaffold_24 | LL |
| Dentigerous Arm Width | lurap1l | HiC_scaffold_24 | LL |
| Dentigerous Arm Width | majin | HiC_scaffold_24 | LL |
| Dentigerous Arm Width | mark2 | HiC_scaffold_24 | LL |
| Dentigerous Arm Width | mb21d2 | HiC_scaffold_24 | LL |
| Dentigerous Arm Width | mpdz | HiC_scaffold_24 | LL |
| Dentigerous Arm Width | mrc1 | HiC_scaffold_24 | LL |
| Dentigerous Arm Width | mrc2 | HiC_scaffold_24 | LL |
| Dentigerous Arm Width | mrpl48 | HiC_scaffold_58 | LL |
| Dentigerous Arm Width | msantd1 | HiC_scaffold_24 | LL |
| Dentigerous Arm Width | msmeg_2408 | HiC_scaffold_24 | LL |
| Dentigerous Arm Width | mus81 | HiC_scaffold_24 | LL |
| Dentigerous Arm Width | myadm | HiC_scaffold_24 | LL |
| Dentigerous Arm Width | myoz2 | HiC_scaffold_24 | LL |
| Dentigerous Arm Width | n4bp1 | HiC_scaffold_24 | LL |
| Dentigerous Arm Width | naa40 | HiC_scaffold_24 | LL |
| Dentigerous Arm Width | nagk | HiC_scaffold_24 | LL |
| Dentigerous Arm Width | ndrg2 | HiC_scaffold_24 | LL |
| Dentigerous Arm Width | ndufs2 | HiC_scaffold_58 | LL |
| Dentigerous Arm Width | nectin4 | HiC_scaffold_58 | LL |
| Dentigerous Arm Width | neurl4 | HiC_scaffold_24 | LL |
| Dentigerous Arm Width | nfib | HiC_scaffold_24 | LL |
| Dentigerous Arm Width | nlgn4x | HiC_scaffold_24 | LL |
| Dentigerous Arm Width | nlrc3 | HiC_scaffold_24 | LL |
| Dentigerous Arm Width | nlrp1 | HiC_scaffold_24 | LL |
| Dentigerous Arm Width | nwd2 | HiC_scaffold_24 | LL |
| Dentigerous Arm Width | obscn | HiC_scaffold_24 | LL |
| Dentigerous Arm Width | oga | HiC_scaffold_24 | LL |
| Dentigerous Arm Width | or131-2 | HiC_scaffold_24 | LL |
| Dentigerous Arm Width | osbp | HiC_scaffold_24 | LL |
| Dentigerous Arm Width | ostc | HiC_scaffold_24 | LL |
| Dentigerous Arm Width | otub1 | HiC_scaffold_58 | LL |
| Dentigerous Arm Width | ovol1 | HiC_scaffold_24 | LL |
| Dentigerous Arm Width | p2ry1 | HiC_scaffold_24 | LL |
| Dentigerous Arm Width | paip2b | HiC_scaffold_24 | LL |
| Dentigerous Arm Width | parp14 | HiC_scaffold_24 | LL |
| Dentigerous Arm Width | parp15 | HiC_scaffold_24 | LL |
| Dentigerous Arm Width | parp9 | HiC_scaffold_24 | LL |

| | | | |
|---|---|---|---|
| Dentigerous Arm Width | pcdh7 | HiC_scaffold_24 | LL |
| Dentigerous Arm Width | pced1a | HiC_scaffold_24 | LL |
| Dentigerous Arm Width | pcolce2 | HiC_scaffold_24 | LL |
| Dentigerous Arm Width | pea15 | HiC_scaffold_58 | LL |
| Dentigerous Arm Width | per1 | HiC_scaffold_24 | LL |
| Dentigerous Arm Width | pfkfb1 | HiC_scaffold_24 | LL |
| Dentigerous Arm Width | phf23b | HiC_scaffold_24 | LL |
| Dentigerous Arm Width | pla2r1 | HiC_scaffold_24 | LL |
| Dentigerous Arm Width | plac8l1 | HiC_scaffold_24 | LL |
| Dentigerous Arm Width | plin2 | HiC_scaffold_24 | LL |
| Dentigerous Arm Width | plscr2 | HiC_scaffold_24 | LL |
| Dentigerous Arm Width | polr2a | HiC_scaffold_24 | LL |
| Dentigerous Arm Width | pop7 | HiC_scaffold_24 | LL |
| Dentigerous Arm Width | ppargc1a | HiC_scaffold_24 | LL |
| Dentigerous Arm Width | ppp1r14b | HiC_scaffold_24 | LL |
| Dentigerous Arm Width | ppp2r5b | HiC_scaffold_24 | LL |
| Dentigerous Arm Width | ppp3ca | HiC_scaffold_24 | LL |
| Dentigerous Arm Width | prox1 | HiC_scaffold_24 | LL |
| Dentigerous Arm Width | prss27 | HiC_scaffold_24 | LL |
| Dentigerous Arm Width | prss8 | HiC_scaffold_24 | LL |
| Dentigerous Arm Width | ptprd | HiC_scaffold_24 | LL |
| Dentigerous Arm Width | rab38 | HiC_scaffold_24 | LL |
| Dentigerous Arm Width | rab39b | HiC_scaffold_24 | LL |
| Dentigerous Arm Width | rasgrp2-b | HiC_scaffold_58 | LL |
| Dentigerous Arm Width | rbm4b | HiC_scaffold_58 | LL |
| Dentigerous Arm Width | rbpms | HiC_scaffold_24 | LL |
| Dentigerous Arm Width | rcor2 | HiC_scaffold_24 | LL |
| Dentigerous Arm Width | ripk4 | HiC_scaffold_24 | LL |
| Dentigerous Arm Width | rnf183 | HiC_scaffold_24 | LL |
| Dentigerous Arm Width | rnf223 | HiC_scaffold_24 | LL |
| Dentigerous Arm Width | rpl34 | HiC_scaffold_24 | LL |
| Dentigerous Arm Width | sec24d | HiC_scaffold_24 | LL |
| Dentigerous Arm Width | sema4f | HiC_scaffold_24 | LL |
| Dentigerous Arm Width | sgcz | HiC_scaffold_24 | LL |
| Dentigerous Arm Width | shbg | HiC_scaffold_24 | LL |
| Dentigerous Arm Width | slamf9 | HiC_scaffold_24 | LL |
| Dentigerous Arm Width | slc12a3 | HiC_scaffold_24 | LL |
| Dentigerous Arm Width | slc12a6 | HiC_scaffold_24 | LL |
| Dentigerous Arm Width | slc14a2 | HiC_scaffold_24 | LL |
| Dentigerous Arm Width | slc16a13 | HiC_scaffold_24 | LL |
| Dentigerous Arm Width | slc2a4 | HiC_scaffold_24 | LL |

| | | | |
|---|---|---|---|
| Dentigerous Arm Width | slc8a1 | HiC_scaffold_24 | LL |
| Dentigerous Arm Width | sned1 | HiC_scaffold_24 | LL |
| Dentigerous Arm Width | snx15 | HiC_scaffold_24 | LL |
| Dentigerous Arm Width | spag17 | HiC_scaffold_24 | LL |
| Dentigerous Arm Width | stk26 | HiC_scaffold_24 | LL |
| Dentigerous Arm Width | supt16h | HiC_scaffold_24 | LL |
| Dentigerous Arm Width | synpo2 | HiC_scaffold_24 | LL |
| Dentigerous Arm Width | syt4 | HiC_scaffold_24 | LL |
| Dentigerous Arm Width | taf8 | HiC_scaffold_24 | LL |
| Dentigerous Arm Width | tdrd7b | HiC_scaffold_24 | LL |
| Dentigerous Arm Width | tgas006m08.1 | HiC_scaffold_24 | LL |
| Dentigerous Arm Width | tkfc | HiC_scaffold_24 | LL |
| Dentigerous Arm Width | tmem151b | HiC_scaffold_24 | LL |
| Dentigerous Arm Width | tmem179b | HiC_scaffold_24 | LL |
| Dentigerous Arm Width | tmem55bb | HiC_scaffold_24 | LL |
| Dentigerous Arm Width | tmem88 | HiC_scaffold_24 | LL |
| Dentigerous Arm Width | tmprss15 | HiC_scaffold_24 | LL |
| Dentigerous Arm Width | tnc | HiC_scaffold_24 | LL |
| Dentigerous Arm Width | tnfsf10 | HiC_scaffold_24 | LL |
| Dentigerous Arm Width | tnk2 | HiC_scaffold_24 | LL |
| Dentigerous Arm Width | tox4-b | HiC_scaffold_24 | LL |
| Dentigerous Arm Width | tp53 | HiC_scaffold_24 | LL |
| Dentigerous Arm Width | trbv2 | HiC_scaffold_58 | LL |
| Dentigerous Arm Width | trim27 | HiC_scaffold_24 | LL |
| Dentigerous Arm Width | trim39 | HiC_scaffold_24 | LL |
| Dentigerous Arm Width | trip6 | HiC_scaffold_24 | LL |
| Dentigerous Arm Width | trmt44 | HiC_scaffold_24 | LL |
| Dentigerous Arm Width | tyrp1 | HiC_scaffold_24 | LL |
| Dentigerous Arm Width | ufsp1 | HiC_scaffold_24 | LL |
| Dentigerous Arm Width | ugt2b20 | HiC_scaffold_24 | LL |
| Dentigerous Arm Width | ugt2c1 | HiC_scaffold_24 | LL |
| Dentigerous Arm Width | urgcp | HiC_scaffold_24 | LL |
| Dentigerous Arm Width | vangl2 | HiC_scaffold_24 | LL |
| Dentigerous Arm Width | vbp1 | HiC_scaffold_24 | LL |
| Dentigerous Arm Width | wasf3 | HiC_scaffold_24 | LL |
| Dentigerous Arm Width | ybx1 | HiC_scaffold_24 | LL |
| Dentigerous Arm Width | zbtb38 | HiC_scaffold_24 | LL |
| Dentigerous Arm Width | zdhhc21 | HiC_scaffold_24 | LL |
| Dentigerous Arm Width | zdhhc3 | HiC_scaffold_24 | LL |
| Dentigerous Arm Width | znf638 | HiC_scaffold_24 | LL |
| Jaw closing In-Lever | a1cf | HiC_scaffold_8 | LL |

| | | | |
|---|---|---|---|
| Jaw closing In-Lever | abcc3 | HiC_scaffold_8 | LL |
| Jaw closing In-Lever | acadsb | HiC_scaffold_8 | LL |
| Jaw closing In-Lever | adam12 | HiC_scaffold_8 | LL |
| Jaw closing In-Lever | adap1 | HiC_scaffold_8 | LL |
| Jaw closing In-Lever | ado | HiC_scaffold_8 | LL |
| Jaw closing In-Lever | amdhd2 | HiC_scaffold_8 | LL |
| Jaw closing In-Lever | antxr1 | HiC_scaffold_8 | LL |
| Jaw closing In-Lever | aqp8 | HiC_scaffold_8 | LL |
| Jaw closing In-Lever | arf1 | HiC_scaffold_8 | LL |
| Jaw closing In-Lever | arhgap17 | HiC_scaffold_8 | LL |
| Jaw closing In-Lever | arhgap24 | HiC_scaffold_8 | LL |
| Jaw closing In-Lever | asb12 | HiC_scaffold_8 | LL |
| Jaw closing In-Lever | atp6v0a1 | HiC_scaffold_8 | LL |
| Jaw closing In-Lever | atpaf2 | HiC_scaffold_8 | LL |
| Jaw closing In-Lever | baiap2l1 | HiC_scaffold_8 | LL |
| Jaw closing In-Lever | bbs1 | HiC_scaffold_8 | LL |
| Jaw closing In-Lever | bccip | HiC_scaffold_8 | LL |
| Jaw closing In-Lever | bms1 | HiC_scaffold_8 | LL |
| Jaw closing In-Lever | bricd5 | HiC_scaffold_8 | LL |
| Jaw closing In-Lever | btbd17 | HiC_scaffold_8 | LL |
| Jaw closing In-Lever | bub3 | HiC_scaffold_8 | LL |
| Jaw closing In-Lever | cacna1g | HiC_scaffold_8 | LL |
| Jaw closing In-Lever | cavin1 | HiC_scaffold_8 | LL |
| Jaw closing In-Lever | cbx7 | HiC_scaffold_8 | LL |
| Jaw closing In-Lever | cd163 | HiC_scaffold_8 | LL |
| Jaw closing In-Lever | cdr2l | HiC_scaffold_8 | LL |
| Jaw closing In-Lever | chadl | HiC_scaffold_8 | LL |
| Jaw closing In-Lever | chrm3 | HiC_scaffold_8 | LL |
| Jaw closing In-Lever | chst15 | HiC_scaffold_8 | LL |
| Jaw closing In-Lever | coe3 | HiC_scaffold_8 | LL |
| Jaw closing In-Lever | col14a1 | HiC_scaffold_8 | LL |
| Jaw closing In-Lever | cox19 | HiC_scaffold_8 | LL |
| Jaw closing In-Lever | cpped1 | HiC_scaffold_8 | LL |
| Jaw closing In-Lever | cpxm2 | HiC_scaffold_8 | LL |
| Jaw closing In-Lever | cxcr6 | HiC_scaffold_8 | LL |
| Jaw closing In-Lever | cybc1 | HiC_scaffold_8 | LL |
| Jaw closing In-Lever | d7ertd443e | HiC_scaffold_8 | LL |
| Jaw closing In-Lever | dhrs7ca | HiC_scaffold_8 | LL |
| Jaw closing In-Lever | dnah9 | HiC_scaffold_8 | LL |
| Jaw closing In-Lever | dpysl2 | HiC_scaffold_8 | LL |
| Jaw closing In-Lever | egr2b | HiC_scaffold_8 | LL |

| | | | |
|---|---|---|---|
| Jaw closing In-Lever | elovl6 | HiC_scaffold_8 | LL |
| Jaw closing In-Lever | endod1 | HiC_scaffold_8 | LL |
| Jaw closing In-Lever | ep300 | HiC_scaffold_8 | LL |
| Jaw closing In-Lever | ercc4 | HiC_scaffold_8 | LL |
| Jaw closing In-Lever | exoc6 | HiC_scaffold_8 | LL |
| Jaw closing In-Lever | fads6 | HiC_scaffold_8 | LL |
| Jaw closing In-Lever | fam13a | HiC_scaffold_8 | LL |
| Jaw closing In-Lever | fam171a2 | HiC_scaffold_8 | LL |
| Jaw closing In-Lever | fam53b | HiC_scaffold_8 | LL |
| Jaw closing In-Lever | fasn | HiC_scaffold_8 | LL |
| Jaw closing In-Lever | fdxr | HiC_scaffold_8 | LL |
| Jaw closing In-Lever | fmnl1 | HiC_scaffold_8 | LL |
| Jaw closing In-Lever | foxj1b | HiC_scaffold_8 | LL |
| Jaw closing In-Lever | foxk2 | HiC_scaffold_8 | LL |
| Jaw closing In-Lever | foxl1 | HiC_scaffold_8 | LL |
| Jaw closing In-Lever | frmpd2 | HiC_scaffold_8 | LL |
| Jaw closing In-Lever | galk1 | HiC_scaffold_8 | LL |
| Jaw closing In-Lever | galr2 | HiC_scaffold_8 | LL |
| Jaw closing In-Lever | gas7 | HiC_scaffold_8 | LL |
| Jaw closing In-Lever | gdf10 | HiC_scaffold_8 | LL |
| Jaw closing In-Lever | get4 | HiC_scaffold_8 | LL |
| Jaw closing In-Lever | gid4 | HiC_scaffold_8 | LL |
| Jaw closing In-Lever | gimap4 | HiC_scaffold_8 | LL |
| Jaw closing In-Lever | glp2r | HiC_scaffold_8 | LL |
| Jaw closing In-Lever | gpr142 | HiC_scaffold_8 | LL |
| Jaw closing In-Lever | gpr26 | HiC_scaffold_8 | LL |
| Jaw closing In-Lever | gprc5c | HiC_scaffold_8 | LL |
| Jaw closing In-Lever | grb10 | HiC_scaffold_8 | LL |
| Jaw closing In-Lever | grid2ip | HiC_scaffold_8 | LL |
| Jaw closing In-Lever | grn | HiC_scaffold_8 | LL |
| Jaw closing In-Lever | gsg1l | HiC_scaffold_8 | LL |
| Jaw closing In-Lever | hba1 | HiC_scaffold_8 | LL |
| Jaw closing In-Lever | hbb1 | HiC_scaffold_8 | LL |
| Jaw closing In-Lever | hexd | HiC_scaffold_8 | LL |
| Jaw closing In-Lever | hhex | HiC_scaffold_8 | LL |
| Jaw closing In-Lever | hid1 | HiC_scaffold_8 | LL |
| Jaw closing In-Lever | hmx3b | HiC_scaffold_8 | LL |
| Jaw closing In-Lever | hpdl | HiC_scaffold_8 | LL |
| Jaw closing In-Lever | hs3st3a1 | HiC_scaffold_8 | LL |
| Jaw closing In-Lever | hs3st3b1 | HiC_scaffold_8 | LL |
| Jaw closing In-Lever | itgb4 | HiC_scaffold_8 | LL |

| | | | |
|---|---|---|---|
| Jaw closing In-Lever | jakmip3 | HiC_scaffold_8 | LL |
| Jaw closing In-Lever | jmjd8 | HiC_scaffold_8 | LL |
| Jaw closing In-Lever | kcnj16 | HiC_scaffold_8 | LL |
| Jaw closing In-Lever | kcnj2 | HiC_scaffold_8 | LL |
| Jaw closing In-Lever | kdelr2 | HiC_scaffold_8 | LL |
| Jaw closing In-Lever | kif20b | HiC_scaffold_8 | LL |
| Jaw closing In-Lever | lcmt1 | HiC_scaffold_8 | LL |
| Jaw closing In-Lever | lect2 | HiC_scaffold_8 | LL |
| Jaw closing In-Lever | lhpp | HiC_scaffold_8 | LL |
| Jaw closing In-Lever | llgl2 | HiC_scaffold_8 | LL |
| Jaw closing In-Lever | lmf1 | HiC_scaffold_8 | LL |
| Jaw closing In-Lever | lrrc45 | HiC_scaffold_8 | LL |
| Jaw closing In-Lever | map2k4 | HiC_scaffold_8 | LL |
| Jaw closing In-Lever | map2k6 | HiC_scaffold_8 | LL |
| Jaw closing In-Lever | map3k14 | HiC_scaffold_8 | LL |
| Jaw closing In-Lever | mapk8b | HiC_scaffold_8 | LL |
| Jaw closing In-Lever | mbtd1 | HiC_scaffold_8 | LL |
| Jaw closing In-Lever | meiob | HiC_scaffold_8 | LL |
| Jaw closing In-Lever | mettl9 | HiC_scaffold_8 | LL |
| Jaw closing In-Lever | mfap4 | HiC_scaffold_8 | LL |
| Jaw closing In-Lever | mlst8 | HiC_scaffold_8 | LL |
| Jaw closing In-Lever | mmp21 | HiC_scaffold_8 | LL |
| Jaw closing In-Lever | mms19 | HiC_scaffold_8 | LL |
| Jaw closing In-Lever | mprip | HiC_scaffold_8 | LL |
| Jaw closing In-Lever | mrpl27 | HiC_scaffold_8 | LL |
| Jaw closing In-Lever | mrpl38 | HiC_scaffold_8 | LL |
| Jaw closing In-Lever | mrtfa | HiC_scaffold_8 | LL |
| Jaw closing In-Lever | mrtfb | HiC_scaffold_8 | LL |
| Jaw closing In-Lever | mtfr1l | HiC_scaffold_8 | LL |
| Jaw closing In-Lever | mtr | HiC_scaffold_8 | LL |
| Jaw closing In-Lever | myh16 | HiC_scaffold_8 | LL |
| Jaw closing In-Lever | myh7 | HiC_scaffold_8 | LL |
| Jaw closing In-Lever | myo15a | HiC_scaffold_8 | LL |
| Jaw closing In-Lever | myo15b | HiC_scaffold_8 | LL |
| Jaw closing In-Lever | myocd | HiC_scaffold_8 | LL |
| Jaw closing In-Lever | narf | HiC_scaffold_8 | LL |
| Jaw closing In-Lever | ndufaf4 | HiC_scaffold_8 | LL |
| Jaw closing In-Lever | nme2 | HiC_scaffold_8 | LL |
| Jaw closing In-Lever | noxo1 | HiC_scaffold_8 | LL |
| Jaw closing In-Lever | nploc4 | HiC_scaffold_8 | LL |
| Jaw closing In-Lever | nptx2 | HiC_scaffold_8 | LL |

| | | | |
|---|---|---|---|
| Jaw closing In-Lever | nrbf2 | HiC_scaffold_8 | LL |
| Jaw closing In-Lever | nt5m | HiC_scaffold_8 | LL |
| Jaw closing In-Lever | oat | HiC_scaffold_8 | LL |
| Jaw closing In-Lever | pgp | HiC_scaffold_8 | LL |
| Jaw closing In-Lever | phf5a | HiC_scaffold_8 | LL |
| Jaw closing In-Lever | phyhipl | HiC_scaffold_8 | LL |
| Jaw closing In-Lever | pim3 | HiC_scaffold_8 | LL |
| Jaw closing In-Lever | plau | HiC_scaffold_8 | LL |
| Jaw closing In-Lever | plcd3a | HiC_scaffold_8 | LL |
| Jaw closing In-Lever | plcxd1 | HiC_scaffold_8 | LL |
| Jaw closing In-Lever | polr3d | HiC_scaffold_8 | LL |
| Jaw closing In-Lever | ppp1r3cb | HiC_scaffold_8 | LL |
| Jaw closing In-Lever | prkg1 | HiC_scaffold_8 | LL |
| Jaw closing In-Lever | prss8 | HiC_scaffold_8 | LL |
| Jaw closing In-Lever | pstk | HiC_scaffold_8 | LL |
| Jaw closing In-Lever | pts | HiC_scaffold_8 | LL |
| Jaw closing In-Lever | rab37 | HiC_scaffold_8 | LL |
| Jaw closing In-Lever | rab3gap1 | HiC_scaffold_8 | LL |
| Jaw closing In-Lever | ramp1 | HiC_scaffold_8 | LL |
| Jaw closing In-Lever | rangap1 | HiC_scaffold_8 | LL |
| Jaw closing In-Lever | rasd1 | HiC_scaffold_8 | LL |
| Jaw closing In-Lever | rbp3 | HiC_scaffold_8 | LL |
| Jaw closing In-Lever | reep3 | HiC_scaffold_8 | LL |
| Jaw closing In-Lever | rhbdf1 | HiC_scaffold_8 | LL |
| Jaw closing In-Lever | sap30bp | HiC_scaffold_8 | LL |
| Jaw closing In-Lever | sbk1 | HiC_scaffold_8 | LL |
| Jaw closing In-Lever | shisa6 | HiC_scaffold_8 | LL |
| Jaw closing In-Lever | shisa9 | HiC_scaffold_8 | LL |
| Jaw closing In-Lever | shisa9a | HiC_scaffold_8 | LL |
| Jaw closing In-Lever | slc16a12b | HiC_scaffold_8 | LL |
| Jaw closing In-Lever | slc2a11 | HiC_scaffold_8 | LL |
| Jaw closing In-Lever | slc9a3r1 | HiC_scaffold_8 | LL |
| Jaw closing In-Lever | snx29 | HiC_scaffold_8 | LL |
| Jaw closing In-Lever | sox8 | HiC_scaffold_8 | LL |
| Jaw closing In-Lever | spag9 | HiC_scaffold_8 | LL |
| Jaw closing In-Lever | srcin1 | HiC_scaffold_8 | LL |
| Jaw closing In-Lever | sstr2 | HiC_scaffold_8 | LL |
| Jaw closing In-Lever | st6galnac2 | HiC_scaffold_8 | LL |
| Jaw closing In-Lever | stat5b | HiC_scaffold_8 | LL |
| Jaw closing In-Lever | sult2b1 | HiC_scaffold_8 | LL |
| Jaw closing In-Lever | tcerg1l | HiC_scaffold_8 | LL |

| | | | |
|---|---|---|---|
| Jaw closing In-Lever | tdrkh | HiC_scaffold_8 | LL |
| Jaw closing In-Lever | tex2 | HiC_scaffold_8 | LL |
| Jaw closing In-Lever | tgas113e22.1 | HiC_scaffold_8 | LL |
| Jaw closing In-Lever | thap10 | HiC_scaffold_8 | LL |
| Jaw closing In-Lever | tmem130 | HiC_scaffold_8 | LL |
| Jaw closing In-Lever | tmem238 | HiC_scaffold_8 | LL |
| Jaw closing In-Lever | tmem94 | HiC_scaffold_8 | LL |
| Jaw closing In-Lever | tmprss5 | HiC_scaffold_8 | LL |
| Jaw closing In-Lever | tnfrsf13b | HiC_scaffold_8 | LL |
| Jaw closing In-Lever | tom1l2 | HiC_scaffold_8 | LL |
| Jaw closing In-Lever | trim16 | HiC_scaffold_8 | LL |
| Jaw closing In-Lever | trim39 | HiC_scaffold_8 | LL |
| Jaw closing In-Lever | trim65 | HiC_scaffold_8 | LL |
| Jaw closing In-Lever | trrap | HiC_scaffold_8 | LL |
| Jaw closing In-Lever | trub1 | HiC_scaffold_8 | LL |
| Jaw closing In-Lever | tuba1c | HiC_scaffold_8 | LL |
| Jaw closing In-Lever | tvp23b | HiC_scaffold_8 | LL |
| Jaw closing In-Lever | ubald1 | HiC_scaffold_8 | LL |
| Jaw closing In-Lever | ubtd1 | HiC_scaffold_8 | LL |
| Jaw closing In-Lever | ugt2c1 | HiC_scaffold_8 | LL |
| Jaw closing In-Lever | unc13d | HiC_scaffold_8 | LL |
| Jaw closing In-Lever | unk | HiC_scaffold_8 | LL |
| Jaw closing In-Lever | uros | HiC_scaffold_8 | LL |
| Jaw closing In-Lever | ush1g | HiC_scaffold_8 | LL |
| Jaw closing In-Lever | usp22 | HiC_scaffold_8 | LL |
| Jaw closing In-Lever | uts2r | HiC_scaffold_8 | LL |
| Jaw closing In-Lever | wbp2 | HiC_scaffold_8 | LL |
| Jaw closing In-Lever | wfikkn2 | HiC_scaffold_8 | LL |
| Jaw closing In-Lever | xrcc6 | HiC_scaffold_8 | LL |
| Jaw closing In-Lever | zc3h7b | HiC_scaffold_8 | LL |
| Jaw closing In-Lever | znf235 | HiC_scaffold_8 | LL |
| Jaw closing In-Lever | znf569 | HiC_scaffold_8 | LL |
| Jaw closing In-Lever | znf84 | HiC_scaffold_8 | LL |
| Jaw closing In-Lever | zranb1 | HiC_scaffold_8 | LL |
| Maxillary Head Protrusion | sep7 | HiC_scaffold_53 | LL |
| Maxillary Head Protrusion | abcb1 | HiC_scaffold_53 | LL |
| Maxillary Head Protrusion | abhd16a | HiC_scaffold_53 | LL |
| Maxillary Head Protrusion | acan | HiC_scaffold_53 | LL |
| Maxillary Head Protrusion | adam22 | HiC_scaffold_53 | LL |
| Maxillary Head Protrusion | adamts16 | HiC_scaffold_53 | LL |
| Maxillary Head Protrusion | adar | HiC_scaffold_53 | LL |

| | | | |
|---|---|---|---|
| Maxillary Head Protrusion | adcy2 | HiC_scaffold_53 | LL |
| Maxillary Head Protrusion | agmo | HiC_scaffold_53 | LL |
| Maxillary Head Protrusion | ago1 | HiC_scaffold_53 | LL |
| Maxillary Head Protrusion | ago3 | HiC_scaffold_53 | LL |
| Maxillary Head Protrusion | aicda | HiC_scaffold_53 | LL |
| Maxillary Head Protrusion | aif1l | HiC_scaffold_53 | LL |
| Maxillary Head Protrusion | alg2 | HiC_scaffold_53 | LL |
| Maxillary Head Protrusion | ankib1 | HiC_scaffold_53 | LL |
| Maxillary Head Protrusion | ankmy2 | HiC_scaffold_53 | LL |
| Maxillary Head Protrusion | ankrd28 | HiC_scaffold_53 | LL |
| Maxillary Head Protrusion | apoa1 | HiC_scaffold_53 | LL |
| Maxillary Head Protrusion | apoa4 | HiC_scaffold_53 | LL |
| Maxillary Head Protrusion | apoeb | HiC_scaffold_53 | LL |
| Maxillary Head Protrusion | apoh | HiC_scaffold_53 | LL |
| Maxillary Head Protrusion | aqp10 | HiC_scaffold_53 | LL |
| Maxillary Head Protrusion | arhgef1 | HiC_scaffold_53 | LL |
| Maxillary Head Protrusion | arid1a | HiC_scaffold_53 | LL |
| Maxillary Head Protrusion | arnt | HiC_scaffold_53 | LL |
| Maxillary Head Protrusion | atg12 | HiC_scaffold_53 | LL |
| Maxillary Head Protrusion | atp1a3 | HiC_scaffold_53 | LL |
| Maxillary Head Protrusion | atp6v1c1a | HiC_scaffold_53 | LL |
| Maxillary Head Protrusion | atp8b2 | HiC_scaffold_53 | LL |
| Maxillary Head Protrusion | atxn1 | HiC_scaffold_53 | LL |
| Maxillary Head Protrusion | azi2 | HiC_scaffold_53 | LL |
| Maxillary Head Protrusion | azin1 | HiC_scaffold_53 | LL |
| Maxillary Head Protrusion | baiap2 | HiC_scaffold_53 | LL |
| Maxillary Head Protrusion | bcam | HiC_scaffold_53 | LL |
| Maxillary Head Protrusion | bckdhb | HiC_scaffold_53 | LL |
| Maxillary Head Protrusion | bcl3 | HiC_scaffold_53 | LL |
| Maxillary Head Protrusion | brd2 | HiC_scaffold_53 | LL |
| Maxillary Head Protrusion | brd9 | HiC_scaffold_53 | LL |
| Maxillary Head Protrusion | btg4 | HiC_scaffold_53 | LL |
| Maxillary Head Protrusion | c1orf232 | HiC_scaffold_53 | LL |
| Maxillary Head Protrusion | c1ra | HiC_scaffold_53 | LL |
| Maxillary Head Protrusion | c3ar1 | HiC_scaffold_53 | LL |
| Maxillary Head Protrusion | c4 | HiC_scaffold_53 | LL |
| Maxillary Head Protrusion | c7orf31 | HiC_scaffold_53 | LL |
| Maxillary Head Protrusion | c8orf76 | HiC_scaffold_53 | LL |
| Maxillary Head Protrusion | c8orf88 | HiC_scaffold_53 | LL |
| Maxillary Head Protrusion | ca14 | HiC_scaffold_53 | LL |
| Maxillary Head Protrusion | cacng6 | HiC_scaffold_53 | LL |

| | | | |
|---|---|---|---|
| Maxillary Head Protrusion | cacng7 | HiC_scaffold_53 | LL |
| Maxillary Head Protrusion | cacng8 | HiC_scaffold_53 | LL |
| Maxillary Head Protrusion | carmil1 | HiC_scaffold_53 | LL |
| Maxillary Head Protrusion | casp2 | HiC_scaffold_53 | LL |
| Maxillary Head Protrusion | ccdc106 | HiC_scaffold_53 | LL |
| Maxillary Head Protrusion | ccr3 | HiC_scaffold_53 | LL |
| Maxillary Head Protrusion | ccr4 | HiC_scaffold_53 | LL |
| Maxillary Head Protrusion | ccr5 | HiC_scaffold_53 | LL |
| Maxillary Head Protrusion | cd22 | HiC_scaffold_53 | LL |
| Maxillary Head Protrusion | cd2ap | HiC_scaffold_53 | LL |
| Maxillary Head Protrusion | cd33 | HiC_scaffold_53 | LL |
| Maxillary Head Protrusion | cd4 | HiC_scaffold_53 | LL |
| Maxillary Head Protrusion | cd40 | HiC_scaffold_53 | LL |
| Maxillary Head Protrusion | cd79a | HiC_scaffold_53 | LL |
| Maxillary Head Protrusion | cdc42 | HiC_scaffold_53 | LL |
| Maxillary Head Protrusion | cdc42ep5 | HiC_scaffold_53 | LL |
| Maxillary Head Protrusion | cdca3 | HiC_scaffold_53 | LL |
| Maxillary Head Protrusion | cdh17 | HiC_scaffold_53 | LL |
| Maxillary Head Protrusion | ceacam1 | HiC_scaffold_53 | LL |
| Maxillary Head Protrusion | ceacam2 | HiC_scaffold_53 | LL |
| Maxillary Head Protrusion | ceacam5 | HiC_scaffold_53 | LL |
| Maxillary Head Protrusion | celf3 | HiC_scaffold_53 | LL |
| Maxillary Head Protrusion | cep72 | HiC_scaffold_53 | LL |
| Maxillary Head Protrusion | cers2 | HiC_scaffold_53 | LL |
| Maxillary Head Protrusion | cgn | HiC_scaffold_53 | LL |
| Maxillary Head Protrusion | chrna7 | HiC_scaffold_53 | LL |
| Maxillary Head Protrusion | chrnb2 | HiC_scaffold_53 | LL |
| Maxillary Head Protrusion | ciart | HiC_scaffold_53 | LL |
| Maxillary Head Protrusion | cic | HiC_scaffold_53 | LL |
| Maxillary Head Protrusion | cited3 | HiC_scaffold_53 | LL |
| Maxillary Head Protrusion | cldn12 | HiC_scaffold_53 | LL |
| Maxillary Head Protrusion | clec3b | HiC_scaffold_53 | LL |
| Maxillary Head Protrusion | clk2 | HiC_scaffold_53 | LL |
| Maxillary Head Protrusion | clptm1l | HiC_scaffold_53 | LL |
| Maxillary Head Protrusion | clstn3 | HiC_scaffold_53 | LL |
| Maxillary Head Protrusion | cmc1 | HiC_scaffold_53 | LL |
| Maxillary Head Protrusion | cmlkr1 | HiC_scaffold_53 | LL |
| Maxillary Head Protrusion | cnfn | HiC_scaffold_53 | LL |
| Maxillary Head Protrusion | cnot3 | HiC_scaffold_53 | LL |
| Maxillary Head Protrusion | cnp-1 | HiC_scaffold_53 | LL |
| Maxillary Head Protrusion | cnr2 | HiC_scaffold_53 | LL |

| | | | |
|---|---|---|---|
| Maxillary Head Protrusion | col14a1 | HiC_scaffold_53 | LL |
| Maxillary Head Protrusion | coq3 | HiC_scaffold_53 | LL |
| Maxillary Head Protrusion | cox6b1 | HiC_scaffold_53 | LL |
| Maxillary Head Protrusion | cpq | HiC_scaffold_53 | LL |
| Maxillary Head Protrusion | cpvl | HiC_scaffold_53 | LL |
| Maxillary Head Protrusion | crabp2 | HiC_scaffold_53 | LL |
| Maxillary Head Protrusion | creb5 | HiC_scaffold_53 | LL |
| Maxillary Head Protrusion | crot | HiC_scaffold_53 | LL |
| Maxillary Head Protrusion | CP | HiC_scaffold_53 | LL |
| Maxillary Head Protrusion | csmd3 | HiC_scaffold_53 | LL |
| Maxillary Head Protrusion | csnk2a1 | HiC_scaffold_53 | LL |
| Maxillary Head Protrusion | csnk2b | HiC_scaffold_53 | LL |
| Maxillary Head Protrusion | cspg5 | HiC_scaffold_53 | LL |
| Maxillary Head Protrusion | cthrc1 | HiC_scaffold_53 | LL |
| Maxillary Head Protrusion | ctrl | HiC_scaffold_53 | LL |
| Maxillary Head Protrusion | cxcr3 | HiC_scaffold_53 | LL |
| Maxillary Head Protrusion | cxcr3.2 | HiC_scaffold_53 | LL |
| Maxillary Head Protrusion | cxcr4-b | HiC_scaffold_53 | LL |
| Maxillary Head Protrusion | cyp21a2 | HiC_scaffold_53 | LL |
| Maxillary Head Protrusion | cyp4b1 | HiC_scaffold_53 | LL |
| Maxillary Head Protrusion | cyth2 | HiC_scaffold_53 | LL |
| Maxillary Head Protrusion | cyth3 | HiC_scaffold_53 | LL |
| Maxillary Head Protrusion | d215 | HiC_scaffold_53 | LL |
| Maxillary Head Protrusion | dbi | HiC_scaffold_53 | LL |
| Maxillary Head Protrusion | dcaf13 | HiC_scaffold_53 | LL |
| Maxillary Head Protrusion | dcbld1 | HiC_scaffold_53 | LL |
| Maxillary Head Protrusion | ddr1 | HiC_scaffold_53 | LL |
| Maxillary Head Protrusion | dedd2 | HiC_scaffold_53 | LL |
| Maxillary Head Protrusion | dennd3 | HiC_scaffold_53 | LL |
| Maxillary Head Protrusion | depdc1b | HiC_scaffold_53 | LL |
| Maxillary Head Protrusion | deptor | HiC_scaffold_53 | LL |
| Maxillary Head Protrusion | derl1 | HiC_scaffold_53 | LL |
| Maxillary Head Protrusion | dgat1 | HiC_scaffold_53 | LL |
| Maxillary Head Protrusion | dgkb | HiC_scaffold_53 | LL |
| Maxillary Head Protrusion | dnah11 | HiC_scaffold_53 | LL |
| Maxillary Head Protrusion | dnali1 | HiC_scaffold_53 | LL |
| Maxillary Head Protrusion | dop1a | HiC_scaffold_53 | LL |
| Maxillary Head Protrusion | dpy19l1 | HiC_scaffold_53 | LL |
| Maxillary Head Protrusion | dpys | HiC_scaffold_53 | LL |
| Maxillary Head Protrusion | dscc1 | HiC_scaffold_53 | LL |
| Maxillary Head Protrusion | e2f3 | HiC_scaffold_53 | LL |

| | | | |
|---|---|---|---|
| Maxillary Head Protrusion | ebag9 | HiC_scaffold_53 | LL |
| Maxillary Head Protrusion | ecm1 | HiC_scaffold_53 | LL |
| Maxillary Head Protrusion | edn1 | HiC_scaffold_53 | LL |
| Maxillary Head Protrusion | efhb | HiC_scaffold_53 | LL |
| Maxillary Head Protrusion | efna1 | HiC_scaffold_53 | LL |
| Maxillary Head Protrusion | efna3 | HiC_scaffold_53 | LL |
| Maxillary Head Protrusion | eif3e | HiC_scaffold_53 | LL |
| Maxillary Head Protrusion | emc2 | HiC_scaffold_53 | LL |
| Maxillary Head Protrusion | emg1 | HiC_scaffold_53 | LL |
| Maxillary Head Protrusion | entpd3 | HiC_scaffold_53 | LL |
| Maxillary Head Protrusion | epb41 | HiC_scaffold_53 | LL |
| Maxillary Head Protrusion | ephb5 | HiC_scaffold_53 | LL |
| Maxillary Head Protrusion | epn1 | HiC_scaffold_53 | LL |
| Maxillary Head Protrusion | erf | HiC_scaffold_53 | LL |
| Maxillary Head Protrusion | erp44 | HiC_scaffold_53 | LL |
| Maxillary Head Protrusion | esrp1 | HiC_scaffold_53 | LL |
| Maxillary Head Protrusion | etfb | HiC_scaffold_53 | LL |
| Maxillary Head Protrusion | ethe1 | HiC_scaffold_53 | LL |
| Maxillary Head Protrusion | etv1 | HiC_scaffold_53 | LL |
| Maxillary Head Protrusion | eva1b | HiC_scaffold_53 | LL |
| Maxillary Head Protrusion | fabp4 | HiC_scaffold_53 | LL |
| Maxillary Head Protrusion | fam110c | HiC_scaffold_53 | LL |
| Maxillary Head Protrusion | fam131b | HiC_scaffold_53 | LL |
| Maxillary Head Protrusion | fam189b | HiC_scaffold_53 | LL |
| Maxillary Head Protrusion | fam83a | HiC_scaffold_53 | LL |
| Maxillary Head Protrusion | fam8a1 | HiC_scaffold_53 | LL |
| Maxillary Head Protrusion | faxc | HiC_scaffold_53 | LL |
| Maxillary Head Protrusion | fbxl4 | HiC_scaffold_53 | LL |
| Maxillary Head Protrusion | fbxw7 | HiC_scaffold_53 | LL |
| Maxillary Head Protrusion | fcgbp | HiC_scaffold_53 | LL |
| Maxillary Head Protrusion | fdps | HiC_scaffold_53 | LL |
| Maxillary Head Protrusion | fez2 | HiC_scaffold_53 | LL |
| Maxillary Head Protrusion | ffar2 | HiC_scaffold_53 | LL |
| Maxillary Head Protrusion | ffar3 | HiC_scaffold_53 | LL |
| Maxillary Head Protrusion | fhl3 | HiC_scaffold_53 | LL |
| Maxillary Head Protrusion | fhod3 | HiC_scaffold_53 | LL |
| Maxillary Head Protrusion | fkbp14 | HiC_scaffold_53 | LL |
| Maxillary Head Protrusion | flcn | HiC_scaffold_53 | LL |
| Maxillary Head Protrusion | flot1 | HiC_scaffold_53 | LL |
| Maxillary Head Protrusion | foxj2 | HiC_scaffold_53 | LL |
| Maxillary Head Protrusion | foxo3 | HiC_scaffold_53 | LL |

| | | | |
|---|---|---|---|
| Maxillary Head Protrusion | frrs1l | HiC_scaffold_53 | LL |
| Maxillary Head Protrusion | fsbp | HiC_scaffold_53 | LL |
| Maxillary Head Protrusion | fxyd1 | HiC_scaffold_53 | LL |
| Maxillary Head Protrusion | fzd6 | HiC_scaffold_53 | LL |
| Maxillary Head Protrusion | galnt1 | HiC_scaffold_53 | LL |
| Maxillary Head Protrusion | galr1 | HiC_scaffold_53 | LL |
| Maxillary Head Protrusion | gapdh | HiC_scaffold_53 | LL |
| Maxillary Head Protrusion | gba | HiC_scaffold_53 | LL |
| Maxillary Head Protrusion | gdf6a | HiC_scaffold_53 | LL |
| Maxillary Head Protrusion | gem | HiC_scaffold_53 | LL |
| Maxillary Head Protrusion | glcci1 | HiC_scaffold_53 | LL |
| Maxillary Head Protrusion | glipr2 | HiC_scaffold_53 | LL |
| Maxillary Head Protrusion | gmeb1 | HiC_scaffold_53 | LL |
| Maxillary Head Protrusion | gnb3 | HiC_scaffold_53 | LL |
| Maxillary Head Protrusion | gnl2 | HiC_scaffold_53 | LL |
| Maxillary Head Protrusion | gpatch3 | HiC_scaffold_53 | LL |
| Maxillary Head Protrusion | gpn2 | HiC_scaffold_53 | LL |
| Maxillary Head Protrusion | gpr20 | HiC_scaffold_53 | LL |
| Maxillary Head Protrusion | gpr42 | HiC_scaffold_53 | LL |
| Maxillary Head Protrusion | grb10 | HiC_scaffold_53 | LL |
| Maxillary Head Protrusion | grik3 | HiC_scaffold_53 | LL |
| Maxillary Head Protrusion | grik5 | HiC_scaffold_53 | LL |
| Maxillary Head Protrusion | grina | HiC_scaffold_53 | LL |
| Maxillary Head Protrusion | grwd1 | HiC_scaffold_53 | LL |
| Maxillary Head Protrusion | gsdme | HiC_scaffold_53 | LL |
| Maxillary Head Protrusion | gsg1l | HiC_scaffold_53 | LL |
| Maxillary Head Protrusion | gsk3a | HiC_scaffold_53 | LL |
| Maxillary Head Protrusion | gstk1 | HiC_scaffold_53 | LL |
| Maxillary Head Protrusion | gtpbp10 | HiC_scaffold_53 | LL |
| Maxillary Head Protrusion | h2-eb1 | HiC_scaffold_53 | LL |
| Maxillary Head Protrusion | hacl1 | HiC_scaffold_53 | LL |
| Maxillary Head Protrusion | hamp | HiC_scaffold_53 | LL |
| Maxillary Head Protrusion | hamp1 | HiC_scaffold_53 | LL |
| Maxillary Head Protrusion | has1 | HiC_scaffold_53 | LL |
| Maxillary Head Protrusion | has2 | HiC_scaffold_53 | LL |
| Maxillary Head Protrusion | hcn4 | HiC_scaffold_53 | LL |
| Maxillary Head Protrusion | hdac9b | HiC_scaffold_53 | LL |
| Maxillary Head Protrusion | hepacam2 | HiC_scaffold_53 | LL |
| Maxillary Head Protrusion | herpud2 | HiC_scaffold_53 | LL |
| Maxillary Head Protrusion | hey1 | HiC_scaffold_53 | LL |
| Maxillary Head Protrusion | hibadh | HiC_scaffold_53 | LL |

| | | | |
|---|---|---|---|
| Maxillary Head Protrusion | hivep3 | HiC_scaffold_53 | LL |
| Maxillary Head Protrusion | hlf | HiC_scaffold_53 | LL |
| Maxillary Head Protrusion | hoxa10b | HiC_scaffold_53 | LL |
| Maxillary Head Protrusion | hoxa11b | HiC_scaffold_53 | LL |
| Maxillary Head Protrusion | hoxa9b | HiC_scaffold_53 | LL |
| Maxillary Head Protrusion | hspa8 | HiC_scaffold_53 | LL |
| Maxillary Head Protrusion | hspb6 | HiC_scaffold_53 | LL |
| Maxillary Head Protrusion | iffo1 | HiC_scaffold_53 | LL |
| Maxillary Head Protrusion | iglon5 | HiC_scaffold_53 | LL |
| Maxillary Head Protrusion | il16 | HiC_scaffold_53 | LL |
| Maxillary Head Protrusion | il20ra | HiC_scaffold_53 | LL |
| Maxillary Head Protrusion | il6r | HiC_scaffold_53 | LL |
| Maxillary Head Protrusion | ing4 | HiC_scaffold_53 | LL |
| Maxillary Head Protrusion | ino80c | HiC_scaffold_53 | LL |
| Maxillary Head Protrusion | invs | HiC_scaffold_53 | LL |
| Maxillary Head Protrusion | iqcg | HiC_scaffold_53 | LL |
| Maxillary Head Protrusion | irgc | HiC_scaffold_53 | LL |
| Maxillary Head Protrusion | irx2 | HiC_scaffold_53 | LL |
| Maxillary Head Protrusion | irx4 | HiC_scaffold_53 | LL |
| Maxillary Head Protrusion | isoc2 | HiC_scaffold_53 | LL |
| Maxillary Head Protrusion | ispd | HiC_scaffold_53 | LL |
| Maxillary Head Protrusion | itga10 | HiC_scaffold_53 | LL |
| Maxillary Head Protrusion | jazf1 | HiC_scaffold_53 | LL |
| Maxillary Head Protrusion | josd2 | HiC_scaffold_53 | LL |
| Maxillary Head Protrusion | kcnj2 | HiC_scaffold_53 | LL |
| Maxillary Head Protrusion | kcnk5 | HiC_scaffold_53 | LL |
| Maxillary Head Protrusion | kdf1 | HiC_scaffold_53 | LL |
| Maxillary Head Protrusion | khdc4 | HiC_scaffold_53 | LL |
| Maxillary Head Protrusion | kirrel1 | HiC_scaffold_53 | LL |
| Maxillary Head Protrusion | klf10 | HiC_scaffold_53 | LL |
| Maxillary Head Protrusion | krtcap2 | HiC_scaffold_53 | LL |
| Maxillary Head Protrusion | lag3 | HiC_scaffold_53 | LL |
| Maxillary Head Protrusion | laptm4b | HiC_scaffold_53 | LL |
| Maxillary Head Protrusion | lars2 | HiC_scaffold_53 | LL |
| Maxillary Head Protrusion | lenep | HiC_scaffold_53 | LL |
| Maxillary Head Protrusion | leng1 | HiC_scaffold_53 | LL |
| Maxillary Head Protrusion | leng8 | HiC_scaffold_53 | LL |
| Maxillary Head Protrusion | leng9 | HiC_scaffold_53 | LL |
| Maxillary Head Protrusion | lim2 | HiC_scaffold_53 | LL |
| Maxillary Head Protrusion | limd1 | HiC_scaffold_53 | LL |
| Maxillary Head Protrusion | lin37 | HiC_scaffold_53 | LL |

| | | | |
|---|---|---|---|
| Maxillary Head Protrusion | lipe | HiC_scaffold_53 | LL |
| Maxillary Head Protrusion | lmtk3 | HiC_scaffold_53 | LL |
| Maxillary Head Protrusion | lpcat1 | HiC_scaffold_53 | LL |
| Maxillary Head Protrusion | lsr | HiC_scaffold_53 | LL |
| Maxillary Head Protrusion | m6pr | HiC_scaffold_53 | LL |
| Maxillary Head Protrusion | macc1 | HiC_scaffold_53 | LL |
| Maxillary Head Protrusion | mag | HiC_scaffold_53 | LL |
| Maxillary Head Protrusion | maip1 | HiC_scaffold_53 | LL |
| Maxillary Head Protrusion | mal2 | HiC_scaffold_53 | LL |
| Maxillary Head Protrusion | malsu1 | HiC_scaffold_53 | LL |
| Maxillary Head Protrusion | mamu-dra | HiC_scaffold_53 | LL |
| Maxillary Head Protrusion | man1c1 | HiC_scaffold_53 | LL |
| Maxillary Head Protrusion | maneal | HiC_scaffold_53 | LL |
| Maxillary Head Protrusion | map7d1 | HiC_scaffold_53 | LL |
| Maxillary Head Protrusion | matn4 | HiC_scaffold_53 | LL |
| Maxillary Head Protrusion | mboat1 | HiC_scaffold_53 | LL |
| Maxillary Head Protrusion | mboat7 | HiC_scaffold_53 | LL |
| Maxillary Head Protrusion | mbp | HiC_scaffold_53 | LL |
| Maxillary Head Protrusion | mcam | HiC_scaffold_53 | LL |
| Maxillary Head Protrusion | mdc1 | HiC_scaffold_53 | LL |
| Maxillary Head Protrusion | me1 | HiC_scaffold_53 | LL |
| Maxillary Head Protrusion | mecr | HiC_scaffold_53 | LL |
| Maxillary Head Protrusion | med18 | HiC_scaffold_53 | LL |
| Maxillary Head Protrusion | megf8 | HiC_scaffold_53 | LL |
| Maxillary Head Protrusion | meox2 | HiC_scaffold_53 | LL |
| Maxillary Head Protrusion | mep1b | HiC_scaffold_53 | LL |
| Maxillary Head Protrusion | mindy3 | HiC_scaffold_53 | LL |
| Maxillary Head Protrusion | mios | HiC_scaffold_53 | LL |
| Maxillary Head Protrusion | mlf2 | HiC_scaffold_53 | LL |
| Maxillary Head Protrusion | mms22l | HiC_scaffold_53 | LL |
| Maxillary Head Protrusion | mpp6 | HiC_scaffold_53 | LL |
| Maxillary Head Protrusion | mpra | HiC_scaffold_53 | LL |
| Maxillary Head Protrusion | mpv17l | HiC_scaffold_53 | LL |
| Maxillary Head Protrusion | mrc1 | HiC_scaffold_53 | LL |
| Maxillary Head Protrusion | mrpl13 | HiC_scaffold_53 | LL |
| Maxillary Head Protrusion | mrpl17 | HiC_scaffold_53 | LL |
| Maxillary Head Protrusion | mrpl51 | HiC_scaffold_53 | LL |
| Maxillary Head Protrusion | mrps15 | HiC_scaffold_53 | LL |
| Maxillary Head Protrusion | mrs2 | HiC_scaffold_53 | LL |
| Maxillary Head Protrusion | mrtfb | HiC_scaffold_53 | LL |
| Maxillary Head Protrusion | msh5 | HiC_scaffold_53 | LL |

| | | | |
|---|---|---|---|
| Maxillary Head Protrusion | mtdh | HiC_scaffold_53 | LL |
| Maxillary Head Protrusion | mtf1 | HiC_scaffold_53 | LL |
| Maxillary Head Protrusion | mtss1 | HiC_scaffold_53 | LL |
| Maxillary Head Protrusion | mycbp | HiC_scaffold_53 | LL |
| Maxillary Head Protrusion | myh10 | HiC_scaffold_53 | LL |
| Maxillary Head Protrusion | mypop | HiC_scaffold_53 | LL |
| Maxillary Head Protrusion | nacad | HiC_scaffold_53 | LL |
| Maxillary Head Protrusion | nat14 | HiC_scaffold_53 | LL |
| Maxillary Head Protrusion | ncapd2 | HiC_scaffold_53 | LL |
| Maxillary Head Protrusion | ndufa3 | HiC_scaffold_53 | LL |
| Maxillary Head Protrusion | ndufv1 | HiC_scaffold_53 | LL |
| Maxillary Head Protrusion | necap1 | HiC_scaffold_53 | LL |
| Maxillary Head Protrusion | nectin1 | HiC_scaffold_53 | LL |
| Maxillary Head Protrusion | nectin2 | HiC_scaffold_53 | LL |
| Maxillary Head Protrusion | nek10 | HiC_scaffold_53 | LL |
| Maxillary Head Protrusion | nfatc1 | HiC_scaffold_53 | LL |
| Maxillary Head Protrusion | ngfr | HiC_scaffold_53 | LL |
| Maxillary Head Protrusion | nhlrc1 | HiC_scaffold_53 | LL |
| Maxillary Head Protrusion | nkd2l | HiC_scaffold_53 | LL |
| Maxillary Head Protrusion | nmt2 | HiC_scaffold_53 | LL |
| Maxillary Head Protrusion | nop2 | HiC_scaffold_53 | LL |
| Maxillary Head Protrusion | notch1 | HiC_scaffold_53 | LL |
| Maxillary Head Protrusion | nov | HiC_scaffold_53 | LL |
| Maxillary Head Protrusion | nphs1 | HiC_scaffold_53 | LL |
| Maxillary Head Protrusion | nptx2 | HiC_scaffold_53 | LL |
| Maxillary Head Protrusion | nr0b2 | HiC_scaffold_53 | LL |
| Maxillary Head Protrusion | nr1d2 | HiC_scaffold_53 | LL |
| Maxillary Head Protrusion | nr4a3 | HiC_scaffold_53 | LL |
| Maxillary Head Protrusion | nrsn1 | HiC_scaffold_53 | LL |
| Maxillary Head Protrusion | nxph1 | HiC_scaffold_53 | LL |
| Maxillary Head Protrusion | oprd1 | HiC_scaffold_53 | LL |
| Maxillary Head Protrusion | osbpl3 | HiC_scaffold_53 | LL |
| Maxillary Head Protrusion | oscp1 | HiC_scaffold_53 | LL |
| Maxillary Head Protrusion | osgin2 | HiC_scaffold_53 | LL |
| Maxillary Head Protrusion | oxr1 | HiC_scaffold_53 | LL |
| Maxillary Head Protrusion | p3h3 | HiC_scaffold_53 | LL |
| Maxillary Head Protrusion | pafah1b3 | HiC_scaffold_53 | LL |
| Maxillary Head Protrusion | pde11a | HiC_scaffold_53 | LL |
| Maxillary Head Protrusion | pdp1 | HiC_scaffold_53 | LL |
| Maxillary Head Protrusion | pex5 | HiC_scaffold_53 | LL |
| Maxillary Head Protrusion | phactr4b | HiC_scaffold_53 | LL |

| | | | |
|---|---|---|---|
| Maxillary Head Protrusion | phc1 | HiC_scaffold_53 | LL |
| Maxillary Head Protrusion | phldb3 | HiC_scaffold_53 | LL |
| Maxillary Head Protrusion | pigv | HiC_scaffold_53 | LL |
| Maxillary Head Protrusion | pim2 | HiC_scaffold_53 | LL |
| Maxillary Head Protrusion | pip5k1a | HiC_scaffold_53 | LL |
| Maxillary Head Protrusion | pitpnc1 | HiC_scaffold_53 | LL |
| Maxillary Head Protrusion | plcl2 | HiC_scaffold_53 | LL |
| Maxillary Head Protrusion | plec | HiC_scaffold_53 | LL |
| Maxillary Head Protrusion | plekhg4b | HiC_scaffold_53 | LL |
| Maxillary Head Protrusion | plekhg6 | HiC_scaffold_53 | LL |
| Maxillary Head Protrusion | pnisr | HiC_scaffold_53 | LL |
| Maxillary Head Protrusion | pon2 | HiC_scaffold_53 | LL |
| Maxillary Head Protrusion | pop1 | HiC_scaffold_53 | LL |
| Maxillary Head Protrusion | pou2f2 | HiC_scaffold_53 | LL |
| Maxillary Head Protrusion | pou3f1 | HiC_scaffold_53 | LL |
| Maxillary Head Protrusion | pou3f2 | HiC_scaffold_53 | LL |
| Maxillary Head Protrusion | ppp1r8 | HiC_scaffold_53 | LL |
| Maxillary Head Protrusion | prdm9 | HiC_scaffold_53 | LL |
| Maxillary Head Protrusion | prkca | HiC_scaffold_53 | LL |
| Maxillary Head Protrusion | proser3 | HiC_scaffold_53 | LL |
| Maxillary Head Protrusion | prpf3 | HiC_scaffold_53 | LL |
| Maxillary Head Protrusion | prpf31 | HiC_scaffold_53 | LL |
| Maxillary Head Protrusion | prss35 | HiC_scaffold_53 | LL |
| Maxillary Head Protrusion | psenen | HiC_scaffold_53 | LL |
| Maxillary Head Protrusion | psmd4 | HiC_scaffold_53 | LL |
| Maxillary Head Protrusion | ptdss1 | HiC_scaffold_53 | LL |
| Maxillary Head Protrusion | ptk2 | HiC_scaffold_53 | LL |
| Maxillary Head Protrusion | ptp4a3 | HiC_scaffold_53 | LL |
| Maxillary Head Protrusion | ptpn3 | HiC_scaffold_53 | LL |
| Maxillary Head Protrusion | ptpn6 | HiC_scaffold_53 | LL |
| Maxillary Head Protrusion | ptpru | HiC_scaffold_53 | LL |
| Maxillary Head Protrusion | ptx2 | HiC_scaffold_53 | LL |
| Maxillary Head Protrusion | pxn1 | HiC_scaffold_53 | LL |
| Maxillary Head Protrusion | rab5a | HiC_scaffold_53 | LL |
| Maxillary Head Protrusion | rabac1 | HiC_scaffold_53 | LL |
| Maxillary Head Protrusion | racgap1 | HiC_scaffold_53 | LL |
| Maxillary Head Protrusion | rad54b | HiC_scaffold_53 | LL |
| Maxillary Head Protrusion | rapgef5 | HiC_scaffold_53 | LL |
| Maxillary Head Protrusion | rarb | HiC_scaffold_53 | LL |
| Maxillary Head Protrusion | rasip1 | HiC_scaffold_53 | LL |
| Maxillary Head Protrusion | rbfox1l | HiC_scaffold_53 | LL |

| | | | |
|---|---|---|---|
| Maxillary Head Protrusion | rbm12b | HiC_scaffold_53 | LL |
| Maxillary Head Protrusion | rbm24 | HiC_scaffold_53 | LL |
| Maxillary Head Protrusion | rbp1 | HiC_scaffold_53 | LL |
| Maxillary Head Protrusion | rcc1 | HiC_scaffold_53 | LL |
| Maxillary Head Protrusion | rec8 | HiC_scaffold_53 | LL |
| Maxillary Head Protrusion | rft2 | HiC_scaffold_53 | LL |
| Maxillary Head Protrusion | rhbdl2 | HiC_scaffold_53 | LL |
| Maxillary Head Protrusion | rims2 | HiC_scaffold_53 | LL |
| Maxillary Head Protrusion | rnf144b | HiC_scaffold_53 | LL |
| Maxillary Head Protrusion | rnf41 | HiC_scaffold_53 | LL |
| Maxillary Head Protrusion | rpa2 | HiC_scaffold_53 | LL |
| Maxillary Head Protrusion | rpp38 | HiC_scaffold_53 | LL |
| Maxillary Head Protrusion | rps19 | HiC_scaffold_53 | LL |
| Maxillary Head Protrusion | rps27l | HiC_scaffold_53 | LL |
| Maxillary Head Protrusion | rps9 | HiC_scaffold_53 | LL |
| Maxillary Head Protrusion | rragc | HiC_scaffold_53 | LL |
| Maxillary Head Protrusion | rsph4a | HiC_scaffold_53 | LL |
| Maxillary Head Protrusion | rspo1 | HiC_scaffold_53 | LL |
| Maxillary Head Protrusion | rt1-b | HiC_scaffold_53 | LL |
| Maxillary Head Protrusion | rundc3b | HiC_scaffold_53 | LL |
| Maxillary Head Protrusion | rusc1 | HiC_scaffold_53 | LL |
| Maxillary Head Protrusion | rxrba | HiC_scaffold_53 | LL |
| Maxillary Head Protrusion | s100a1 | HiC_scaffold_53 | LL |
| Maxillary Head Protrusion | s100a16 | HiC_scaffold_53 | LL |
| Maxillary Head Protrusion | s100a6 | HiC_scaffold_53 | LL |
| Maxillary Head Protrusion | s100g | HiC_scaffold_53 | LL |
| Maxillary Head Protrusion | sall3 | HiC_scaffold_53 | LL |
| Maxillary Head Protrusion | sbk2 | HiC_scaffold_53 | LL |
| Maxillary Head Protrusion | scgn | HiC_scaffold_53 | LL |
| Maxillary Head Protrusion | scrt2 | HiC_scaffold_53 | LL |
| Maxillary Head Protrusion | sdc2-b | HiC_scaffold_53 | LL |
| Maxillary Head Protrusion | serinc1 | HiC_scaffold_53 | LL |
| Maxillary Head Protrusion | setdb1b | HiC_scaffold_53 | LL |
| Maxillary Head Protrusion | shc1 | HiC_scaffold_53 | LL |
| Maxillary Head Protrusion | she | HiC_scaffold_53 | LL |
| Maxillary Head Protrusion | shisa7 | HiC_scaffold_53 | LL |
| Maxillary Head Protrusion | siglec1 | HiC_scaffold_53 | LL |
| Maxillary Head Protrusion | siglec14 | HiC_scaffold_53 | LL |
| Maxillary Head Protrusion | siglec5 | HiC_scaffold_53 | LL |
| Maxillary Head Protrusion | slc25a32 | HiC_scaffold_53 | LL |
| Maxillary Head Protrusion | slc25a40 | HiC_scaffold_53 | LL |

| | | | |
|---|---|---|---|
| Maxillary Head Protrusion | slc2a3 | HiC_scaffold_53 | LL |
| Maxillary Head Protrusion | slc40a1 | HiC_scaffold_53 | LL |
| Maxillary Head Protrusion | slc50a1 | HiC_scaffold_53 | LL |
| Maxillary Head Protrusion | slc6a3 | HiC_scaffold_53 | LL |
| Maxillary Head Protrusion | smarcc1 | HiC_scaffold_53 | LL |
| Maxillary Head Protrusion | smg9 | HiC_scaffold_53 | LL |
| Maxillary Head Protrusion | smp | HiC_scaffold_53 | LL |
| Maxillary Head Protrusion | smpd5 | HiC_scaffold_53 | LL |
| Maxillary Head Protrusion | snip1 | HiC_scaffold_53 | LL |
| Maxillary Head Protrusion | snx27 | HiC_scaffold_53 | LL |
| Maxillary Head Protrusion | sostdc1 | HiC_scaffold_53 | LL |
| Maxillary Head Protrusion | sox11 | HiC_scaffold_53 | LL |
| Maxillary Head Protrusion | sp8b | HiC_scaffold_53 | LL |
| Maxillary Head Protrusion | spaca6 | HiC_scaffold_53 | LL |
| Maxillary Head Protrusion | sphk2 | HiC_scaffold_53 | LL |
| Maxillary Head Protrusion | spire1 | HiC_scaffold_53 | LL |
| Maxillary Head Protrusion | spsb1 | HiC_scaffold_53 | LL |
| Maxillary Head Protrusion | sqle | HiC_scaffold_53 | LL |
| Maxillary Head Protrusion | st14 | HiC_scaffold_53 | LL |
| Maxillary Head Protrusion | steap4 | HiC_scaffold_53 | LL |
| Maxillary Head Protrusion | stk40 | HiC_scaffold_53 | LL |
| Maxillary Head Protrusion | stmn2 | HiC_scaffold_53 | LL |
| Maxillary Head Protrusion | sybu | HiC_scaffold_53 | LL |
| Maxillary Head Protrusion | taf12 | HiC_scaffold_53 | LL |
| Maxillary Head Protrusion | tap1 | HiC_scaffold_53 | LL |
| Maxillary Head Protrusion | tarsl2 | HiC_scaffold_53 | LL |
| Maxillary Head Protrusion | tbc1d15 | HiC_scaffold_53 | LL |
| Maxillary Head Protrusion | tbc1d31 | HiC_scaffold_53 | LL |
| Maxillary Head Protrusion | tbrg4 | HiC_scaffold_53 | LL |
| Maxillary Head Protrusion | tcea3 | HiC_scaffold_53 | LL |
| Maxillary Head Protrusion | tent4a | HiC_scaffold_53 | LL |
| Maxillary Head Protrusion | tent5a | HiC_scaffold_53 | LL |
| Maxillary Head Protrusion | tex10 | HiC_scaffold_53 | LL |
| Maxillary Head Protrusion | tfpt | HiC_scaffold_53 | LL |
| Maxillary Head Protrusion | them4 | HiC_scaffold_53 | LL |
| Maxillary Head Protrusion | themis2 | HiC_scaffold_53 | LL |
| Maxillary Head Protrusion | thsd4 | HiC_scaffold_53 | LL |
| Maxillary Head Protrusion | tlr13 | HiC_scaffold_53 | LL |
| Maxillary Head Protrusion | tmc7 | HiC_scaffold_53 | LL |
| Maxillary Head Protrusion | tmem106b | HiC_scaffold_53 | LL |
| Maxillary Head Protrusion | tmem107 | HiC_scaffold_53 | LL |

| | | | |
|---|---|---|---|
| Maxillary Head Protrusion | tmem158 | HiC_scaffold_53 | LL |
| Maxillary Head Protrusion | tmem222 | HiC_scaffold_53 | LL |
| Maxillary Head Protrusion | tmem238 | HiC_scaffold_53 | LL |
| Maxillary Head Protrusion | tmem244 | HiC_scaffold_53 | LL |
| Maxillary Head Protrusion | tmem245 | HiC_scaffold_53 | LL |
| Maxillary Head Protrusion | tmem65 | HiC_scaffold_53 | LL |
| Maxillary Head Protrusion | tmem67 | HiC_scaffold_53 | LL |
| Maxillary Head Protrusion | tmem74 | HiC_scaffold_53 | LL |
| Maxillary Head Protrusion | tmprss6 | HiC_scaffold_53 | LL |
| Maxillary Head Protrusion | tnc | HiC_scaffold_53 | LL |
| Maxillary Head Protrusion | tnfrsf11b | HiC_scaffold_53 | LL |
| Maxillary Head Protrusion | tnfrsf1a | HiC_scaffold_53 | LL |
| Maxillary Head Protrusion | tnfrsf6b | HiC_scaffold_53 | LL |
| Maxillary Head Protrusion | tpbg | HiC_scaffold_53 | LL |
| Maxillary Head Protrusion | tpi1b | HiC_scaffold_53 | LL |
| Maxillary Head Protrusion | tppp | HiC_scaffold_53 | LL |
| Maxillary Head Protrusion | trhr | HiC_scaffold_53 | LL |
| Maxillary Head Protrusion | trim16 | HiC_scaffold_53 | LL |
| Maxillary Head Protrusion | trim25 | HiC_scaffold_53 | LL |
| Maxillary Head Protrusion | trim46 | HiC_scaffold_53 | LL |
| Maxillary Head Protrusion | trio | HiC_scaffold_53 | LL |
| Maxillary Head Protrusion | trip13 | HiC_scaffold_53 | LL |
| Maxillary Head Protrusion | trp3 | HiC_scaffold_53 | LL |
| Maxillary Head Protrusion | trpv5 | HiC_scaffold_53 | LL |
| Maxillary Head Protrusion | tshz1 | HiC_scaffold_53 | LL |
| Maxillary Head Protrusion | tstd3 | HiC_scaffold_53 | LL |
| Maxillary Head Protrusion | ttk | HiC_scaffold_53 | LL |
| Maxillary Head Protrusion | ttyh1 | HiC_scaffold_53 | LL |
| Maxillary Head Protrusion | tuft1 | HiC_scaffold_53 | LL |
| Maxillary Head Protrusion | twist1 | HiC_scaffold_53 | LL |
| Maxillary Head Protrusion | tyrobp | HiC_scaffold_53 | LL |
| Maxillary Head Protrusion | ube2s | HiC_scaffold_53 | LL |
| Maxillary Head Protrusion | ubxn11 | HiC_scaffold_53 | LL |
| Maxillary Head Protrusion | upp1 | HiC_scaffold_53 | LL |
| Maxillary Head Protrusion | usf2 | HiC_scaffold_53 | LL |
| Maxillary Head Protrusion | utp11 | HiC_scaffold_53 | LL |
| Maxillary Head Protrusion | uts2r | HiC_scaffold_53 | LL |
| Maxillary Head Protrusion | vamp1 | HiC_scaffold_53 | LL |
| Maxillary Head Protrusion | vars | HiC_scaffold_53 | LL |
| Maxillary Head Protrusion | vim | HiC_scaffold_53 | LL |
| Maxillary Head Protrusion | virma | HiC_scaffold_53 | LL |

| | | | |
|---|---|---|---|
| Maxillary Head Protrusion | vmo1 | HiC_scaffold_53 | LL |
| Maxillary Head Protrusion | vsig10l | HiC_scaffold_53 | LL |
| Maxillary Head Protrusion | vwde | HiC_scaffold_53 | LL |
| Maxillary Head Protrusion | wdtc1 | HiC_scaffold_53 | LL |
| Maxillary Head Protrusion | wnt4 | HiC_scaffold_53 | LL |
| Maxillary Head Protrusion | xrcc1 | HiC_scaffold_53 | LL |
| Maxillary Head Protrusion | yqjl | HiC_scaffold_53 | LL |
| Maxillary Head Protrusion | yrdc | HiC_scaffold_53 | LL |
| Maxillary Head Protrusion | zadh2 | HiC_scaffold_53 | LL |
| Maxillary Head Protrusion | zbtb7b | HiC_scaffold_53 | LL |
| Maxillary Head Protrusion | zc3h12a | HiC_scaffold_53 | LL |
| Maxillary Head Protrusion | zdhhc11 | HiC_scaffold_53 | LL |
| Maxillary Head Protrusion | zdhhc18 | HiC_scaffold_53 | LL |
| Maxillary Head Protrusion | zhx2 | HiC_scaffold_53 | LL |
| Maxillary Head Protrusion | znf208 | HiC_scaffold_53 | LL |
| Maxillary Head Protrusion | znf226 | HiC_scaffold_53 | LL |
| Maxillary Head Protrusion | znf236 | HiC_scaffold_53 | LL |
| Maxillary Head Protrusion | znf282 | HiC_scaffold_53 | LL |
| Maxillary Head Protrusion | znf385d | HiC_scaffold_53 | LL |
| Maxillary Head Protrusion | znf436 | HiC_scaffold_53 | LL |
| Maxillary Head Protrusion | znf516 | HiC_scaffold_53 | LL |
| Maxillary Head Protrusion | znf524 | HiC_scaffold_53 | LL |
| Maxillary Head Protrusion | znf574 | HiC_scaffold_53 | LL |
| Maxillary Head Protrusion | znf585b | HiC_scaffold_53 | LL |
| Maxillary Head Protrusion | znf628 | HiC_scaffold_53 | LL |
| Maxillary Head Protrusion | znf687a | HiC_scaffold_53 | LL |
| Maxillary Head Protrusion | znf737 | HiC_scaffold_53 | LL |
| Maxillary Head Protrusion | znf79 | HiC_scaffold_53 | LL |
| Maxillary Head Protrusion | znf865 | HiC_scaffold_53 | LL |
